# Testing the predictions of reinforcement: long-term empirical data from a damselfly mosaic hybrid zone

**DOI:** 10.1101/2023.09.20.537982

**Authors:** Luis Rodrigo Arce-Valdés, Andrea Viviana Ballén-Guapacha, Anais Rivas-Torres, Jesús Ramsés Chávez-Ríos, Maren Wellenreuther, Bengt Hansson, Rosa Ana Sánchez Guillén

## Abstract

Theoretical work suggests that reinforcement can cause the strengthening of prezygotic isolation in sympatry by mitigating the costs of maladaptive hybridization. However, only a handful of studies have tested all predictions of this theory in natural populations. We investigated reinforcement in a mosaic hybrid zone of the damselflies *Ischnura elegans* and *I. graellsii.* Firstly, we tested if the conditions of reinforcement were fulfilled by quantifying whether hybridization was costly, and prezygotic isolation was strengthening in sympatry compared with in allopatry. Secondly, we investigated three specific predictions of reinforcement: rarer female effect, presence of concordant prezygotic and postzygotic isolation asymmetries in sympatry, and greater premating asymmetries associated with weaker postzygotic isolation in sympatry. We found that reinforcement has strengthened mechanical isolation in one cross direction between species in sympatry. Our study details a case of reinforcement and heterospecific gene flow causing opposite effects between reciprocal heterospecific crosses and describes a natural model in which reproductive isolation is built by the simultaneous effects of reinforcement, the lock-and-key model, Bateson-Dobzhansky-Müller incompatibilities and Haldane’s rule.

## Introduction

One central goal of evolutionary biology is to understand the processes that lead to the origin of reproductive isolation (RI) during speciation. Reinforcement is a process that can strengthen reproductive barriers and is one of the most widely discussed mechanisms of speciation (Coyne and Orr 2004; Lukhtanov 2011). This phenomenon, proposed and popularized by Dobzhansky (1937, 1940), describes one way in which natural selection can favor speciation (Noor 1999). Reinforcement acts on formerly allopatric, closely related species that come into secondary contact in *de novo* created regions of sympatry. If individuals show variation in their ability to distinguish conspecifics from heterospecifics, some of them may occasionally try to mate with heterospecifics. Reduced fitness of maladaptive hybrids will cause natural selection to reduce the frequency of alleles that are linked with a diminished heterospecific discrimination ability, thus acting to reduce the costs of hybridization (West-Eberhard 1986). This gradually enhances prezygotic isolation between incipient species by Reproductive Character Displacement (RCD), i.e., it enhances the development of greater phenotypic divergence of reproductive traits in sympatry compared with an allopatry scenario (Howard 1993). Reinforcement acts usually either on barriers acting before (prezygotic-premating barriers) or after mating, but before zygote development (prezygotic-postmating barriers; Coyne 1974; Coyne and Orr 2004; Matute 2010b). Theory suggests that this process is capable of gradually reducing the extent of heterospecific matings in sympatric populations over time, and eventually, that this can lead to the cessation of gene flow between sympatric populations, and ultimately speciation (Dobzhansky 1937).

Historically, reinforcement theory has been viewed as a controversial idea (Coyne and Orr 2004), in the main because empirical evidence has been scarce. For instance, reinforcement predicts stronger prezygotic isolation in heterospecific crosses in sympatry than in allopatry (Coyne and Orr 1989; Howard 1993) and indeed, some evidence in support for this pattern was found in nature (Ehrman 1965; Littlejohn 1965; Ratciliffe and Grant 1983; Noor 1995). However, just as quickly as evidence was documented in support for this prediction, there was also a rise of alternative explanations for this enhanced isolation in sympatry. For example, the Templeton effect, or differential fusion, posits that only species that have already achieved strong isolation in allopatry will remain isolated in sympatry; others will merge into a single taxon upon coming in contact (Paterson 1978; Templeton 1981). Thus, higher prezygotic isolation can be observed in sympatry without invoking any selective force. Additional alternative explanations include ecological character displacement (Otte 1989; Noor 1999; Coyne and Orr 2004) and RCD in response to runaway sexual selection (Day 2000). Since Coyne and Orr’s seminal work on *Drosophila* (Coyne and Orr 1989), advocates of the reinforcement theory responded to some of these criticisms by proposing other predictions that could distinguish reinforcement from alternative processes. Firstly, since hybridization costs are usually higher for females than for males, reinforcement theory predicts higher RCD in females than in males (Coyne and Orr 2004). Secondly, since the rarer (the species with the smaller range) or smaller population size species is more frequently involved in heterospecific matings owing to its low frequency in sympatry, reinforcement theory predicts higher isolation in the reciprocal cross direction including the female of the rarer species than in the one including the female of the common species (rarer female effect; Yukilevich 2012). Thirdly, since hybrids produced from the two reciprocal cross directions usually differ in fitness (Turelli and Moyle 2007), reinforcement theory predicts a quicker strengthening of the premating isolation in the cross direction producing hybrids with lower fitness (Yukilevich 2012). Fourthly, since asymmetrical reinforcement increases premating asymmetries, and gene flow purges Bateson-Dobzhansky-Müller (BDM) incompatibilities in sympatry, reinforcement theory predicts both greater premating asymmetries and weaker postzygotic isolation in sympatry than in allopatry (Turelli et al. 2014). Nowadays, reinforcement has been detected across ubiquitous taxa. This indicates that speciation via reinforcement can be widespread in both vertebrate (Hostert 1997; Vallin et al. 2012; Pfennig and Rice 2014; Baiz et al. 2019; St. John and Fuller 2021) and invertebrate animals (Coyne and Orr 1989; Nosil et al. 2003; Lessios 2007; Souza et al. 2008; Dillon et al. 2011; Porretta and Urbanelli 2012; Mérot et al. 2017; Yukilevich 2021). While research on plants is being developed (Ramsey et al. 2003; Moyle et al. 2004; Silvertown et al. 2005; Hopkins 2013; Pellegrino 2016; Roda et al. 2017), research on fungal species (Turner et al. 2010; Giraud and Gourbière 2012) is lagging behind and not much is known so far. Despite the growing body of empirical evidence in invertebrate and vertebrate species in support of reinforcement as an important evolutionary process, not much is known about how consistently this kind of reinforcement occurs in several contact regions of the same pair of species, or about the factors influencing its evolution.

The damselfly species *Ischnura elegans* and *I. graellsii* (Odonata: Coenagrionidae), which in the early 1900s came into secondary contact in Spain (Fig. 1), are a powerful model system to study the evolution of RI. The expansion of *I. elegans* has resulted in a mottled hybrid region, with two secondary contact zones (Sánchez-Guillén et al. 2011, 2023). Mosaic and mottled hybrid zones, i.e., sympatric areas consisting of patches of alternating populations of each parental species and admixed populations (Rand and Harrison 1989), are ideal natural testbeds to study the evolution of reinforcement, for instance, its repeatability across multiple contact areas within a hybrid zone (Cain et al. 1999; Hoskin and Higgie 2013). This is the case of the north-west Spanish hybrid zone, which is characterized by having introgressed populations of each parental species and hybrid populations in which most individuals display different degrees of introgression, i.e., a unimodal distribution (Sánchez-Guillén et al. 2023). Theory predicts that when sympatric speciation occurs, disruptive selection (such as reinforcement) should convert a unimodal distribution of genotypes to a bimodal one (Kondrashov et al. 1998; Jiggins and Mallet 2000). RI between *I. elegans* and *I. graellsii* in the north-west hybrid zone is incomplete and asymmetric. While isolation is almost complete in crosses of *I. graellsii* males and *I. elegans* females owing to mechanical incompatibilities, hybridization usually occurs in the opposite direction (Monetti et al. 2002; Sánchez-Guillén et al. 2012). The incomplete RI, the frequency distribution of the hybrid classes (Sánchez-Guillén et al. 2023), the colonization and recolonization events, and the exceptional long-distribution data on this system all indicate that this system is a good candidate example to evaluate reinforcement.

**Figure 1.**
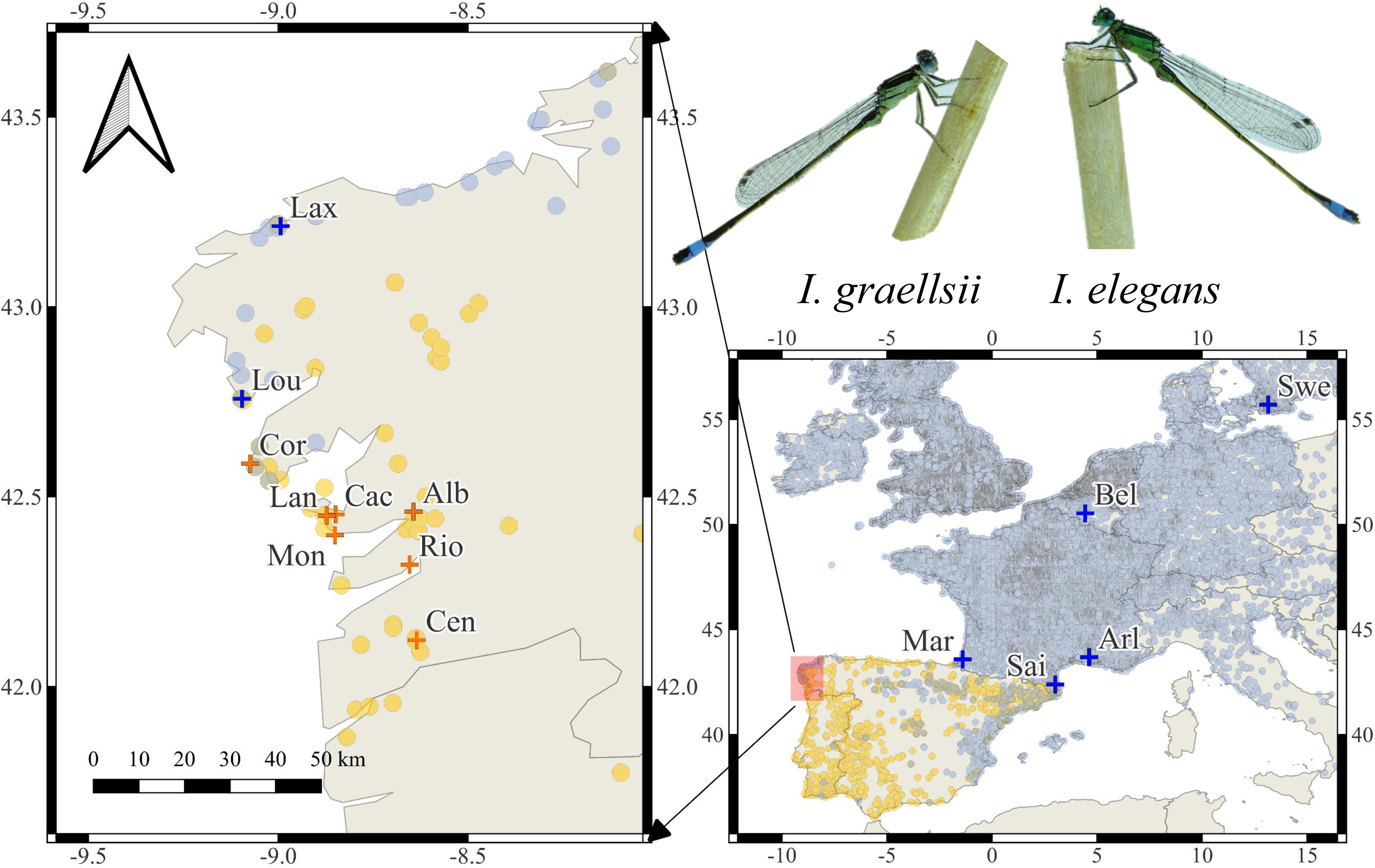
*Ischnura elegans* (blue) and *I. graellsii* (orange) field observations across the north-west Spanish hybrid zone (left) and continental Europe (down-right) from 1758 to 2022 shared by Adolfo Cordero Rivera (*Personal communication*). Crosses show sampled localities. In the top right *I. graellsii* and *I. elegans* males.

In this study, firstly, we evaluated reinforcement in the north-west hybrid zone, and compared the strengths of five reproductive barriers (Fig. 2) in heterospecific crosses of *I. elegans* and *I. graellsii* from the hybrid zone with the strengths of the same five reproductive barriers in heterospecific crosses from allopatric populations. Secondly, we measured the same reproductive barriers in hybrid crosses and backcrosses. We interpreted these measurements as postzygotic barriers and, therefore, as hybridization costs. Reinforcement theory is based on the principle that hybridization costs should be positively correlated with selective pressures directing prezygotic isolation (Ortiz-Barrientos et al. 2009). Thirdly, since theoretical and empirical evidence suggests that the breakdown of reproductive barriers is more likely than reinforcement (Abbott et al. 2013), we used a dataset measuring the same reproductive barriers in other populations from this hybrid zone (Sánchez-Guillén et al. 2012) as a replicate to evaluate the consistency of the hybridization outcomes.

**Figure 2.**
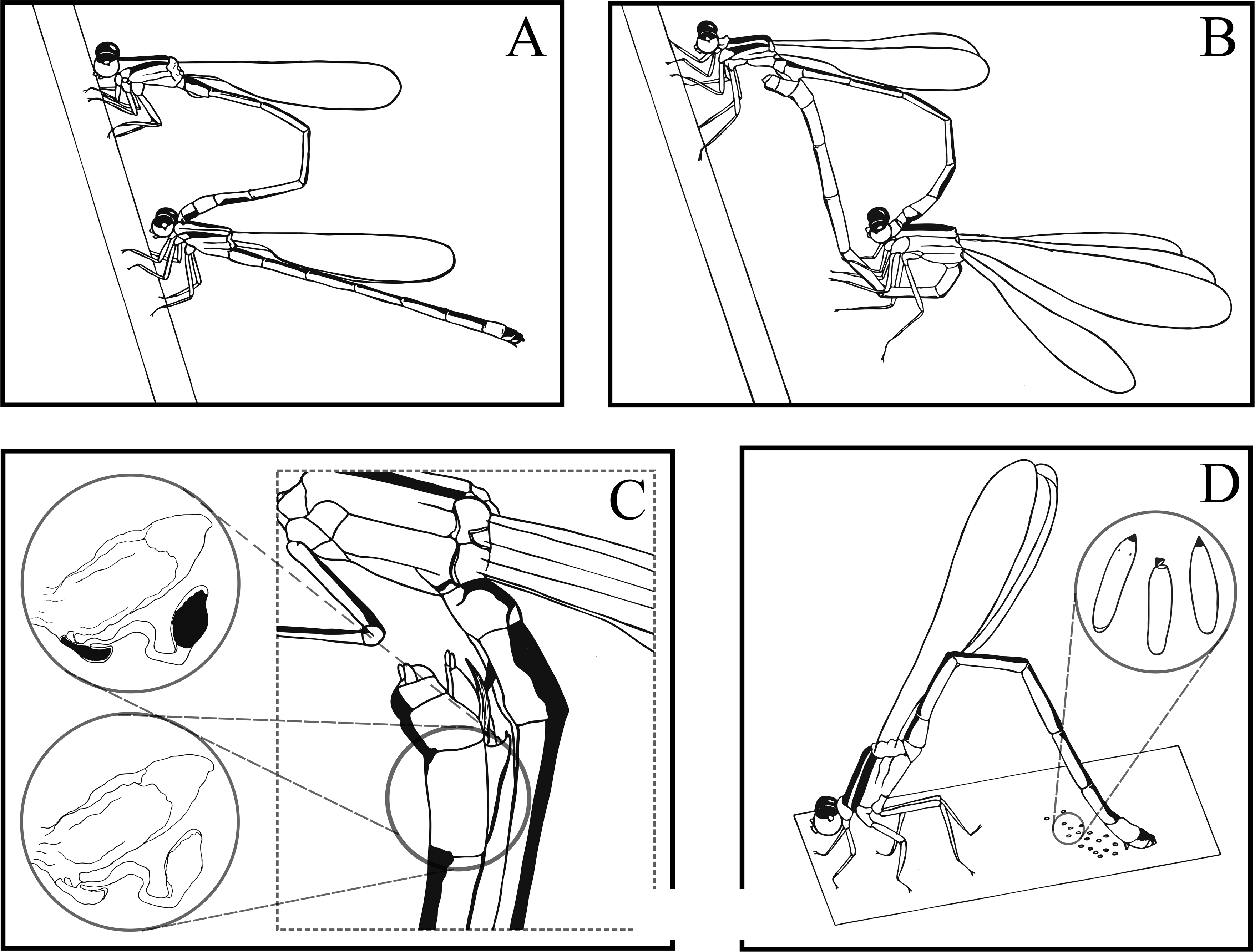
Schematic representation of damselfly reproduction and reproductive barriers measured. A) *Ischnura* damselflies achieving the tandem position (mechanical barrier). B) The female bends her abdomen and primary genitalia come into contact, achieving the mating position (mechanical-tactile barrier). C) Sperm transfer induces the female to oviposit (oviposition barrier). Left-up close-up: sperm is deposited in the female bursa and spermatheca. Left-down close-up: Empty female reproductive tract. D) Female laying eggs. We counted the numbers of eggs in the first three clutches (females were allowed to laid one egg clutch per day, starting from the second day of mating) and averaged them (eggs per clutch index; fecundity barrier). We also measured the ratio of fertile eggs (with visible larvae eyes or opened due to hatching) to the total number of eggs (fertility barrier).

## Materials and Methods

### Sympatry zone (north-west Spanish hybrid zone) description

The north-west Spanish hybrid zone, henceforth called the sympatry zone, is found mainly along the Galician coast (Fig. 1). This sympatry zone is a mosaic hybrid zone in which the frequencies of *I. elegans* and *I. graellsii* vary between populations and thus in their degrees of introgression (Sánchez-Guillén et al. 2023). First records of *I. elegans* in the sympatric zone come from 1980 in Louro (both species), 1987 in Doniños (only *I. elegans*) and 1995 in Foz (both species and hybrids). After that, in between 2000 and 2001, we found *I. elegans* with the occasional presence of *I. graellsii* in Laxe, Carnota and Louro, and between 2001 and 2003 we found both species and hybrids in Cederia and the Corrubedo complex (Table S1). All these populations were, previously to these dates, allopatric for *I. graellsii* (details in Table S1). Currently, these populations mainly consist of introgressed populations of *I. elegans* or introgressed *I. graellsii*, and only one of these populations, Louro, from which *I. elegans* was removed because of salinization of the lagoon in 2010, was after that recolonized in 2013 by both species, and displays different degrees of genetic admixture (introgressed, hybrids, backcrosses, etc.; Sánchez-Guillén et al. 2023). Although the sympatric region has only one hybrid population (Louro, since 2013), we expected to find evidence of reinforcement: firstly, because included populations have recently experienced hybridization, resulting in some cases with the local extinction of *I. graellsii* (Laxe, Doniños, Louro, Foz; Table S1), and in other cases the local extinction of *I. elegans* (e.g., Corrubedo complex; Table S1). The local extinction of one of the hybridizing species has been found in several reinforcement models, when one species outnumbers the other (Servedio and Noor 2003). Our second reason for this expectation was because we detected a signature of RCD of the shape of the *I. elegans* and *I. graellsii* female’s thorax involved in the formation of the copula (Ballén-Guapacha *et al*., *in press*).

### Field samplings

We sampled five pure (allopatric) *I. elegans* populations [one in Sweden (Lund), one in Belgium (De Maten), and three in France (Arles, Saint Cyprien and Marais D’Orx; Table S1 and Fig. 1)], and four *I. graellsii* populations [one pure (allopatric) *I. graellsii* population in Spain (Riomaior) and three *I. graellsii* populations in the Lanzada complex (Lanzada, Montalvo and Cachadas) as localities with putative influence of *I. elegans* owing to its geographic position between the sympatric localities of the Corrubedo complex in the north and the *I. graellsii* allopatric localities in the south (Fig. 1)]. From the north-west Spanish hybrid zone we sampled one *I. elegans* population from the sympatric region (Laxe). Additionally, to evaluate the consistency of the hybridization outcomes, we included (in our data-set) data from the north-west Spanish hybrid zone published in a previous study (Table 1; Sánchez-Guillén et al. 2012), so that, we added to our data-set: two pure (allopatric) *I. graellsii* populations from Spain (Alba, and Centeans), and two populations from the north-west Spanish hybrid zone: one *I. graellsii* population from the Corrubedo complex (Corrubedo, Xuño and Vilar), and two *I. elegans* populations (Laxe and Louro; Table 1; Sánchez-Guillén et al. 2012). We categorized crosses involving these localities as either allopatric or sympatric according to the population of origin of the *I. elegans* individuals with which they were crossed. Finally, several crosses lacked measurements of some of the reproductive barriers we measured (Table 1). These crosses as well as those with a sample size of less than three during the mechanical barrier estimation were excluded from the cumulative RI estimates (Table 1).

**Table 1.**
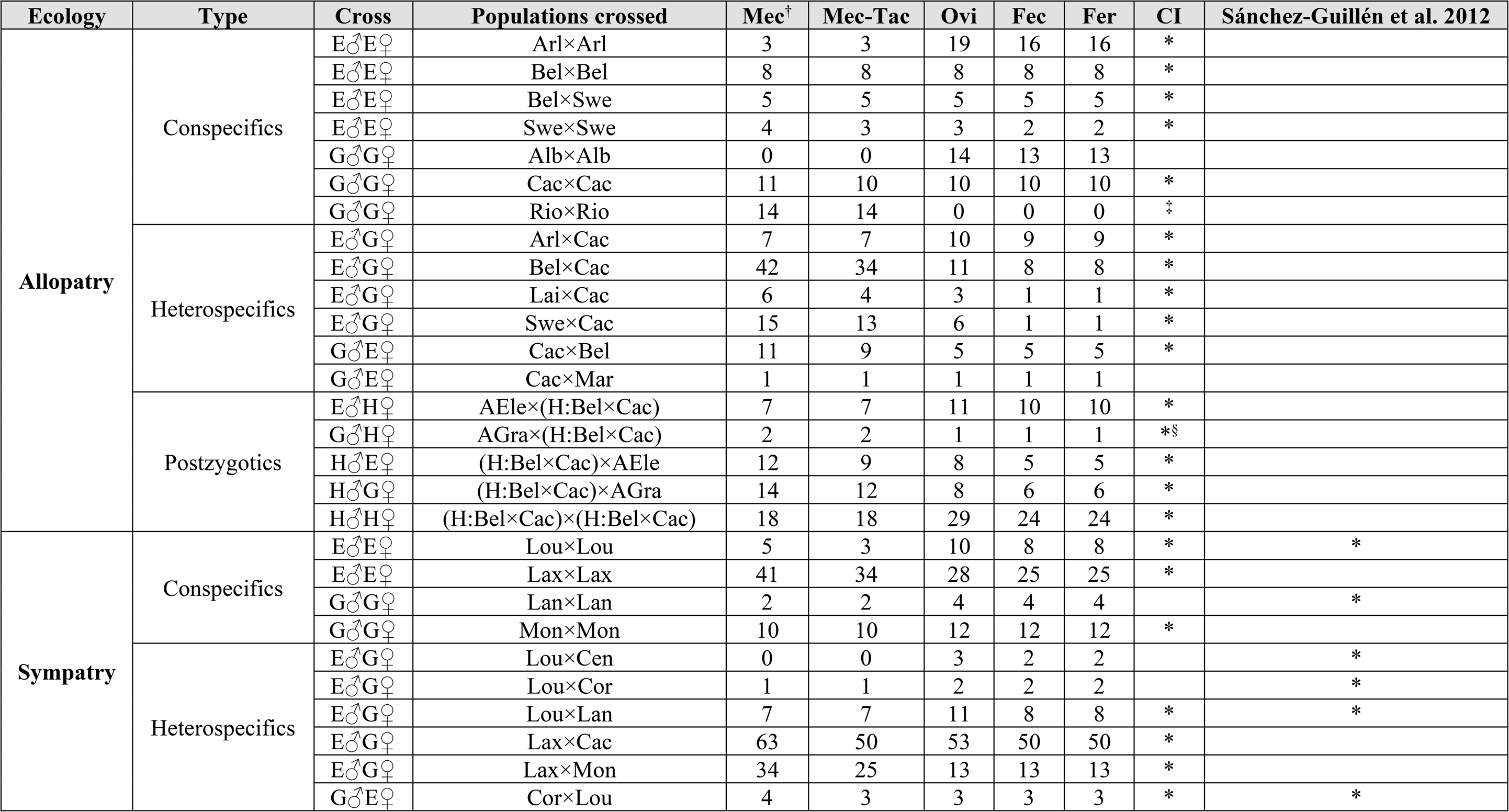
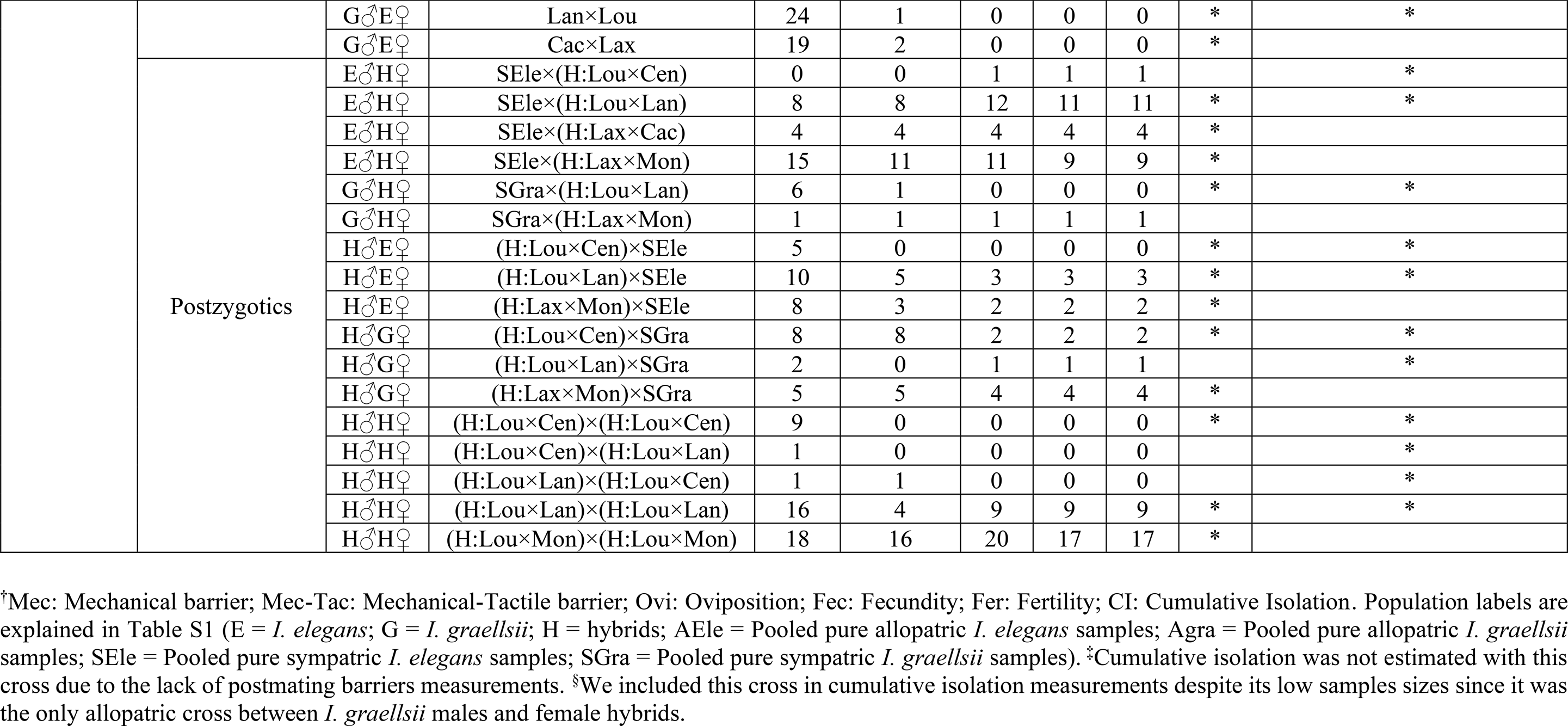
Sample sizes per reproductive barrier measured for each population cross pair. Although all data was used for the absolute isolation estimates and GLM modeling only crosses with a sample size equal or higher than 3 during the mechanical barrier were used for cumulative isolation (CI) estimates (*). The last column shows data reanalyzed from Sánchez-Guillén et al. (2012).

### Rearing in the laboratory and mating trials

Last-instar larvae and tenerals were maintained in the laboratory, until they reached sexual maturity, with the conditions described by Van Gossum et al. (2003). Males and females were kept separated in 50 x 50 x 50 cm wooden insectaries (Van Gossum et al. 2003). During mating trials sexually mature males and females were placed in additional wooden insectaries for observations. We repeated the methods implemented by Sánchez-Guillén et al. (2012). In short, choice trials were made by placing multiple sexually mature male and female damselflies of both species in contact during the hours in which they are most reproductively active (i.e., from 9:00 to 12:00 for *I. elegans* and from 12:00 to 17:00 for *I. graellsii*; thus, observations usually took place between 9:00 and 17:00). The numbers of males and females per insectary were determined by the availability of sexually mature individuals per day. We did not consider mate preference as a reproductive barrier because of the high variability in the frequencies of the species during the experiments. Random individuals per sex were placed in each insectary. All males and non-mated females were placed in daily mating trials until females mated or they died. Couples which successfully mated were isolated until sexual intercourse finished. Then, mated females were isolated and we provided them with the conditions to oviposit (Van Gossum et al. 2003; Sánchez-Guillén et al. 2012). Larvae were reared up to adulthood following standardized protocols (Van Gossum et al. 2003; Sánchez-Guillén et al. 2012), and mating trials were repeated in the following generations.

### Reproduction in *Ischnura* and reproductive barriers

In damselflies the “tandem position” is achieved when the male successfully grasps the female (by her prothorax) using his caudal appendages (Corbet 1999). Copulation begins when the female bends her abdomen and mating organs (genitals) come in contact. This position is usually referred to as “wheel position” (Cordero 1989). Once copulation is achieved, the male first removes sperm from the female’s bursa and spermatheca from previous matings and, after that, inseminates the female. After copulation the female lays eggs until the sperm is finished or she mates again (Fig. 2).

We measured five sequential reproductive barriers: two premating barriers that prevent the tandem (mechanical barrier) and wheel (mechanical-tactile barrier) positions and three postmating barriers that prevent or reduce oviposition, fecundity, and fertility (Fig. 2; Table 2; Text S1). We used each male-female couple or mated female as units of observations for premating and postmating barriers respectively (Table 1). To prevent pseudo-replicates, we avoided the use of several observations from the same male-female pair (Text S1).

**Table 2.** Summary of absolute reproductive isolation formulas per barrier (fitness component). We used the formula proposed by Sobel and Chen (2014): 𝑅𝐼=1− 𝑜𝑏𝑠𝑒𝑟𝑣𝑒𝑑 ℎ𝑦𝑏𝑟𝑖𝑑𝑖𝑧𝑎𝑡𝑖𝑜𝑛𝑒𝑥𝑝𝑒𝑐𝑡𝑒𝑑 ℎ𝑦𝑏𝑟𝑖𝑑𝑖𝑧𝑎𝑡𝑖𝑜𝑛 which represents the proportional decrease of hybridization relative to the null expectation.

| Fitness component | Formula | Isolation range | Estimate |
| --- | --- | --- | --- |
| **Premating: estimated using as replicates male-female interacting couples** |  |  |  |
| I. Mechanical | $RI = 1 - \frac{\text{observed hybridization (number of succesful tandems)}}{\text{expected hybridization (number of tandem attempts)}}$ | 0–1 | Incompatibility between secondary genitalia to form the **tandem position** |
| II. Mechanical-Tactile | $RI = 1 - \frac{\text{observed hybridization (number of succesful copulations)}}{\text{expected hybridization (number of succesful tandems)}}$ | 0–1 | Male fails to stimulate the female to form the **wheel position** or primary genitalia are incompatible |
| **Postmating: estimated using as replicates isolated mated females, i.e., which successfully formed copulation positions with a single male** |  |  |  |
| I. Oviposition | $RI = 1 - \frac{\text{observed hybridization (number of mated females that laid eggs)}}{\text{expected hybridizaton (number of total mated females)}}$ | 0–1 | Sperm fails to stimulate females' **oviposition** |
| II. Fecundity | $RI = 1 - \frac{\text{observed hybridization } (2 * \frac{\sum_{i=1}^n \text{eggs per clutch index}}{n})}{\text{expected hybridization (maximum eggs per clutch Sp1 + maximum eggs per clutch Sp2)}}$ | 0–1 | Sperm reduces rate of females' oviposition (**fecundity**) |
| III. Fertility | $RI = 1 - \frac{\sum_{i=1}^n \frac{\text{observed hybridization (number of fertile eggs)}}{\text{expected hybridization (total laid eggs)}}}{n}$ | 0–1 | Poor transfer or sperm storage, inability of gametes in foreign reproductive tract, poor movement or cross-attraction, or failure of fertilization when gametes contact each other. |

In allopatric crosses, all five reproductive barriers were measured across two generations. F_0_ consisted of conspecific crosses of *I. elegans,* conspecific crosses of *I. graellsii*, and heterospecific crosses of *I. elegans* males and *I. graellsii* females, and *vice versa;* and F_1_ consisted of backcrosses between both species’ males and females with F_1_ hybrids from the opposite sex and crosses between F_1_-hybrids. In sympatric crosses, we were able to additionally measure hybrid crosses and backcrosses in second generation hybrids (F_2_); however, to increase our sample sizes of postzygotic barriers we pooled data from the F_1_ and F_2_ generations. Each barrier was estimated using two values: i) An absolute value that goes from 0 to 1, in which 0 means reproductive barrier absence (complete gene flow) and 1 means complete isolation (gene flow absence); and ii) a relative contribution factor to the total cumulative isolation. See table 1 for the complete list of crosses categorized between the allopatric and sympatric ecologies.

### Absolute and relative strength of the reproductive barriers

Strength of the reproductive barriers in heterospecific and hybrid crosses is frequently estimated using conspecific crosses of one or both parental species as controls (Sánchez-Guillén et al. 2012; Barnard et al. 2017; St. John and Fuller 2021). These controls help measure the mating preference between a conspecific and a heterospecific cross (Sobel and Chen 2014) and are made employing indices such as the Stalker’s Index (Stalker 1942). However, since our main interest was to compare the probability of gene flow between *I. elegans* and *I. graellsii* from allopatry versus from the sympatry zone, we used the formula proposed by Sobel and Chen (2014):

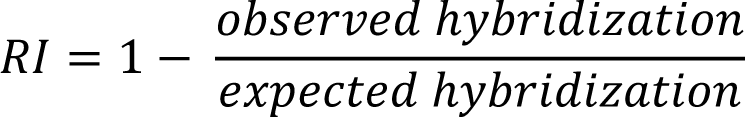

which represents the proportional decrease of hybridization relative to the null expectation (random mating; Table 2). The advantage of this formula for our purpose, which is to compare RI between allopatry and sympatry, is that it can be used to calculate average values and variances when replicated measurements of RI are available. Thus, confidence intervals can also be calculated, and used to calculate the potential range of average reproductive isolation (see Sobel and Chen 2014 for further details). A detailed description on our estimations of each of the five reproductive barriers can be found in the Supplementary Text S1.

To estimate the contribution of each barrier to the total cumulative isolation in sequential stages of reproduction, i.e., its relative contribution, we employed the multiplicative function of individual components developed by Coyne and Orr (1989, 1997) and later modified by Ramsey et al. (2003) to include any number of reproductive barriers (Sobel and Chen 2014). We estimated the cumulative contribution (CC) of a component to the RI at a stage n with the following formula:

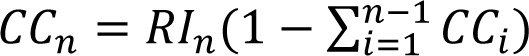

### GLM modeling

To evaluate the effects of the different types of crosses and the “ecology” (sympatry *vs* allopatry) on RI we modeled and compared generalized linear models (GLMs) for each reproductive barrier. For prezygotic barriers (F_0_ generation crosses) we measured the influence of population origin by categorizing them between intrapopulation and interpopulation crosses to create a new variable that we called “geography”. Then, we modeled GLMs of each reproductive barrier isolation as a function of all possible combinations of the types of crosses, the “ecology”, the “geography” and the interaction between the types of crosses and the “ecology” variables. GLMs were modeled using the *glm()* function in R 4.3.0 (R Core Team 2023) and compared using the AICc value with the *dredge()* function of the MuMIN 1.47.5 library (Barton 2009). We modeled the mechanical (successful tandem = 1 *vs* unsuccessful tandem = 0), mechanical-tactile (successful mating = 1 *vs* unsuccessful mating = 0), oviposition (mated female that laid eggs = 1 *vs* mated female that did not laid eggs = 0) and fertility (fertile egg = 1 *vs* unfertile egg = 0) barriers using the binomial distribution, and the fecundity barrier (eggs per clutch index) using the Poisson distribution. We selected as the most probable model per reproductive barrier the one with the lowest AICc score. Pairwise statistical comparisons for the types of crosses and the interaction between the types of crosses and the “ecology” variables were made through post hoc GLMs if these variables were included in the most probable model. This procedure was also applied to postzygotic barriers (F_1_ and F_2_ generation crosses), with the single difference that we did not include the “geography” variable. This variable was excluded because second and third generation crosses highly increased the number of possible combinations of geographical origins of the ancestors of the crossed samples (E. g. crosses between samples whose parents are from the same population, *vs* crosses between samples product of intrapopulation crosses but whose parents come from different populations, *vs* crosses between a sample from an intrapopulation cross and a sample from an interpopulation cross, etc.). Statistical significance tests were used to assess five theoretical predictions of reinforcement (Table 3). See the Supplementary Text S2 for details.

**Table 3.** Summary of reinforcement theoretical predictions tested in *Ischnura elegans* and *I. graellsii*.

| Predictions | Expected patterns | Observed patterns<sup>†</sup> |  |  | Result |
| --- | --- | --- | --- | --- | --- |
|  |  | Sympatry |  | Allopatry |  |
| Strengthening of prezygotic barriers in sympatry (Dobzhansky 1937, 1940) | Stronger **prezygotic** isolation in sympatry than in allopatry | Mechanical RI G♂E♀ (Cac×Lax) | > | Mechanical RI G♂E♀ (Cac×Bel & Cac×Mar) | ✓ |
|  |  | Mechanical RI G♂E♀ (Cor×Lou) | ≈ |  | X |
|  |  | Mechanical RI G♂E♀ (Lan×Lou) | > |  | ✓ |
| Rarer female effect (Yukilevich 2012) | Stronger **prezygotic** isolation in sympatry, but not in allopatry, in the cross-involving females of the rarer species<sup>‡</sup> | This prediction could not be tested as allopatric samples presented strong reproductive isolation asymmetries, thus, confounding the effects population frequencies could have had on reinforcement selective pressures. |  |  | NA |
| Concordant prezygotic and postzygotic isolation asymmetries (Yukilevich 2012) | The asymmetry, in the strength of RI between reciprocal crosses, has the same direction in prezygotic and postzygotic barriers in sympatry (but not in allopatry). | This prediction could not be tested in our data because RI was complete or almost complete (100–94.7%) in one reciprocal cross direction in both allopatry (between *I. elegans* males and *I. graellsii* females) and sympatry (between *I. graellsii* males and *I. elegans* females). This made impossible the comparison between reciprocal crosses either in allopatry or in sympatry. |  |  | NA |
| Greater premating asymmetries and weaker postzygotic isolation in sympatry than in allopatry (Turelli et al. 2014) | Species pairs with asymmetric postzygotic isolation have: i) higher premating asymmetries and ii) weaker postzygotic isolation in sympatry than in allopatry. | i) Mechanical asymmetry (All sympatric data) | > | Mechanical asymmetry (All allopatric data) | ✓ |
|  |  | i) Mechanical asymmetry (CorvsLou) | ≈ | Mechanical asymmetry (BelvsCac) | X |
|  |  | i) Mechanical asymmetry (LanvsLou) | > |  | ✓ |
|  |  | i) Mechanical asymmetry (CacvsLax) | > |  | ✓ |
|  |  | ii) Postzygotic RI E♂H♀ | < | Postzygotic RI E♂H♀ | ?<sup>‡</sup> |
|  |  | ii) Postzygotic RI G♂H♀ | > | Postzygotic RI G♂H♀ | ?<sup>‡</sup> |
|  |  | ii) Postzygotic RI H♂H♀ | < | Postzygotic RI H♂H♀ | ?<sup>‡</sup> |
|  |  | ii) Postzygotic RI H♂E♀ | < | Postzygotic RI H♂E♀ | ?<sup>‡</sup> |
|  |  | ii) Postzygotic RI H♂G♀ | < | Postzygotic RI H♂G♀ | ?<sup>‡</sup> |

<sup>†</sup>E: *I. elegans*; G: *I. graellsii*; H: Hybrid. <sup>‡</sup>Inconclusive, since we could not rear up hybrids from both directions either from allopatry or from sympatry; thus we cannot distinguish if the weaker postzygotic isolation in sympatry was due to purging of BDM incompatibilities via gene flow as predicted by reinforcement, or because species inherit BDM incompatibilities asymmetrically.

## Results

### Rearing experiments

Reproductive barriers were measured considering each male and female pair (premating) and mated female (postmating) as units of observation respectively. While allopatric reproductive barrier measurements were made with between 125 and 180 units of observation per barrier, sympatric reproductive barriers estimations included between 191 and 327 units of observation per barrier (Table 1). While in allopatric crosses reproductive barriers were measured in between one and four pairs of populations in sympatric crosses reproductive barriers were measured in between two and five pairs of populations (Table 1).

### Conspecific crosses

Conspecific crosses behaved similarly between allopatry and sympatry, although *I. elegans* crosses were more successful (i.e., with lower isolation) between allopatric populations than sympatric populations (Fig. 3). In all cases, reproductive success between conspecific crosses was precluded by the cumulative action of all reproductive barriers (Fig. 4). In conspecific *I. graellsii* crosses, reproductive success was largely precluded by low fecundity and fertility, as premating barriers were mostly absent in both allopatric and sympatric crosses (Fig. 4). Overall, reproductive success was similar or slightly higher in conspecific crosses than in heterospecific and hybrid crosses (Fig. 3).

**Figure 3.**
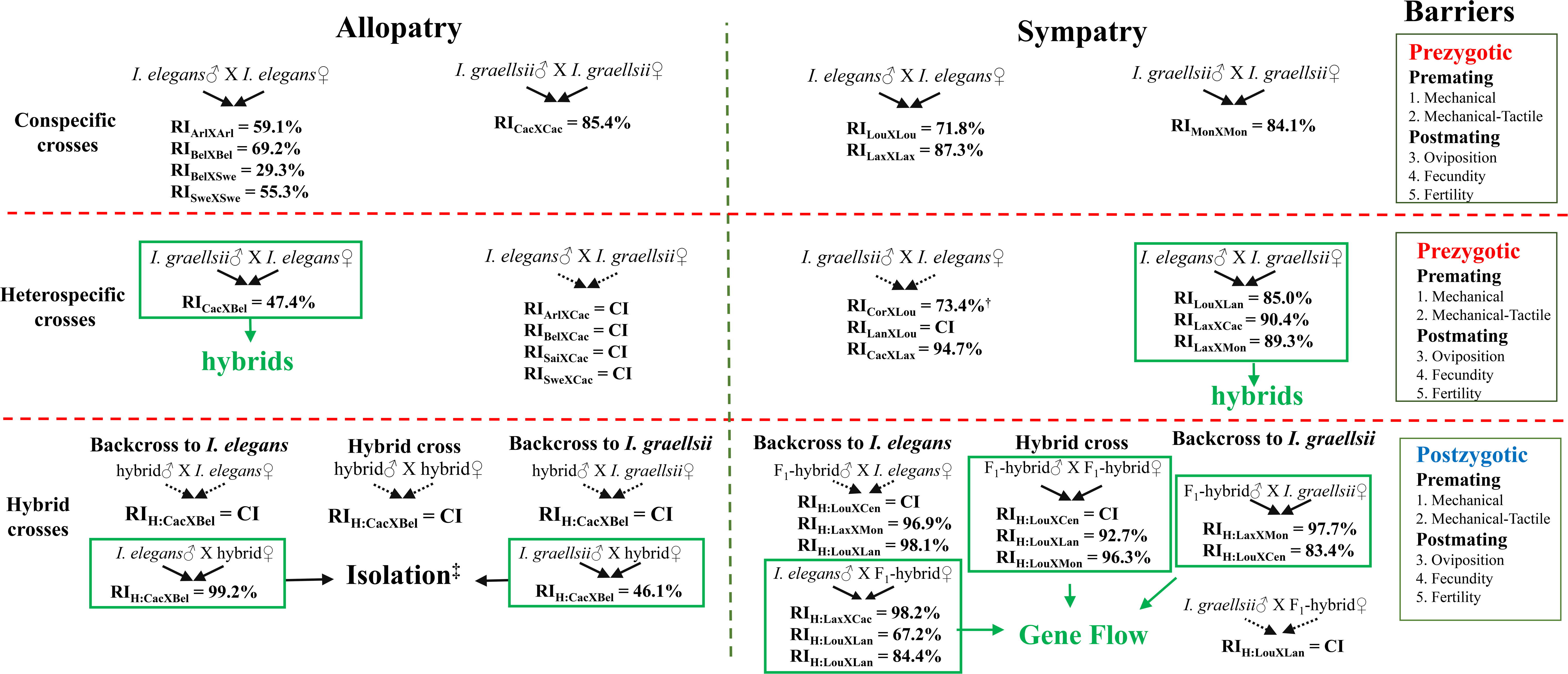
Schematic representation of the hybridization directions between *Ischnura elegans* and *I. graellsii,* comparing crosses between allopatry and sympatry. Solid arrows show gene flow direction and dashed arrows pointing to “CI” letters mark complete isolation. Additionally, we include the total cumulative RI for crosses not in complete isolation. Population labels are explained in Table S1. In allopatry, hybrids were bred only from crosses between *I. graellsii* males and *I. elegans* females, and we could not rear adult F_2_-hybrids. RI was high but not complete in crosses with pure-species males and hybrid females, which leaves the possibility of breeding F_2_-hybrids from these backcrosses. In sympatry, hybrids were bred from crosses between *I. elegans* males and *I. graellsii* females, and most later-generation hybrids were bred from the hybrid crosses and from backcrosses involving *I. elegans* males or *I. graellsii* females ^†^: Crosses between *I. graellsii* males from Corrubedo and *I. elegans* females from Louro did not achieve high total cumulative isolation; however, owing to small sample sizes we could not rear up adult hybrids from this cross. ^‡^: F_2_-hybrid adults from allopatry were not reared because of the low numbers of obtained larvae.

**Figure 4.**
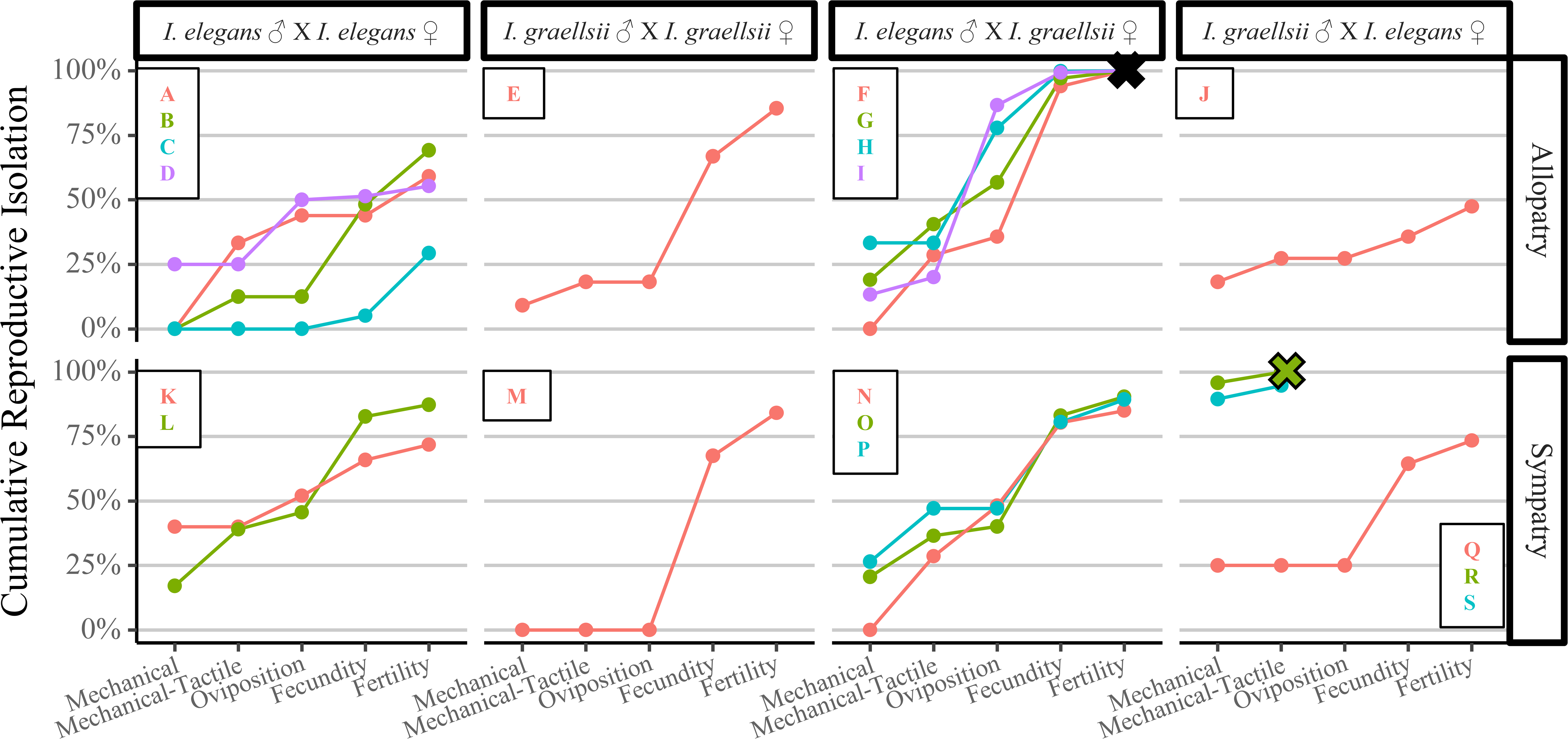
Cumulative RI of five prezygotic barriers in conspecific and heterospecific *Ischnura* crosses from allopatry and sympatry. Only crosses with a sample size equal or higher than 3 during the mechanical barrier were used for cumulative isolation estimates. Color lines within each subplot show data for a population cross pair: A) Arles×Arles; B) Belgium×Belgium; C) Belgium×Sweden; D) Sweden×Sweden; E) Cachadas×Cachadas; F) Arles×Cachadas; G) Belgium×Cachadas; H) SaintCyprien×Cachadas; I) Sweden×Cachadas; J) Cachadas×Belgium; K) Louro×Louro; L) Laxe×Laxe; M) Montalvo×Montalvo; N) Louro×Lanzada; O) Laxe×Cachadas; P) Laxe×Montalvo; Q) Corrubedo×Louro; R) Lanzada×Louro; S) Cachadas×Laxe.

### Reproductive isolation asymmetry

The hybridization direction, i.e., the cross in which hybridization occurs, remained consistent within crosses of different populations within an ecology, but differed between the sympatric and the allopatric ecologies (Fig. 3). In detail, in allopatry, hybridization occurred through crosses between *I. graellsii* males and *I. elegans* females, but was completely precluded in the opposite direction, owing to the cumulative effect of the five measured reproductive barriers (Fig. 4). In contrast, in the sympatry zone, hybridization occurred almost only via *I. elegans* male and *I. graellsii* female crosses. In fact, the mechanical and mechanical-tactile barriers (Fig. 4; Table S2) precluded 94.7% and 100% of the gene flow from the *I. graellsii* males’ and *I. elegans* females’ direction in the sympatry zone in the population crosses of Cachadas and Laxe, and Lanzada and Louro respectively. The exception to this pattern (in crosses between *I. graellsii* males and *I. elegans* females) came from the cross involving Corrubedo and Louro in which total cumulative RI reached only 73.4%. However, since only three females laid eggs in crosses from these populations, low sample sizes precluded us from rearing hybrids from this cross.

Hybrid crosses also differed between allopatric and sympatric ecologies (Fig. 3). In allopatry, matings occurred only via F_1_-hybrid females and *I. elegans* or *I. graellsii* males, i.e., no crosses involving F_1_-hybrid males produced fertile eggs. Although allopatric F_2_-hybrid larvae were bred, the high cumulative RI and low sample sizes made it impossible to obtain any adult F_2_-hybrid. In sympatry, hybrids mated successfully in all cross directions except with *I. graellsii* males. Additionally, RI was complete or almost complete in crosses between hybrid males and *I. elegans* females in all three sympatric interpopulation crosses (Fig. S1). In sympatry, F_2_-hybrids were viable and fertile, and F_3_-hybrids were reared up to adulthood, although no reproductive fitness measurements were made.

### GLM modeling

Prezygotic-barrier GLM modeling and scoring using the AICc suggested that the mechanical barrier was explained by crosses, ecology and the interaction between these two variables (Fig. 5A; Table S3). *Post hoc* comparisons revealed that the heterospecific cross between *I. graellsii* males and *I. elegans* females was significantly different from the other three crosses (p<0.05/6; Table S4). Additionally, significant differences were detected in this cross between the allopatric and sympatric ecology (p<0.05/4; Table S5). In the mechanical-tactile barrier, the null model was selected as the most probable model (Fig. S2A; Table S3). The oviposition barrier was explained by the crosses, ecology and geography (Fig. S2B; Table S3). Finally, both fertility and fecundity barriers were explained by the full model (Figs 5B and 5C; Table S3). All crosses’ fecundities and fertilities were statistically different between allopatry and sympatry (p<0.05/4; Figs. 5B and 5C; Table S5) except the fertility of *I. graellsii* males and *I. elegans* females crosses (p>0.05/4; Table S5).

**Figure 5.**
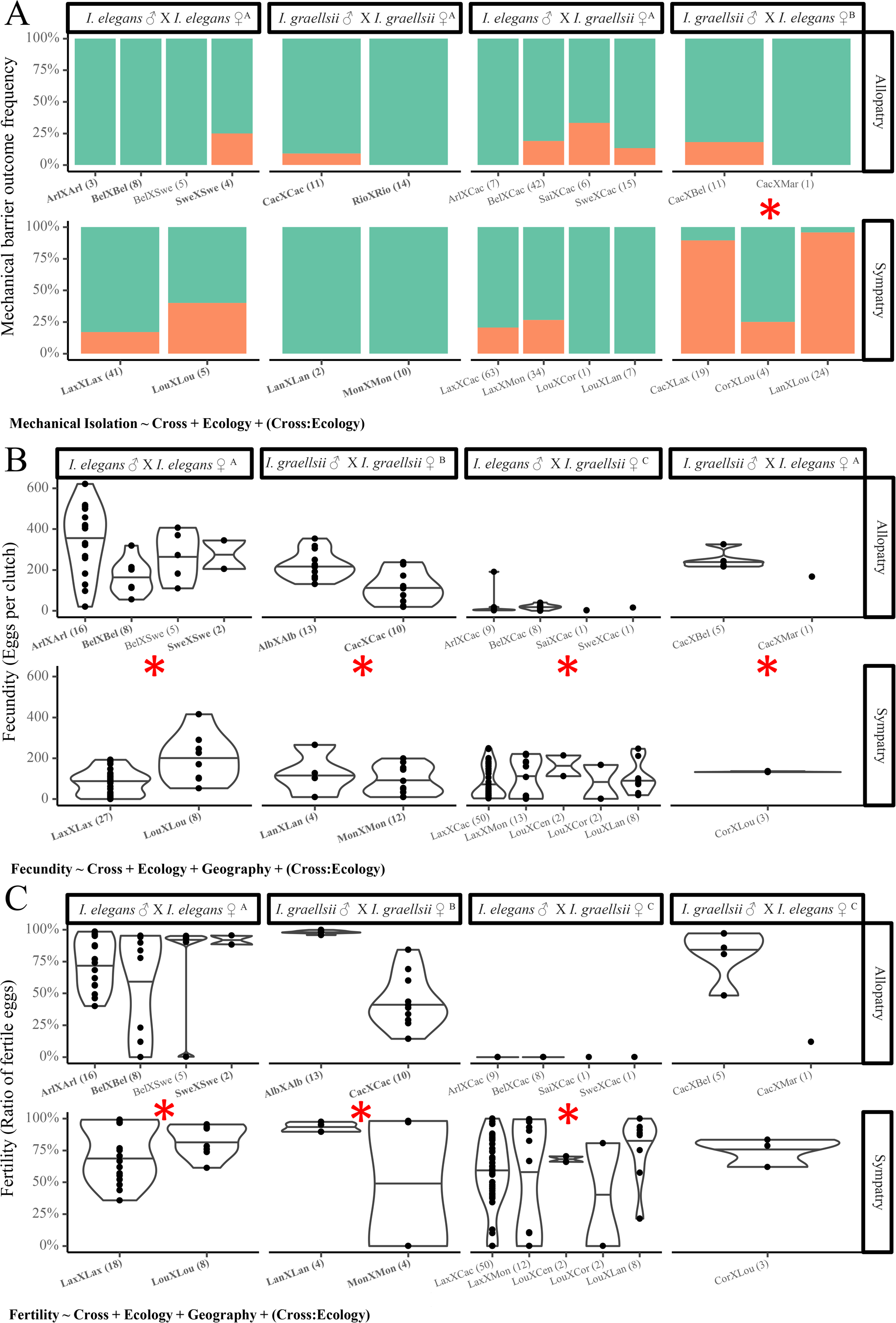
Fitness-component measurements and summary of GLM modeling results for the A) mechanical (green = successful tandem and orange = unsuccessful formation of a tandem), B) fecundity, and C) fertility prezygotic reproductive barriers in *Ischnura*. The equation in the left-bottom corner of each subplot shows the model with the lowest AICc value. Values between parentheses on each population cross show the sample size. Population labels are explained in Table S1. Letters superscripts of crosses boxes at the top of each sublot show different groups inferred with *post hoc* GLM analyses for crosses; e.g., in A) crosses between *I. graellsii* males and *I. elegans* females (*B*) differed significantly in pairwise comparisons from the other three types of crosses (*A*; p<0.05/6). * = *Post hoc* statistically significant differences between the sympatric and allopatric ecology within each cross; **Bold =** Intrapopulation crosses.

Postzygotic-barrier GLM modeling described the mechanical, fecundity and fertility barriers as explained by the crosses, the ecology and the interaction between them (Figs 6 and S3; Table S6). On the other hand, the mechanical-tactile and oviposition barriers were explained only by the ecology (Fig. S3; Table S6). *Post hoc* analyses of the fecundity and fertility barriers showed that each cross had significant differences between allopatry and sympatry (p<0.05/5; Figs. 6A and 6B; Tables S7 and S8).

**Figure 6.**
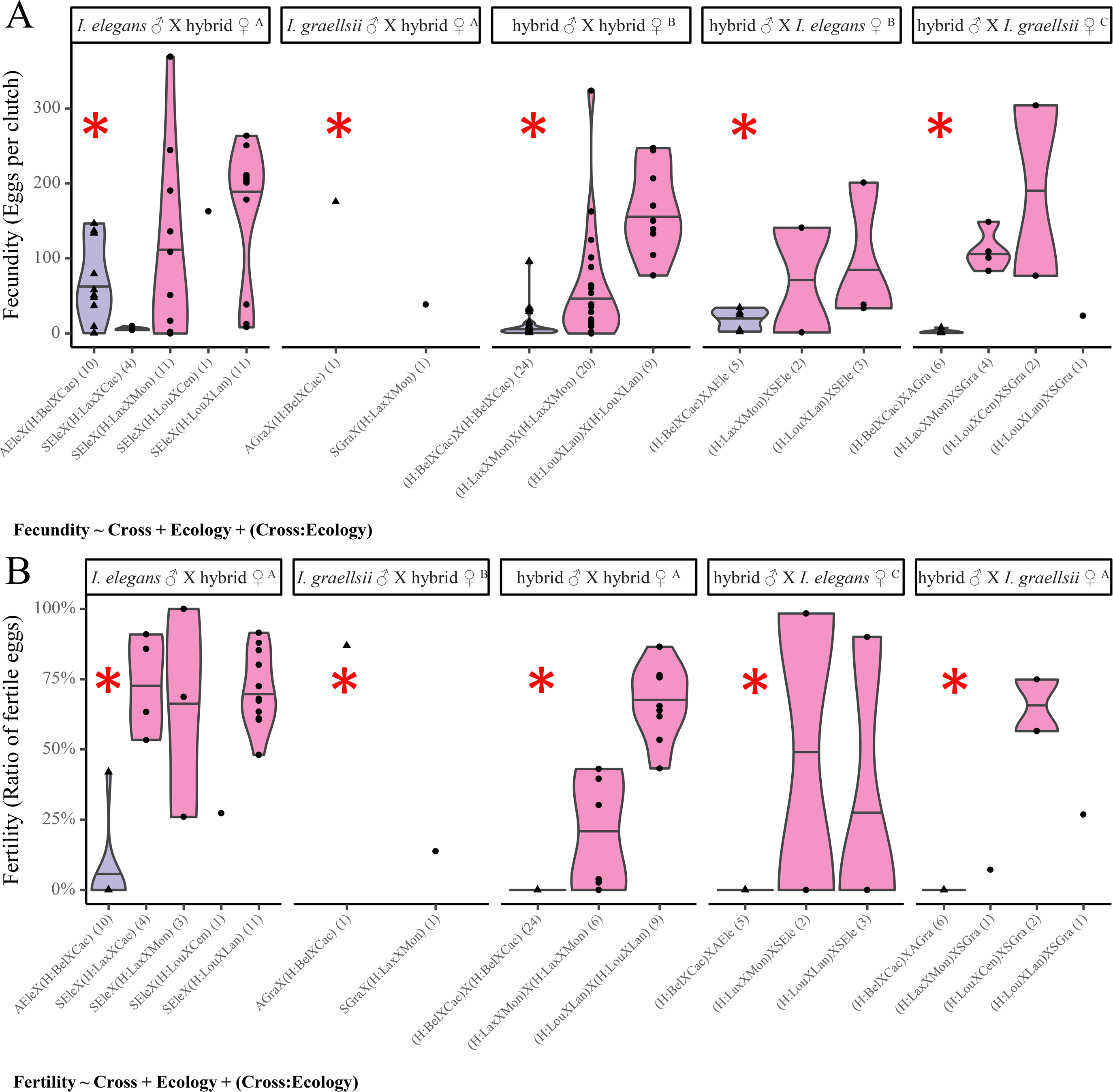
Fitness-component measurements and summary of GLM modeling results for the A) fecundity and B) fertility postzygotic reproductive barriers in *Ischnura*. The equation in the left-bottom corner of each subplot shows the model with the lowest AICc value. Values between parentheses on each population cross show the sample size. Population labels are explained in Table S1 (AEle = Pooled pure allopatric *I. elegans* samples; Agra = Pooled pure allopatric *I. graellsii* samples; SEle = Pooled pure sympatric *I. elegans* samples; SGra = Pooled pure sympatric *I. graellsii* samples). Letters superscripts of crosses boxes at the top of each sublot show different groups inferred with *post hoc* GLM analyses for crosses; e.g., in B) crosses between hybrids with *I. graellsii* males (*B*) and *I. elegans* females (*C*) differed significantly in pairwise comparisons from the other three types of crosses (*A*), and between them (p<0.05/10). Purple and triangles = allopatric crosses; Pink and circles = sympatric crosses; * = *Post hoc* statistically significant differences between the sympatric and allopatric ecology within each cross.

### Testing reinforcement predictions

#### Sympatric strengthening of prezygotic barriers

We detected the classical pattern expected under reinforcement, i.e., stronger prezygotic isolation in sympatry than in allopatry (Table 3; Dobzhansky 1937, 1940), although prezygotic barriers were asymmetric between heterospecific reciprocal crosses. Between *I. graellsii* males and *I. elegans* females, total prezygotic isolation was stronger in sympatry than in allopatry. The exception was the sympatric cross between Corrubedo and Louro, in which prezygotic isolation was similar to allopatry. In the reciprocal cross direction, between *I. elegans* males and *I. graellsii* females, cumulative prezygotic isolation was similar between sympatry and allopatry, although in the latter all population crosses reached complete isolation (Fig. 4).

We detected statistically significant differences in the strength of the mechanical barrier in crosses between *I. graellsii* males and *I. elegans* females in comparison to the reciprocal cross direction and the conspecific crosses of *I. elegans* and *I. graellsii* (Fig. 5A; Table S4). *Post hoc* GLM modeling revealed that in this cross mechanical isolation was stronger in sympatry than in allopatry (Fig. 5A; Table S5). Strong mechanical isolation in crosses between *I. graellsii* males and *I. elegans* females was seen in two out of the three sympatric interpopulation crosses (Fig. 5A). In contrast, in crosses between *I. elegans* males and *I. graellsii* females mechanical isolation was similar to that in the conspecific crosses (Fig. 5A).

#### Rarer female effect

Reinforcement theory predicts that selection will strengthen prezygotic barriers in the cross direction which includes females of the less abundant species (Table 3; Yukilevich 2012). We could not test this prediction as allopatric crosses showed strong asymmetry between both crosses directions (Figs. 1 and 4). Since crosses between *I. elegans* males and *I. graellsii* females were completely isolated in the allopatric condition reinforcement pressures could have only operated in sympatry in the opposite direction (i. e. between *I. graellsii* males and *I. elegans* females) independently of the relative abundance of both species in the sympatry zone.

#### Concordant prezygotic and postzygotic isolation asymmetries

Since costs of hybridization (postzygotic barriers) are usually asymmetric between reciprocal crosses (termed Darwin’s corollary; Darwin 1859; Turelli and Moyle 2007), reinforcement is predicted to be more intense in the reciprocal cross direction which produces more unfit hybrids (Table 3; Yukilevich 2012). Thus, concordant prezygotic and postzygotic isolation asymmetries are expected between reciprocal crosses in sympatry but not in allopatry. This prediction could not be tested in our data because RI was complete in the cross direction between *I. elegans* males and *I. graellsii* females in allopatric crosses (Fig. 3). Additionally, although the reciprocal cross direction, the one between *I. graellsii* males and *I. elegans* females, was not completely precluded in either of the zones (allopatry and sympatry), its high strength and the low sample size of the obtained larvae precluded us to rear them until adulthood.

#### Greater premating asymmetries and weaker postzygotic isolation

Turelli et al. (2014) proposed two additional predictions of the reinforcement theory, such as a more definitive test of Yukilevich (2012) hypothesis about the role of intrinsic postzygotic isolation in reinforcement (concordant isolation asymmetries). They proposed that species pairs that have asymmetric postzygotic barriers in sympatry: i) should present greater premating asymmetries in sympatry than in allopatry; and ii) since allopatrically originated Bateson-Dobzhansky-Müller (BDM) incompatibility alleles are purged in sympatry because of gene flow, species should present a reduction in the strength of intrinsic postzygotic isolation in sympatry relative to allopatry (Table 3; Turelli et al. 2014). Additionally, postzygotic isolation asymmetries should be reduced in sympatry by purging of unidirectionally inherited BDM incompatibilities (Turelli and Moyle 2007). We could not fully test Turelli’s predictions since prezygotic RI was complete (or very high) in one cross direction in both sympatry and allopatry, impeding our ability to test the asymmetry of the postzygotic barriers between reciprocal crosses. However, we compared prezygotic asymmetries between sympatry and allopatry, and estimated an overall measurement of postzygotic isolation using each of the hybrids formed in allopatry and sympatry. We detected greater prezygotic-premating asymmetries (Fig. S4, Table S9), weaker prezygotic-postmating isolation (Fig. 5B; Table S5) and weaker postzygotic isolation in sympatry than in allopatry (Fig. 6; Table S8); however, evidence from this prediction should be taken carefully owing to the assumption we could not fulfill.

Firstly, consistent with the prediction of higher prezygotic-premating asymmetries in sympatry than in allopatry, we detected significant asymmetries in the mechanical barrier in sympatry using all the sympatric data (Fig. S4B; Table S9) and between the reciprocal heterospecific crosses of Lanzada and Louro (Fig. S4E; Table S9), and Cachadas and Laxe (Fig. S4F; Table S9). The exception was between the reciprocal heterospecific crosses of Louro and Corrubedo (Fig. S4D; Table S9). In allopatry, neither by using all data (Fig. S4A; Table S9) nor with the reciprocal heterospecific crosses of Cachadas and Belgium (Fig. S4C; Table S9) were significant premating asymmetries detected. Additionally, all prezygotic barriers in allopatry and prezygotic-postmating barriers in sympatry were stronger in crosses between *I. elegans* males and *I. graellsii* females; however, in sympatry the mechanical barrier was stronger in crosses between *I. graellsii* males and *I. elegans* females (Fig. S4).

Secondly, despite the fact that we could not test the asymmetry of the postzygotic barriers between reciprocal crosses, we detected overall weaker postmating isolation in hybrids from sympatry than from allopatry. While in allopatry all crosses with hybrid males produced a low number of infertile eggs and no F_2_-hybrids could be reared up, in sympatry only crosses between *I. graellsii* males and hybrids were completely isolated and adult F_2_-hybrids could not be bred, reared-up and reproduced (Figs 3 and S2). In all five reproductive barriers the ecology was a significant factor influencing postzygotic isolation (Figs 6 and S3; Table S6), although its effects differed between reproductive barriers. While postzygotic-premating barriers were usually stronger in sympatry than in allopatry (Figs S3A and S3B), in all three postzygotic-postmating barriers allopatric crosses presented stronger isolation than sympatric crosses (Figs 6 and S3C). In fact, four out of the five postzygotic types of crosses presented higher fecundities and fertilities values in sympatric crosses than in allopatric crosses (Fig. 6). The exception was in crosses between *I. graellsii* males and hybrid females that had very low sample sizes both for the allopatric and sympatric ecology (Fig. 6; Table S2).

Interestingly, significant differences between the allopatric and sympatric ecology were also detected in prezygotic-postmating barriers. In conspecific crosses data distribution shows that in sympatry pure crosses produce a lower number of eggs than in allopatry (Fig. 5B), although no clear pattern could be inferred between allopatry and sympatry for fertility values (Fig. 5C). However, heterospecific crosses between *I. elegans* males and *I. graellsii* females presented an increment of both fecundity and fertility in sympatry than in allopatry (Fig. 5). Since Turelli et al. (2014) prediction is based on evidencing gene flow in sympatric heterospecific crosses, this pattern of increased fecundity and fertility in sympatric crosses between *I. elegans* males and *I. graellsii* females could be evidence of the homogenizing effects of historical gene flow in this direction. This is consistent with the fact that sympatric hybridization occurs in this direction (Fig. 3). Recent genomic evidence has shown reduced heterospecific differentiation and increased intraspecific genetic diversity in both *I. elegans* and *I. graellsii* in sympatric samples in comparison to allopatric samples (Sánchez-Guillén et al. 2023), which strengthens the evidence for heterospecific gene flow.

Finally, we not only detected statistically significant asymmetries in sympatry (but not in allopatry) in the mechanical barrier, but also found that the strength of this barrier shifted from being stronger in allopatry in crosses between *I. elegans* males and *I. graellsii* females to being stronger in sympatry in crosses between *I. graellsii* males and *I. elegans* females (Fig. S4). Additionally, sympatric backcrosses (but not allopatric) were successful in a similar way, as were heterospecific crosses in the first generation. Specifically, while hybridization occurred in sympatry in crosses between *I. elegans* males and *I. graellsii* females, backcrosses were successful mostly with either *I. elegans* males or *I. graellsii* females (Fig. 3). On the other hand, backcrosses with *I. graellsii* males or *I. elegans* females were prevented by a strong mechanical barrier (Fig. S1). This pattern suggests that if reinforcement has occurred in the mechanical isolation of *I. graellsii* males and *I. elegans* females, then mechanical isolation could also have been strengthened in backcrosses involving *I. graellsii* males and *I. elegans* females.

## Discussion

Although our data were inconclusive in testing several reinforcement theoretical predictions, our results suggest the presence of reinforcement (Table 3). This is consistent with morphological evidence of RCD in sympatric *I. elegans* and *I. graellsii* females (Ballén-Guapacha *et al., in press*). We detected stronger prezygotic isolation in crosses between *I. graellsii* males and *I. elegans* females in sympatry than in allopatry owing to the strengthening of the mechanical barrier in these crosses. We also identified stronger premating asymmetries in sympatry than in allopatry, an evidence of sympatric gene flow in the form of reduced prezygotic-postmating barriers in sympatry than in allopatry, and similar patterns of premating barriers in prezygotic and postzygotic barriers; i.e., the same mating directions in heterospecific and backcrosses in sympatry but not in allopatry. Data of two out of three population crosses in sympatry revealed a consistent pattern of reinforcement.

### Evolution of mechanical isolation in sympatry

The relative contributions of the five reproductive barriers to RI differed between allopatry and sympatry and between reciprocal heterospecific crosses. In allopatry, premating (mechanical and mechanical-tactile) barriers were moderate and similar between reciprocal crosses, while postmating (oviposition, fecundity, and fertility) barriers were strong and highly asymmetric between reciprocal crosses, preventing 100% of the hybrid formation between *I. elegans* males and *I. graellsii* females. In two out of the three sympatric crosses between *I. graellsii* males and *I. elegans* females, premating barriers were stronger than postmating barriers, and most of the isolation was due to the action of the mechanical barrier preventing the tandem formation. The low mechanical isolation detected in the heterospecific crosses involving *I. graellsii* males from Corrubedo (Fig. 4Q) could be due to a misclassification of hybrids as *I. graellsii*, because of the high prevalence of hybrids in this population during the sampling year (Table S1). In the cross direction between *I*. *elegans* males and *I*. *graellsii* females, gene flow was prevented by the joint action of both premating and postmating barriers.

Mechanical and mechanical-tactile barriers preventing the formation of successful tandem or copula formation are (with a few exceptions; Nava-Bolaños et al. 2017) important reproductive barriers across a variety of non-territorial odonate species, such as the *Enallagma* and *Ischnura* damselflies, which lack visual recognition and precopulatory courtship behaviors (Robertson and Paterson 1982; Barnard et al. 2017; Solano et al. 2018). The role of mechanical barriers in RI has been used as evidence for the lock-and-key model (Paulson 1974; Eberhard 1985; Masly 2012), which suggests that the morphology of sexual structures is under rapid male-female coevolution via reinforcement to enhance RI (Eberhard 1985; Masly 2012), and explains the wide diversity and taxonomic importance of sexual structures (Monetti et al. 2002; Barnard et al. 2017; Solano et al. 2018). Thus, the lock-and-key theory predicts enhanced mechanical isolation in sympatry compared with allopatry, and a correlation with low hybrid fitness (Eberhard 1985; Shapiro and Porter 1989; Brennan and Prum 2015). Our results are consistent with both predictions and suggest that sexual structures involved in the tandem formation could be evolving because of reproductive character displacement (RCD) in *I. elegans* and *I. graellsii*. This is consistent with recent morphological evidence showing RCD in the pronotum of females in sympatry (Ballén-Guapacha *et al., in press*). RCD in these structures could also explain why premating barriers in sympatry behaved similarly in backcrosses and in heterospecific crosses, i.e., reducing gene flow in backcrosses with *I. graellsii* males or *I. elegans* females. If tandem related structures have mainly been reinforced in *I. graellsii* males and *I. elegans* females, then these structures could also be mechanically incompatible with hybrids with intermediate morphology. This provides an explanation to why sympatric backcrossing occurred mainly with *I. elegans* males or *I. graellsii* females.

### Testing specific predictions of reinforcement

Refinements of the reinforcement theory during the 1990s concluded that reinforcement could occur under a broad range of conditions (Coyne and Orr 2004), although several factors need to be fulfilled. For example, the outcomes of hybridization would range from species fusion and extinction to speciation via reinforcement as a function of hybridization costs and initial differences in reproductive characteristics between species (Liou and Price 1994). The higher the hybridization costs (lower fitness of hybridizing individuals), and the higher the initial variance in reproductive characteristics, the higher the probability of speciation via reinforcement (Liou and Price 1994). In allopatry, the cross between *I. elegans* males and *I. graellsii* females is completely isolated by prezygotic barriers. Unsuccessful mating attempts (complete prezygotic isolation) can still act as a selective pressure that strengthens earlier-acting barriers (e.g., premating barriers) to avoid unnecessary wastage of gametes, time, energy (Hoskin and Higgie 2013), or other reproductive costs. However, reinforcement pressures increase as further reproductive barriers act on hybridization. Intrinsic postzygotic isolation is usually more costly (at least to females) than prezygotic isolation, as energy has been invested in maladaptive hybrid formation (Ortiz-Barrientos et al. 2009). Data from allopatric populations showed that crosses between *I. graellsii* males and *I. elegans* females are more prone to be reinforced than the opposite direction, based on the formation of costly F_1_-hybrids which are highly unfit owing to their high infertility, and because both species are morphologically well differentiated by reproductive characters related to the tandem position, i.e., male caudal appendages and female pronotum (Monetti et al. 2002). Importantly, the fact that hybrids from the allopatric crosses between *I. graellsii* males and *I. elegans* females are highly, but not completely, unfit (not achieving complete isolation in the F_1_ generation) suggests that some gene flow is possible, and that these species are not yet “good” species *sensu* Butlin (1987). This distinction is important, as several authors (Butlin 1987; Coyne and Orr 2004) argue that sympatric strengthening of prezygotic isolation in cases in which taxa already produce completely unfit hybrids (no gene flow) in allopatry should not be considered as reinforcement, since such enhancement of prezygotic isolation would have then happened *after* allopatric speciation. Consistently, our sympatric experiments crosses showed evidence that reinforcement has strongly enhanced the prezygotic RI between *I. graellsii* males and *I. elegans* females. This is evident by a stronger prezygotic isolation between *I. graellsii* males and *I. elegans* females in sympatry than in allopatry due to the strengthening of the mechanical barrier.

We could not test neither the rarer female effect nor the “concordant isolation asymmetries” predictions (Yukilevich 2012), and we could only test partially the greater premating asymmetries and weaker postzygotic isolation in sympatry than in allopatry pattern (Turelli et al. 2014) because F_1_-hybrids from one cross direction, in both allopatry and sympatry, were not obtained due to the completeness of the prezygotic isolation. However, the shift in hybridization directions between allopatry and sympatry, the higher mechanical isolation in the latter than the former, and recent evidence of higher RCD in *I. elegans* females than in *I. graellsii* males (Ballén-Guapacha *et al., in press*) is all consistent with the reinforcement of reproductive isolation theory. Future studies should increase the sample size of experimental crosses in an attempt to obtain F_1_ hybrids from both reciprocal cross directions. This will open the possibility to test the predictions that we could not.

Our results show that reinforcement can act rapidly, since differences in prezygotic isolation have been formed at most during the last 100–120 years since the presence of *I. elegans* was detected in Spain (Sánchez-Guillén et al. 2011, 2023; Wellenreuther et al. 2018). Our results are consistent with *Drosophila* experiments showing that reinforcement can act rapidly in just a few generations (Matute 2010a). Additionally, our data show that reinforcement can quickly shift hybridization directions, i.e., from hybridization occurring between *I. graellsii* males and *I. elegans* females in allopatry to between *I. elegans* males and *I. graellsii* females in sympatry. This could be, to our knowledge, the first report of such natural shifting in hybridization directions in a time scale of between tens to some hundreds of generations due to reinforcement. We hypothesize that during the initial secondary contact between *I. elegans* and *I. graellsii,* hybridization should have occurred in the allopatric direction, i.e., between *I. graellsii* males and *I. elegans* females. High hybridization costs of this cross direction (infertile hybrid males) could have induced reinforcement to displace tandem-related reproductive characters in *I. elegans* females (Ballén-Guapacha *et al., in press*), reducing the mechanical compatibility between *I. graellsii* males and *I. elegans* females. However, as introgression occurred between the species (Sánchez-Guillén et al. 2023), purging of BDM incompatibilities reduced postzygotic isolation in sympatry, and reduced heterospecific genetic differentiation could have reduced prezygotic-postmating isolation by increasing heterospecific fecundity and fertility. Since reinforcement could have been occurring mostly between *I. graellsii* males and *I. elegans* females, both the reduction of prezygotic-postmating and postzygotic isolation could have allowed sympatric hybridization to occur in crosses between *I. elegans* males and *I. graellsii* females. Once sympatric hybridization was possible between *I. elegans* males and *I. graellsii* females, reinforcement in this cross direction could occur, albeit slower than in *I. elegans* females because hybridization costs (postzygotic isolation) have been reduced. Whether introgression will increase by hybridization between *I. elegans* males and *I. graellsii* females, or reinforcement will increase prezygotic isolation also in this direction, is an interesting question to evaluate in the future.

While asymmetrically reinforcement has been documented before (Jaenike et al. 2006; Turner et al. 2010; Yukilevich 2012; Zhou and Fuller 2014; Ostevik et al. 2021; St. John and Fuller 2021), to our knowledge this could be the first study suggesting reinforcement and gene flow causing opposite consequences between reciprocal crosses, i.e., reinforcement increasing prezygotic isolation in one direction and gene flow reducing in the other. Future studies should evaluate the asymmetrical effects of reinforcement and gene flow between reciprocal crosses in species pairs in which asymmetrical reinforcement has been documented.

### Weakening of intrinsic postzygotic isolation

In addition to the evidence of reinforcement of mechanical isolation, we detected weaker postzygotic-postmating isolation, and a lower number of hybrid crosses completely isolated by postmating barriers in sympatry than in allopatry. Hybrid fecundity and fertility fitness relative to those of pure species are mixed, and highly dependent on the genetic divergence between the parental species (Burke and Arnold 2001; Orr and Turelli 2001). They range from: i) reductions in both F_1_ and F_2_ hybrids fecundity or fertility (Naisbit et al. 2002); ii) no differences in fecundity and fertility between the parental species and hybrids (Van Der Sluijs et al. 2008); to iii) equal or higher F_1_-hybrid reproductive success than conspecific crosses but lower in F_2_ or later generation hybrids (hybrid breakdown; Vetukhiv 1956; Edmands 1999; Dunham and Argue 2000). Reductions in hybrid fecundity or fertility are best explained by the Bateson-Dobzhansky-Müller (BDM) incompatibilities model (Dobzhansky 1934; Orr 1996). That model describes how reductions in hybrid fitness occur in response to negative interactions between introgressed alleles from different populations and the genomic background of hybrids. Hybrid breakdown due to BDM incompatibilities is more prone to occur as species diverge, accumulate mutations and increase in genetic distance (Orr and Turelli 2001). Despite conflicting evidence as to whether BDM incompatibilities accumulate linearly (Leppälä et al. 2013) or faster (i.e. the snowball effect; Orr 1995; Presgraves 2010) over time, empirical research in both plants (Moyle and Nakazato 2010; Leppälä et al. 2013) and animals (Matute et al. 2010) converges to a continuous accumulation of BDM incompatibilities as taxa diverge. This BDM incompatibilities property, i.e., higher frequency at increased genetic divergence, is consistent with our observations. Overall genetic distance between *I. elegans* and *I. graellsii* in the north-west hybrid zone (F_ST_=0.625) is lower than in allopatry (F_ST_=0.725) (Sánchez-Guillén et al. 2023). However, future studies should attempt to rear up hybrids from *I. elegans* and *I. graellsii* in both cross directions in both ecologies to help distinguish whether the sympatric reduction of postzygotic isolation in sympatry is due to purging via gene flow (Turelli et al. 2014), because species inherit BDM incompatibilities asymmetrically (Turelli and Moyle 2007), or a combination of both of these factors.

Future studies should also formally evaluate the genetic bases of these apparent BDM incompatibilities. Since in allopatric heterospecific crosses, male hybrids were infertile, and since males are the hemizygous sex in these species, some of these BDM incompatibilities may be related to the X chromosome (Haldane’s rule). These results are consistent with recent evidence suggesting a role of the X chromosome in the reproductive isolation of these species (Swaegers et al. 2022). Evidence gathered since the origin of the Haldane’s rule in 1922 (Haldane 1922) has established this phenomenon as one of the most robust generalizations in evolution (Delph and Demuth 2016), i.e., that hybrids from the heterogametic (or hemizygous; Koevoets and Beukeboom 2009) sex are the ones with reduced fitness. Not only are there plenty of cases reported in vertebrates, invertebrates and plants (reviewed in Schilthuizen et al. 2011; Delph and Demuth 2016), but also recent evidence has shown that there are a high number of independent evolutionary origins of the Haldane’s rule (Delph and Demuth 2016).

## Conclusions

Our results provide not only new empirical evidence of reinforcement of RI in Odonata, but also contribute to a better understanding of the mechanisms leading to speciation, by describing a natural model in which several mechanisms such as reinforcement, Bateson-Dobzhansky-Müller incompatibilities and the Haldane’s rule are driving RI simultaneously. Our work describes a case where reinforcement increases prezygotic isolation in one cross direction, while simultaneously, gene flow weakens postzygotic isolation in the opposite cross direction. Since the study of the asymmetrical effects of reinforcement between reciprocal crosses (Jaenike et al. 2006; Turner et al. 2010; Yukilevich 2012; Zhou and Fuller 2014; Ostevik et al. 2021; St. John and Fuller 2021) is an important growing field in evolutionary biology, our study opens the possibility of testing the interaction between these processes in other taxa.

## Data availability

All datasets and scripts used in this manuscript were uploaded to OSF at: https://osf.io/k6jyg/?view_only=c68a5102dea44045ab9dd922c425e7f3

## Acknowledgements

The research was funded by the Mexican CONACYT (282922 to RAS-G), the Royal Physiological Society in Lund, the Nilsson-Ehle Foundation (36118 to RAS-G), the Marie Curie Intraeuropean Fellowship (to RAS-G, MW and BH), and the Swedish Research Council (621-2016-689 to BH). LRA-V and AVB-G received PhD student grants by the Mexican CONACyT. We are very grateful to Adolfo Cordero Rivera, who kindly allowed us to use his laboratory (Ecología Evolutiva y Conservación, Universidad de Vigo) and material for rearing experiments and to Janet Nolasco-Soto for technical support and help with data collection. Maria L. Castillo, Diana E. Carrillo-Lara and Rodrigo Lasa-Covarrubias helped us with materials and advice during the 2019–2020 damselfly rearing experiment. We are also very grateful with Janne Swaegers and Enrique Ibarra-Laclette who took part in LRA-V’s thesis committee. This manuscript is part of LRA-V’s PhD thesis.

---

**Fig. S1.**
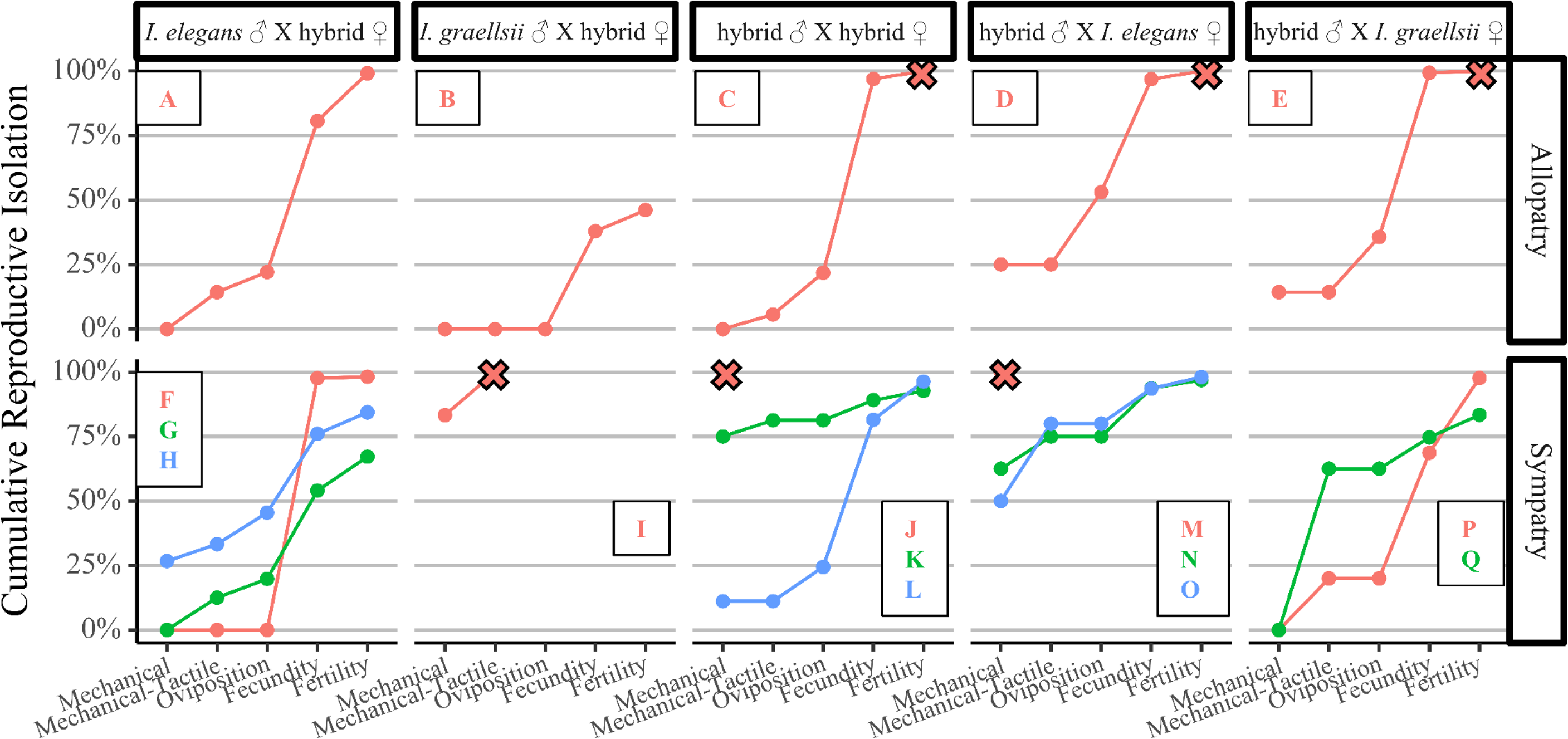
Cumulative RI of five postzygotic barriers in conspecific and heterospecific crosses from allopatry and sympatry. Color lines within each subplot show data for a population cross pair: A) AElle×(H:Bel×Cac); B) AGra×(H:Bel×Cac); C) (H:Bel×Cac)×(H:Bel×Cac); D) (H:Bel×Cac)×AElle; E) (H:Bel×Cac)×AGra; F) SEle×(H:Lax×Cac); G) SEle×(H:Lou×Lan); H) SEle×(H:Lax×Mon); I) SGra×(H:Lou×Lan); J) (H:Lou×Cen)×(H:Lou×Cen); K) (H:Lou×Lan)×(H:Lou×Lan); L) (H:Lou×Mon)×(H:Lou×Mon); M) (H:Lou×Cen)×SEle; N) (H:Lax×Mon)×SEle; O) (H:Lou×Lan)×SEle; P) (H:Lax×Mon)×SGra; Q) (H:Lax×Cen)×SGra.. Population labels are explained in the Table S1 (AEle = Pooled pure allopatric *I. elegans* samples; Agra = Pooled pure allopatric *I. graellsii* samples; SEle = Pooled pure sympatric *I. elegans* samples; SGra = Pooled pure sympatric *I. graellsii* samples).

**Fig. S2.**
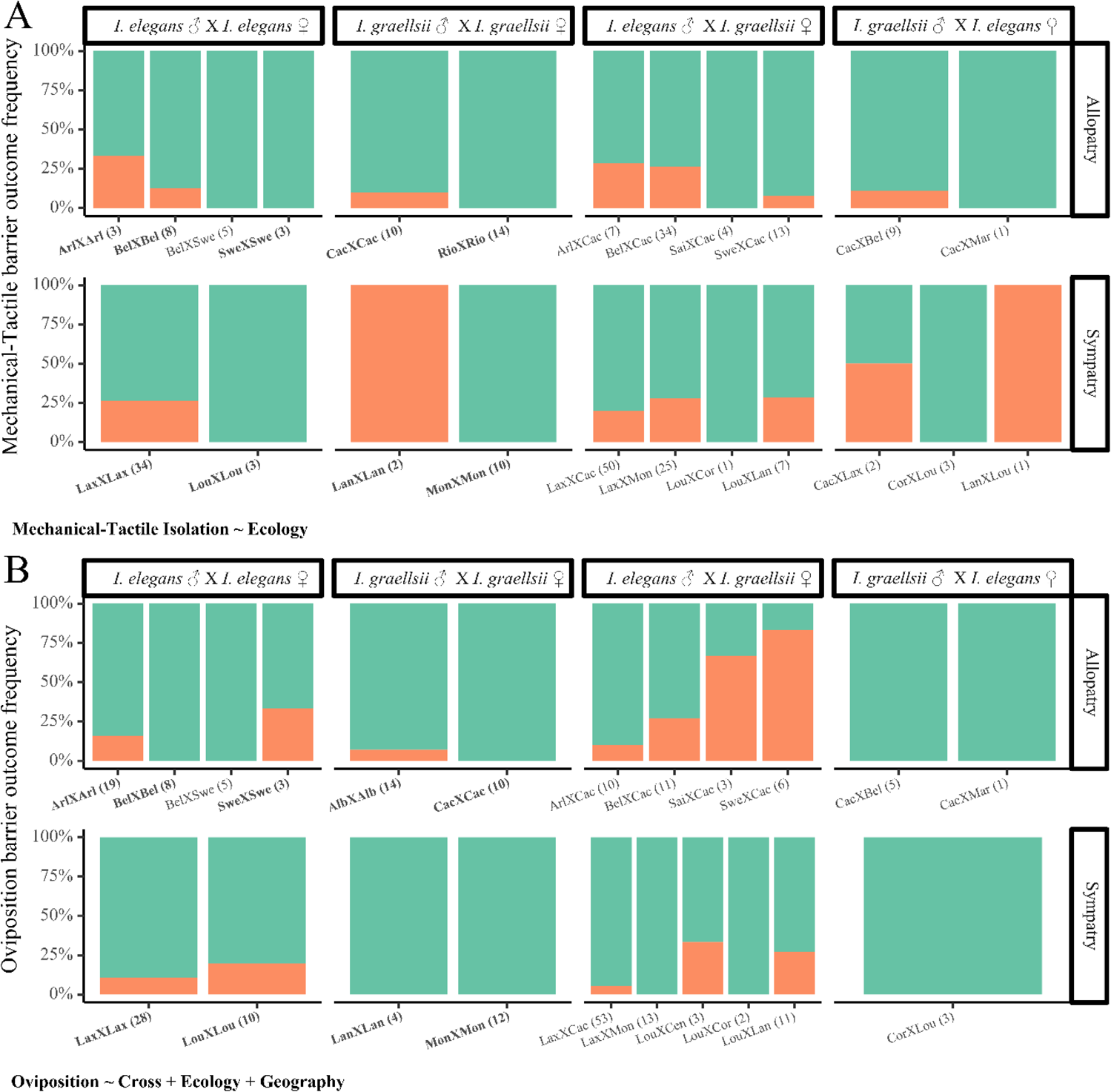
Fitness components measurements for *Ischnura* and summary of GLM modeling results for the A) mechanical-tactile (green = successful mating and orange = unsuccessful mating) and B) oviposition prezygotic reproductive barriers (green = successful oviposition and orange = unsuccessful oviposition). The equation in the left-bottom corner of each subplot shows the model with the lowest AICc value. Values between parentheses on each population cross show the sample size. Population labels are explained in the Table S1. No statistically significant differences between crosses were detected with *post hoc* GLM (p > 0.05/6). **Bold** = Intrapopulation crosses.

**Fig. S3.**
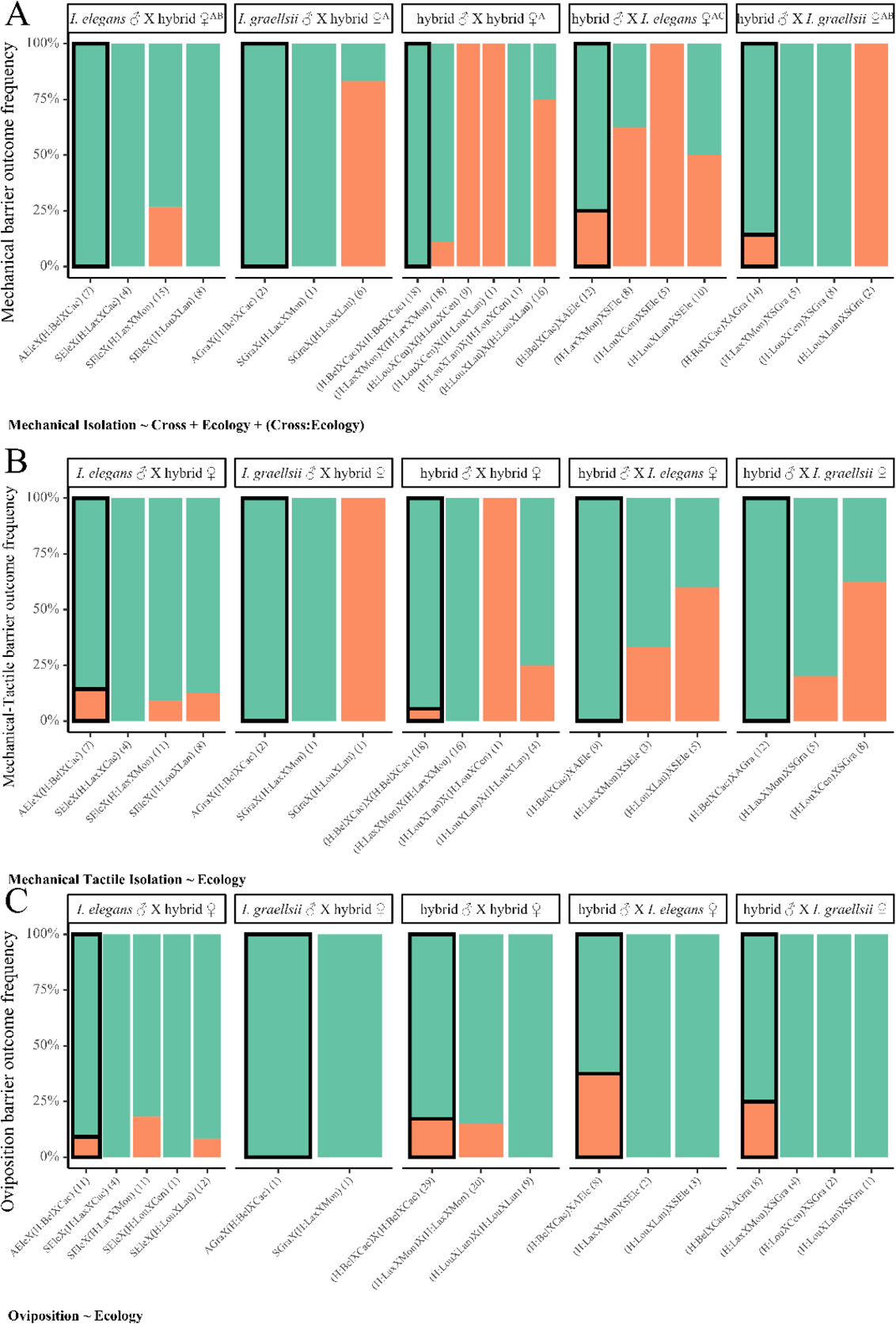
Fitness components measurements for *Ischnura* and summary of GLM modeling results for the A) mechanical (green = successful tandem and orange = unsuccessful tandem), B) mechanical-tactile (green = successful mating and orange = unsuccessful mating) and C) oviposition (green = successful oviposition and orange = unsuccessful oviposition) postzygotic reproductive barriers. The equation in the left-bottom corner of each subplot shows the model with the lowest AICc value. Values between parentheses on each population cross show the sample size. Population labels are explained in the Table S1 (AEle = Pooled pure allopatric *I. elegans* samples; Agra = Pooled pure allopatric *I. graellsii* samples; SEle = Pooled pure sympatric *I. elegans* samples; SGra = Pooled pure sympatric *I. graellsii* samples). Letters superscripts of crosses boxes at the top of each sublot show different groups inferred with *post hoc* GLM analyses for crosses. In A) crosses between hybrids and *I. elegans* females (*AC*) differed significantly in pairwise comparisons from their reciprocal cross and from crosses between hybrids and *I. graellsii* females (*AB*; p<0.05/10).

**Fig. S4.**
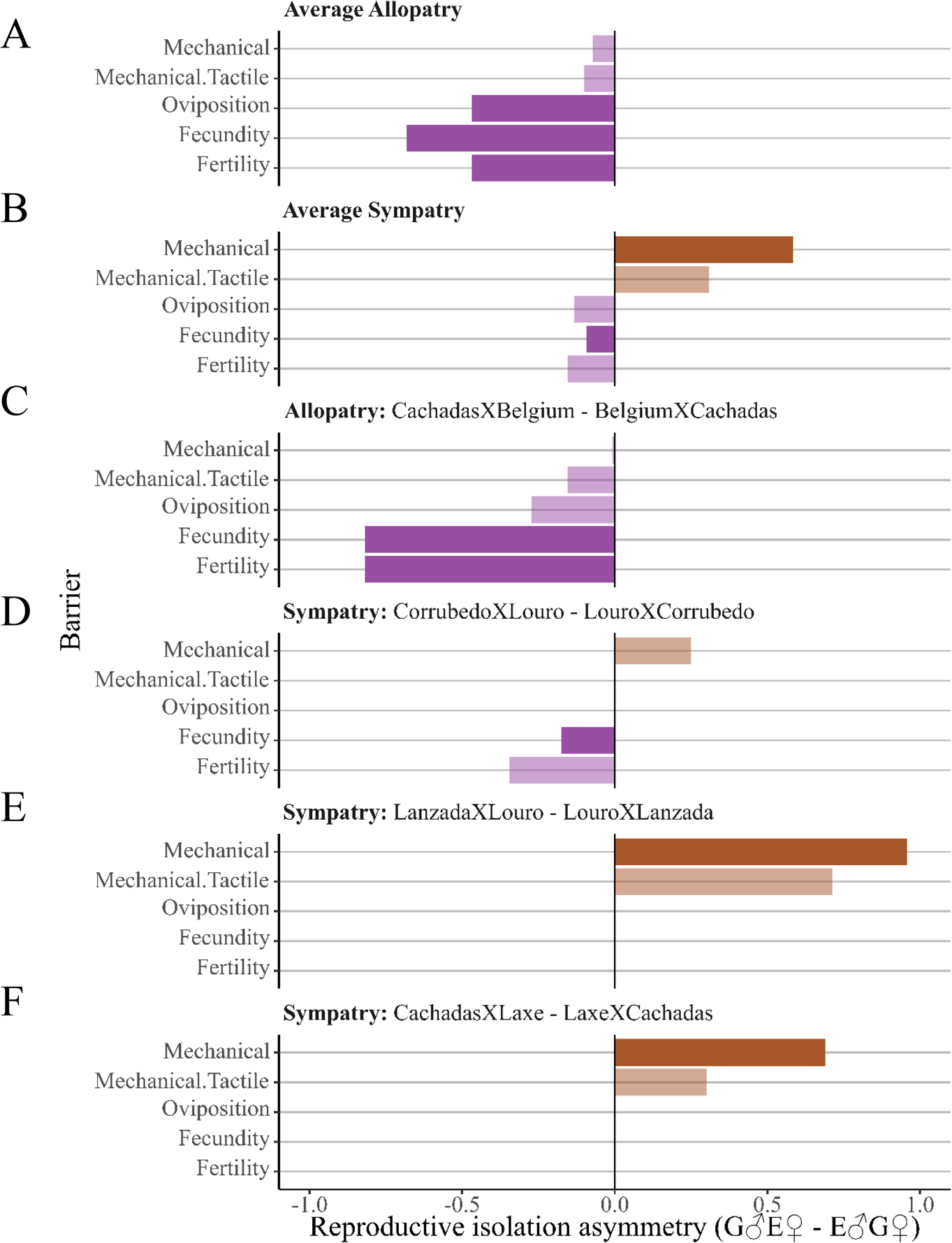
Prezygotic reproductive asymmetries in A) allopatry (averaged between all populations), B) sympatry (averaged between all populations) and populations in which both reciprocal crosses were sampled both in allopatry and sympatry (E to F). Asymmetries were measured as the absolute RI from crosses between *Ischnura graellsii* males with *I. elegans* females minus the RI in the reciprocal direction and are categorized between stronger isolation in the former (brown) or stronger isolation in the latter (purple). Solid bars represent barriers in which GLM models including the cross variable presented lower AICc values than models excluding it. GLM modeling for allopatry and sympatry were done using all data; the plot shows the difference between population crosses averages. E: *I. elegans*; G: *I. graellsii*.

**Table S1.**
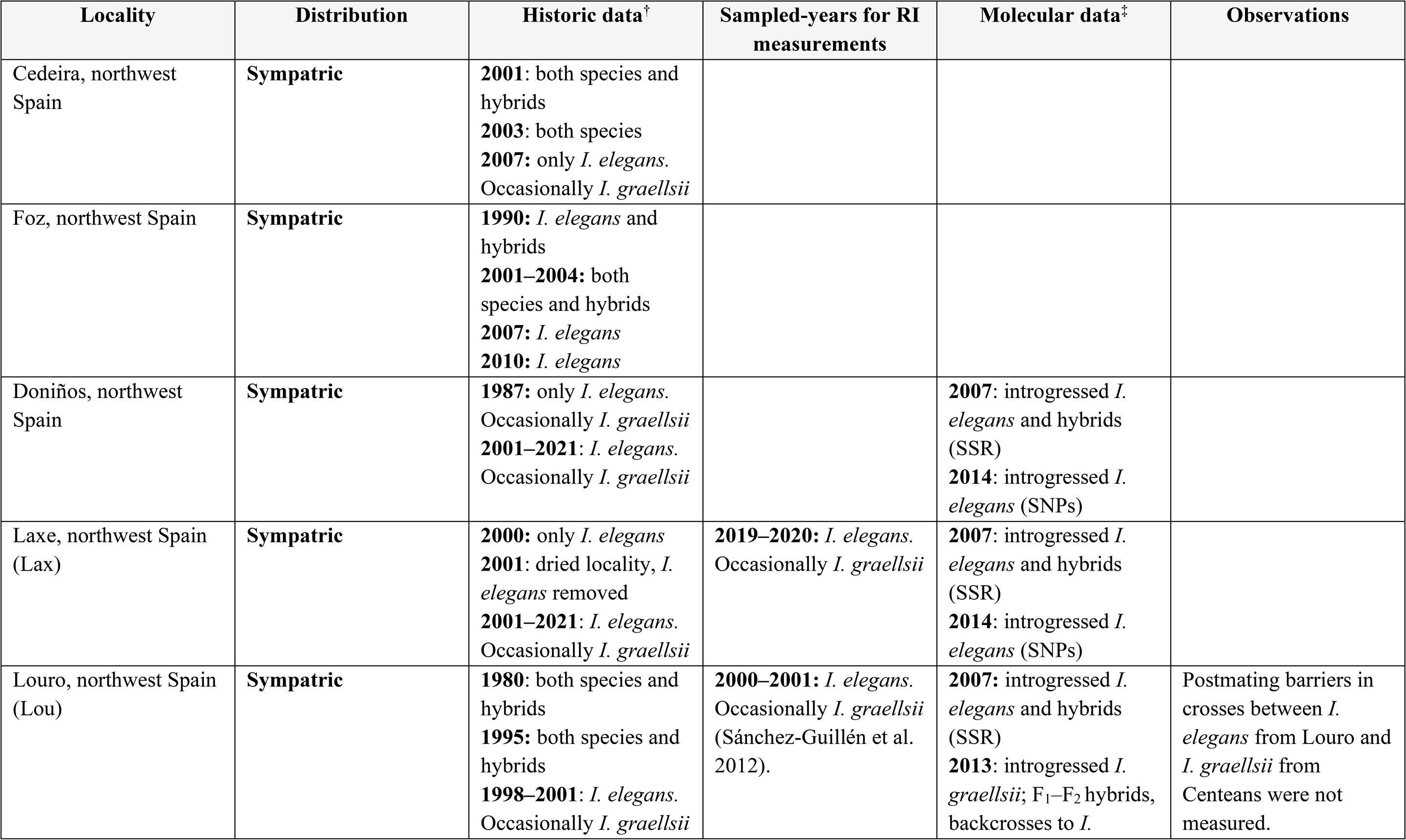
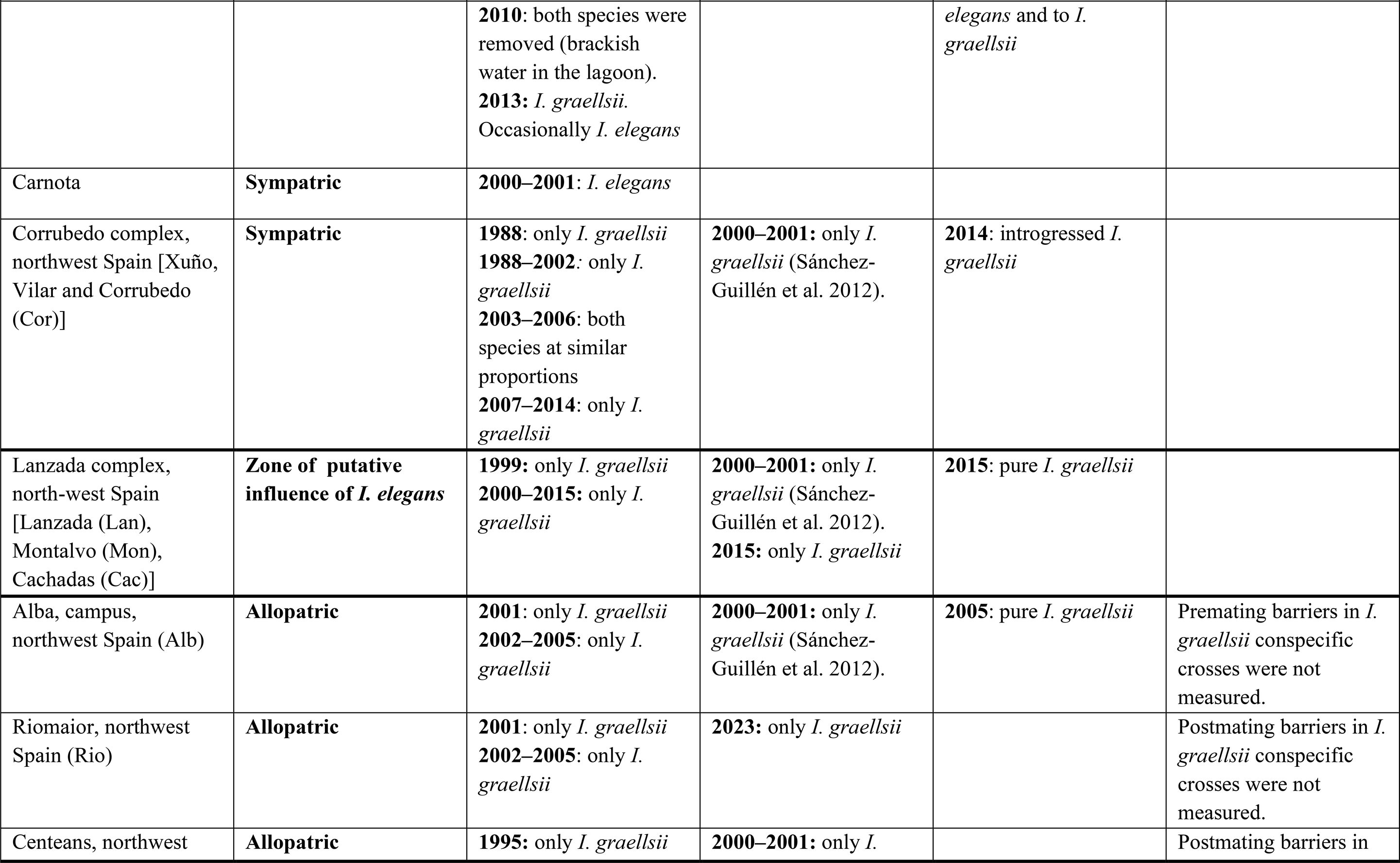
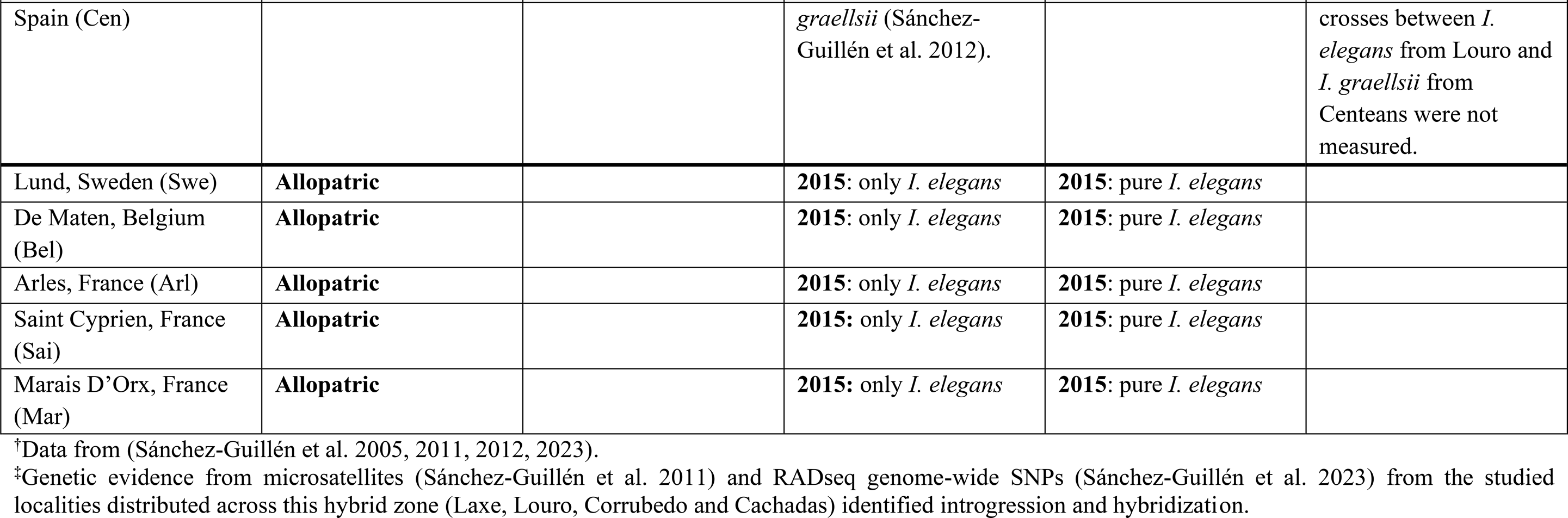
Historical and molecular data from the sampled localities of *Ischnura* damselflies.

**Table S2.**
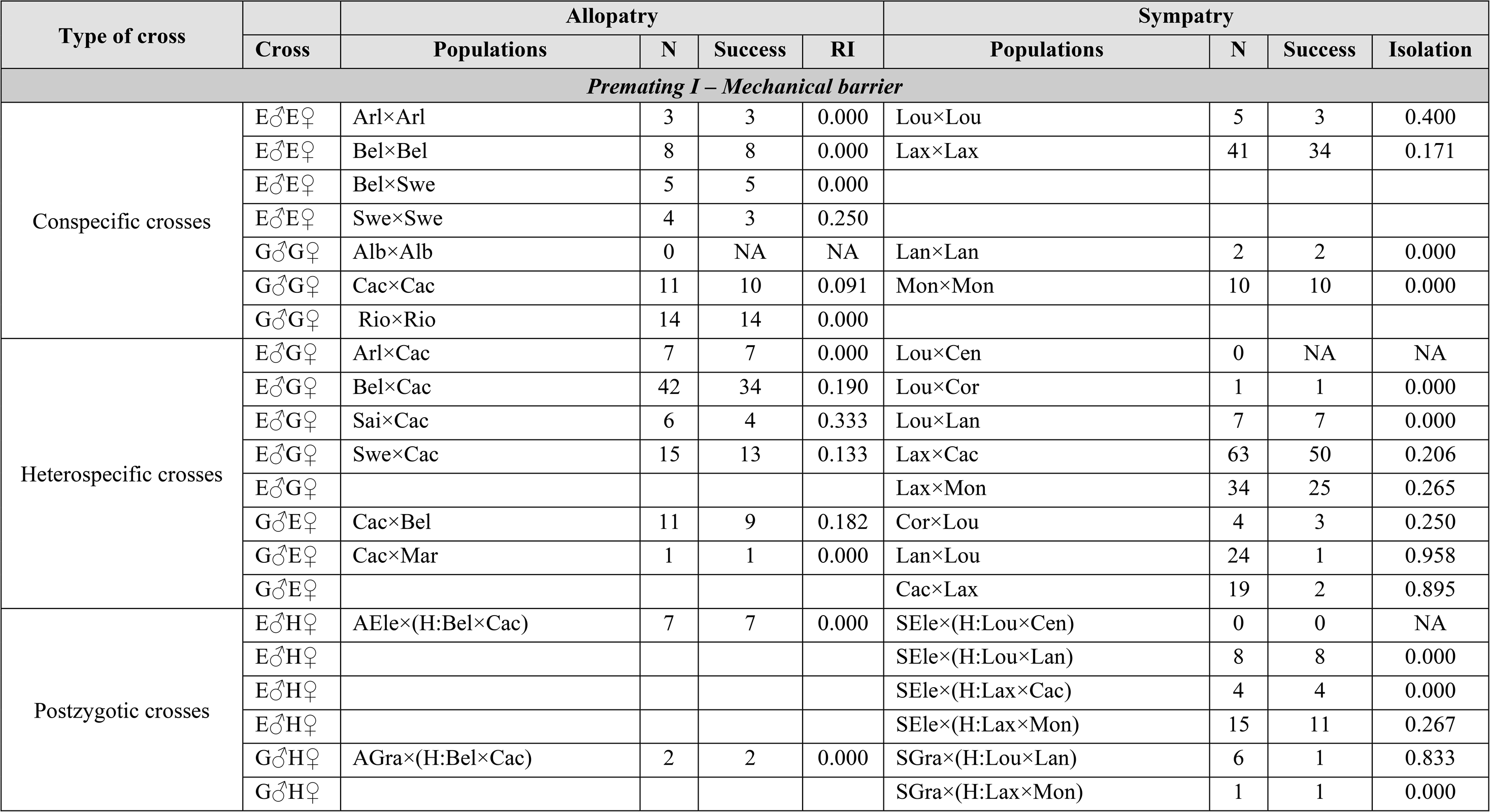
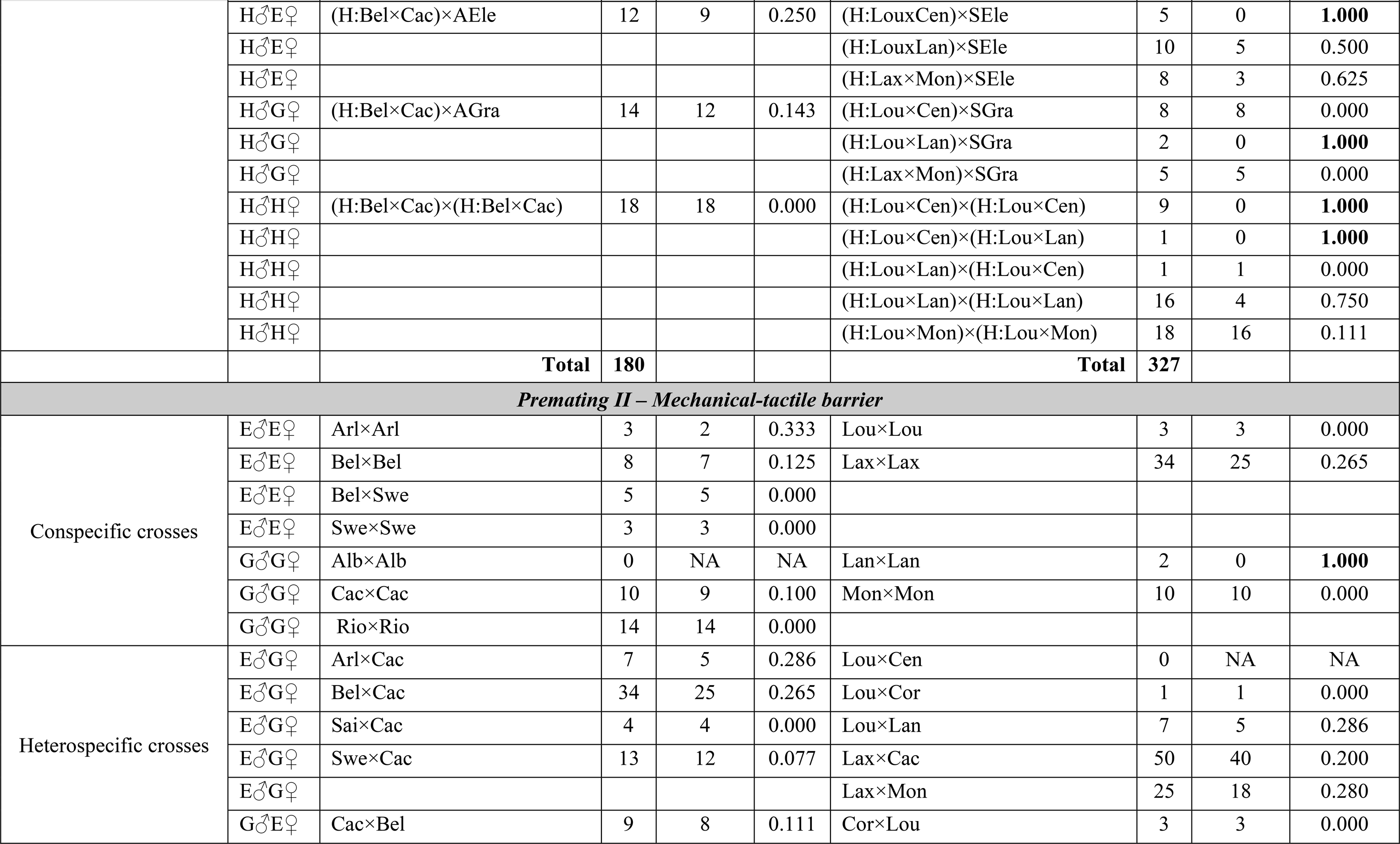
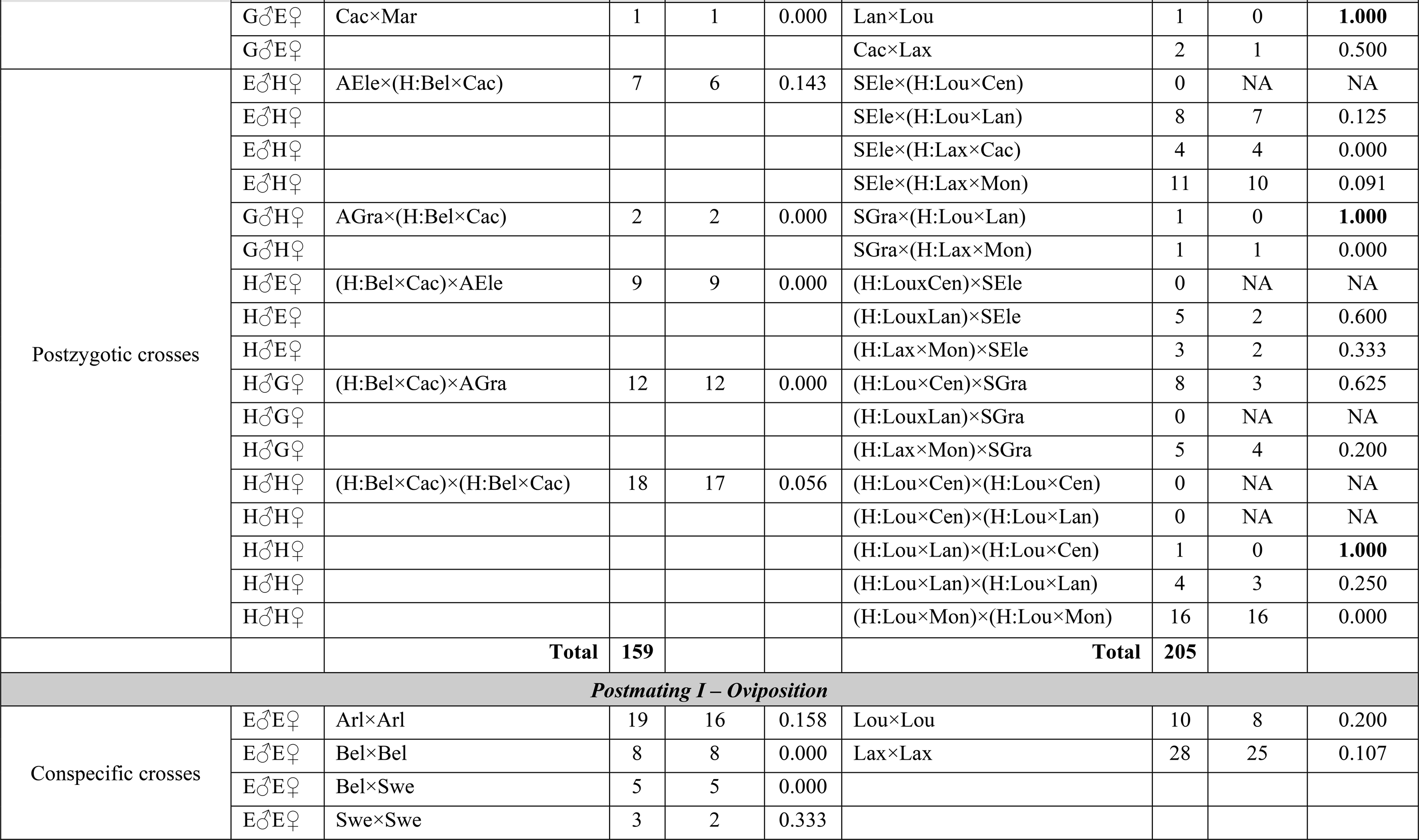
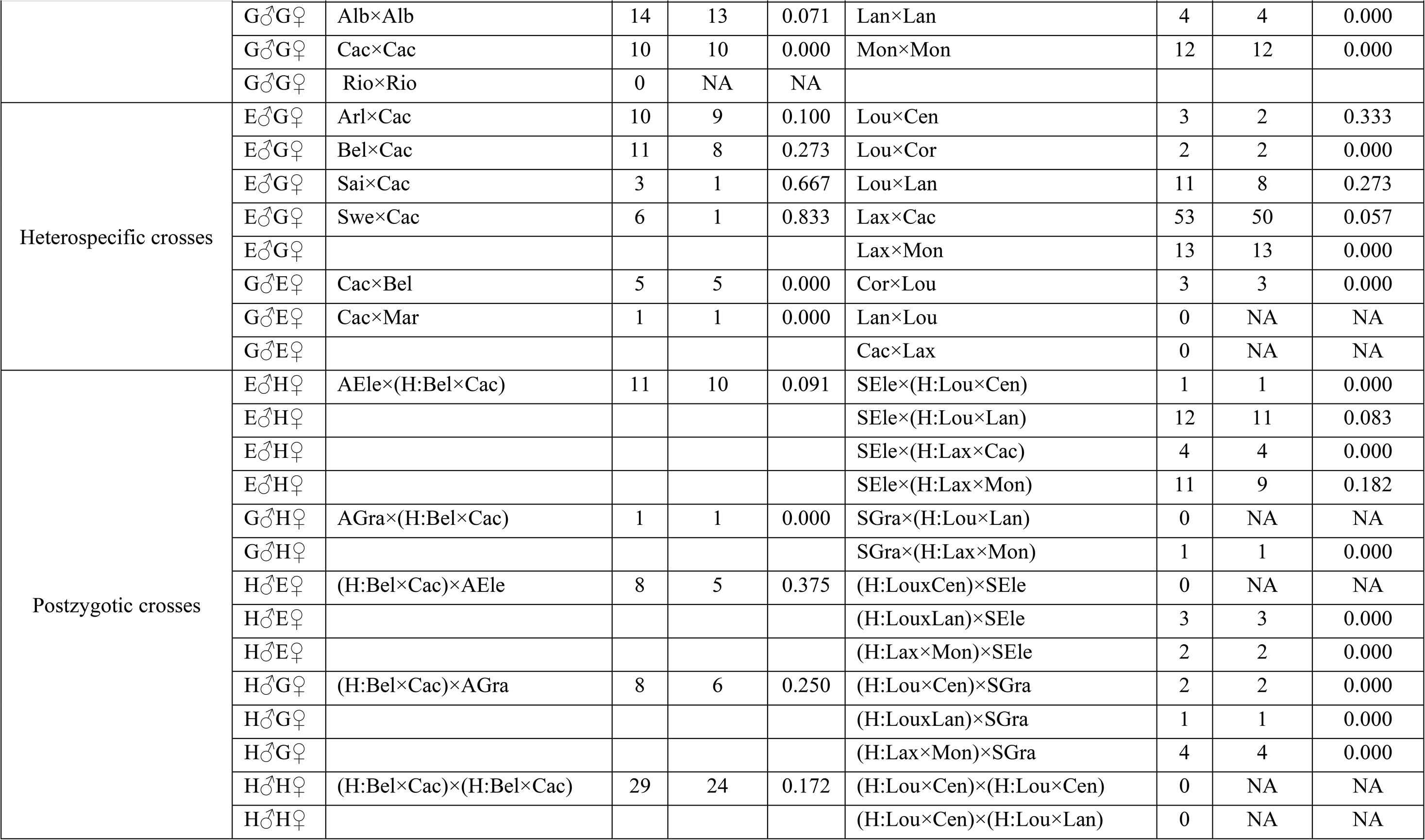
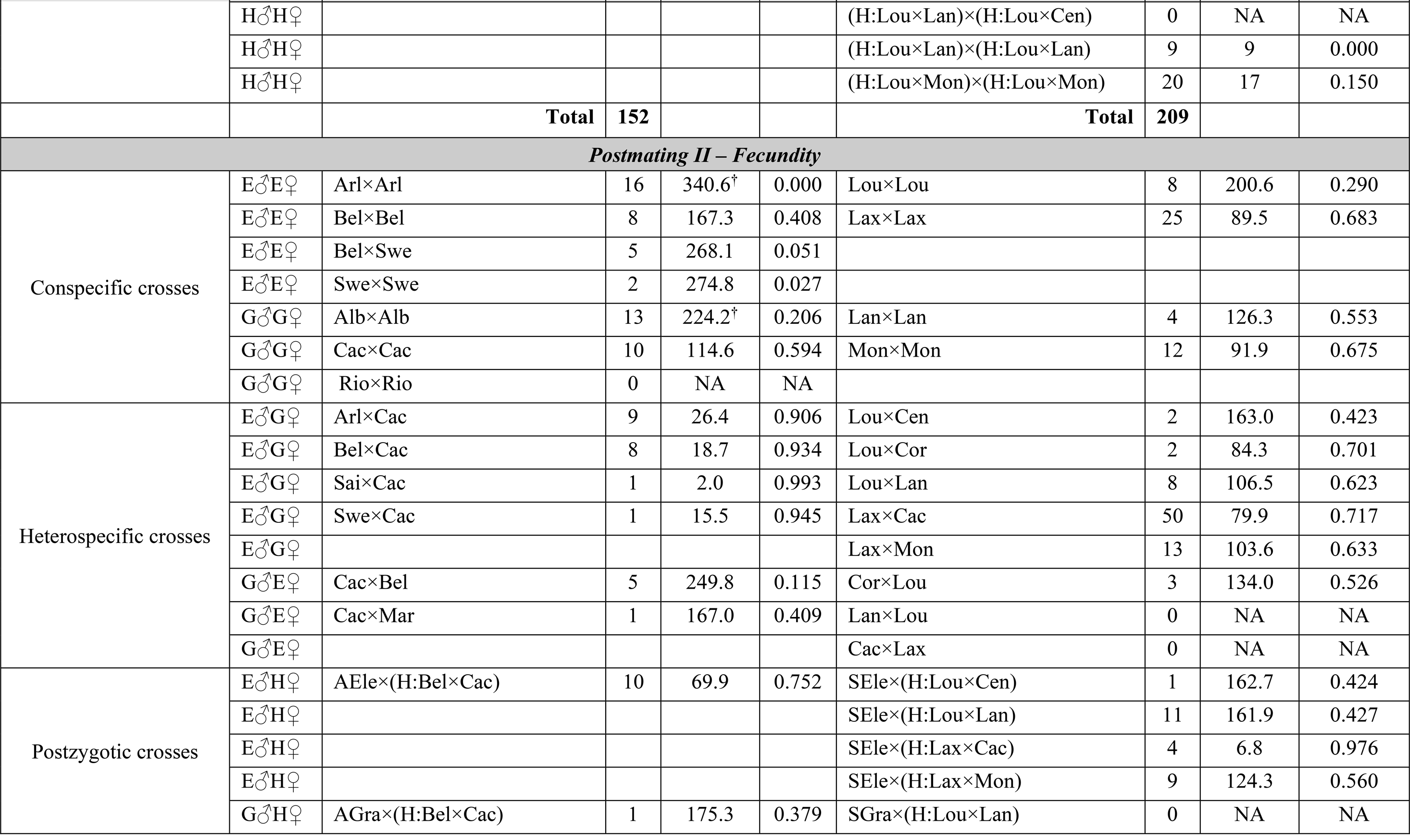
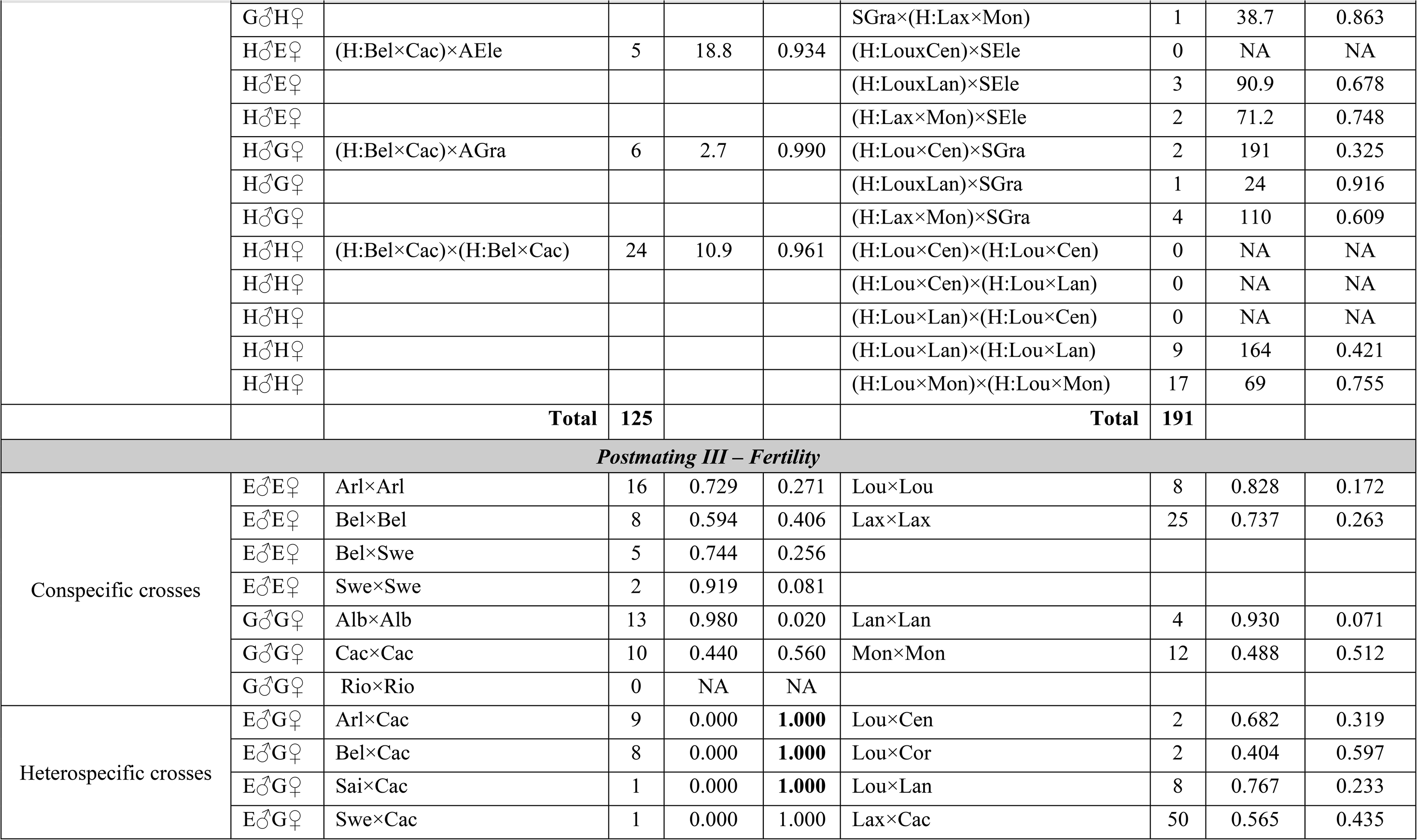
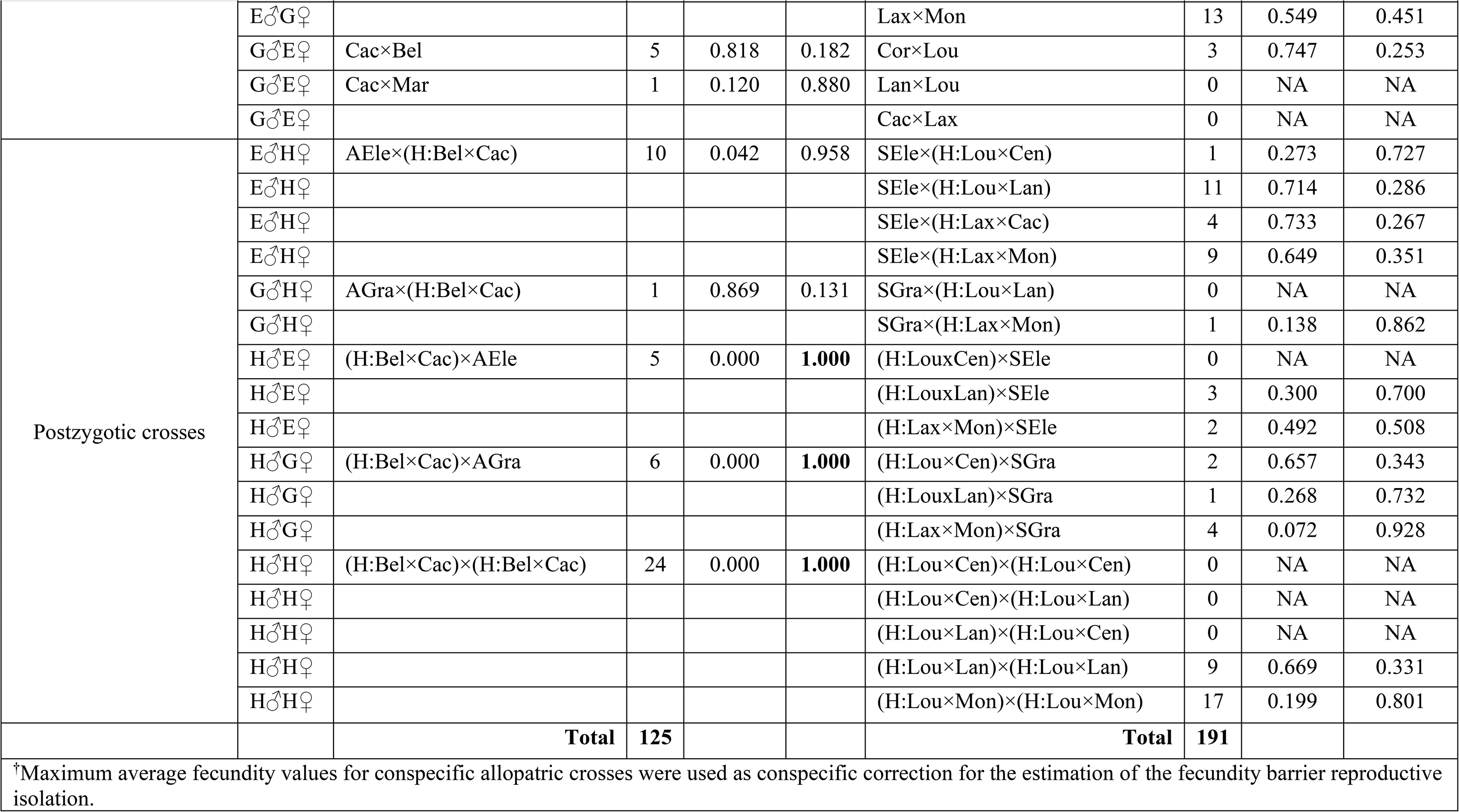
Sample size (a single interaction was recorded for each male and female *Ischnura* couple for premating barriers, while sample sizes refer to number of females for postmating barriers), success events and absolute reproductive isolation per cross between populations. G = *Ischnura graellsii;* E = *I. elegans.* Population labels are explained in the Table S1 (AEle = Pooled pure allopatric *I. elegans* samples; Agra = Pooled pure allopatric *I. graellsii* samples; SEle = Pooled pure sympatric *I. elegans* samples; SGra = Pooled pure sympatric *I. graellsii* samples).

| Type of cross |  | Allopatry |  |  |  | Sympatry |  |  |  |
| --- | --- | --- | --- | --- | --- | --- | --- | --- | --- |
|  | Cross | Populations | N | Success | RI | Populations | N | Success | Isolation |
| *Premating I – Mechanical barrier* |  |  |  |  |  |  |  |  |  |
| Conspecific crosses | E♂E♀ | Arl×Arl | 3 | 3 | 0.000 | Lou×Lou | 5 | 3 | 0.400 |
|  | E♂E♀ | Bel×Bel | 8 | 8 | 0.000 | Lax×Lax | 41 | 34 | 0.171 |
|  | E♂E♀ | Bel×Swe | 5 | 5 | 0.000 |  |  |  |  |
|  | E♂E♀ | Swe×Swe | 4 | 3 | 0.250 |  |  |  |  |
|  | G♂G♀ | Alb×Alb | 0 | NA | NA | Lan×Lan | 2 | 2 | 0.000 |
|  | G♂G♀ | Cac×Cac | 11 | 10 | 0.091 | Mon×Mon | 10 | 10 | 0.000 |
|  | G♂G♀ | Rio×Rio | 14 | 14 | 0.000 |  |  |  |  |
| Heterospecific crosses | E♂G♀ | Arl×Cac | 7 | 7 | 0.000 | Lou×Cen | 0 | NA | NA |
|  | E♂G♀ | Bel×Cac | 42 | 34 | 0.190 | Lou×Cor | 1 | 1 | 0.000 |
|  | E♂G♀ | Sai×Cac | 6 | 4 | 0.333 | Lou×Lan | 7 | 7 | 0.000 |
|  | E♂G♀ | Swe×Cac | 15 | 13 | 0.133 | Lax×Cac | 63 | 50 | 0.206 |
|  | E♂G♀ |  |  |  |  | Lax×Mon | 34 | 25 | 0.265 |
|  | G♂E♀ | Cac×Bel | 11 | 9 | 0.182 | Cor×Lou | 4 | 3 | 0.250 |
|  | G♂E♀ | Cac×Mar | 1 | 1 | 0.000 | Lan×Lou | 24 | 1 | 0.958 |
|  | G♂E♀ |  |  |  |  | Cac×Lax | 19 | 2 | 0.895 |
| Postzygotic crosses | E♂H♀ | AEle×(H:Bel×Cac) | 7 | 7 | 0.000 | SEle×(H:Lou×Cen) | 0 | 0 | NA |
|  | E♂H♀ |  |  |  |  | SEle×(H:Lou×Lan) | 8 | 8 | 0.000 |
|  | E♂H♀ |  |  |  |  | SEle×(H:Lax×Cac) | 4 | 4 | 0.000 |
|  | E♂H♀ |  |  |  |  | SEle×(H:Lax×Mon) | 15 | 11 | 0.267 |
|  | G♂H♀ | AGra×(H:Bel×Cac) | 2 | 2 | 0.000 | SGra×(H:Lou×Lan) | 6 | 1 | 0.833 |
|  | G♂H♀ |  |  |  |  | SGra×(H:Lax×Mon) | 1 | 1 | 0.000 |
|  | Cross | Populations | N | Success | RI | Populations | N | Success | Isolation |
|  | H♂E♀ | (H:Bel×Cac)×AEle | 12 | 9 | 0.250 | (H:LouxCen)×SEle | 5 | 0 | **1.000** |
|  | H♂E♀ |  |  |  |  | (H:LouxLan)×SEle | 10 | 5 | 0.500 |
|  | H♂E♀ |  |  |  |  | (H:Lax×Mon)×SEle | 8 | 3 | 0.625 |
|  | H♂G♀ | (H:Bel×Cac)×AGra | 14 | 12 | 0.143 | (H:Lou×Cen)×SGra | 8 | 8 | 0.000 |
|  | H♂G♀ |  |  |  |  | (H:Lou×Lan)×SGra | 2 | 0 | **1.000** |
|  | H♂G♀ |  |  |  |  | (H:Lax×Mon)×SGra | 5 | 5 | 0.000 |
|  | H♂H♀ | (H:Bel×Cac)×(H:Bel×Cac) | 18 | 18 | 0.000 | (H:Lou×Cen)×(H:Lou×Cen) | 9 | 0 | **1.000** |
|  | H♂H♀ |  |  |  |  | (H:Lou×Cen)×(H:Lou×Lan) | 1 | 0 | **1.000** |
|  | H♂H♀ |  |  |  |  | (H:Lou×Lan)×(H:Lou×Cen) | 1 | 1 | 0.000 |
|  | H♂H♀ |  |  |  |  | (H:Lou×Lan)×(H:Lou×Lan) | 16 | 4 | 0.750 |
|  | H♂H♀ |  |  |  |  | (H:Lou×Mon)×(H:Lou×Mon) | 18 | 16 | 0.111 |
|  |  | **Total** | **180** |  |  | **Total** | **327** |  |  |
| **Premating II – Mechanical-tactile barrier** |  |  |  |  |  |  |  |  |  |
| Conspecific crosses | E♂E♀ | Arl×Arl | 3 | 2 | 0.333 | Lou×Lou | 3 | 3 | 0.000 |
|  | E♂E♀ | Bel×Bel | 8 | 7 | 0.125 | Lax×Lax | 34 | 25 | 0.265 |
|  | E♂E♀ | Bel×Swe | 5 | 5 | 0.000 |  |  |  |  |
|  | E♂E♀ | Swe×Swe | 3 | 3 | 0.000 |  |  |  |  |
|  | G♂G♀ | Alb×Alb | 0 | NA | NA | Lan×Lan | 2 | 0 | **1.000** |
|  | G♂G♀ | Cac×Cac | 10 | 9 | 0.100 | Mon×Mon | 10 | 10 | 0.000 |
|  | G♂G♀ | Rio×Rio | 14 | 14 | 0.000 |  |  |  |  |
| Heterospecific crosses | E♂G♀ | Arl×Cac | 7 | 5 | 0.286 | Lou×Cen | 0 | NA | NA |
|  | E♂G♀ | Bel×Cac | 34 | 25 | 0.265 | Lou×Cor | 1 | 1 | 0.000 |
|  | E♂G♀ | Sai×Cac | 4 | 4 | 0.000 | Lou×Lan | 7 | 5 | 0.286 |
|  | E♂G♀ | Swe×Cac | 13 | 12 | 0.077 | Lax×Cac | 50 | 40 | 0.200 |
|  | E♂G♀ |  |  |  |  | Lax×Mon | 25 | 18 | 0.280 |
|  | G♂E♀ | Cac×Bel | 9 | 8 | 0.111 | Cor×Lou | 3 | 3 | 0.000 |

| Type of cross | Cross | Allopatry |  |  |  | Sympatry |  |  |  |
| --- | --- | --- | --- | --- | --- | --- | --- | --- | --- |
|  |  | Populations | N | Success | RI | Populations | N | Success | Isolation |
|  | G♂E♀ | Cac×Mar | 1 | 1 | 0.000 | Lan×Lou | 1 | 0 | **1.000** |
|  | G♂E♀ |  |  |  |  | Cac×Lax | 2 | 1 | 0.500 |
| Postzygotic crosses | E♂H♀ | AEle×(H:Bel×Cac) | 7 | 6 | 0.143 | SEle×(H:Lou×Cen) | 0 | NA | NA |
|  | E♂H♀ |  |  |  |  | SEle×(H:Lou×Lan) | 8 | 7 | 0.125 |
|  | E♂H♀ |  |  |  |  | SEle×(H:Lax×Cac) | 4 | 4 | 0.000 |
|  | E♂H♀ |  |  |  |  | SEle×(H:Lax×Mon) | 11 | 10 | 0.091 |
|  | G♂H♀ | AGra×(H:Bel×Cac) | 2 | 2 | 0.000 | SGra×(H:Lou×Lan) | 1 | 0 | **1.000** |
|  | G♂H♀ |  |  |  |  | SGra×(H:Lax×Mon) | 1 | 1 | 0.000 |
|  | H♂E♀ | (H:Bel×Cac)×AEle | 9 | 9 | 0.000 | (H:LouxCen)×SEle | 0 | NA | NA |
|  | H♂E♀ |  |  |  |  | (H:LouxLan)×SEle | 5 | 2 | 0.600 |
|  | H♂E♀ |  |  |  |  | (H:Lax×Mon)×SEle | 3 | 2 | 0.333 |
|  | H♂G♀ | (H:Bel×Cac)×AGra | 12 | 12 | 0.000 | (H:Lou×Cen)×SGra | 8 | 3 | 0.625 |
|  | H♂G♀ |  |  |  |  | (H:LouxLan)×SGra | 0 | NA | NA |
|  | H♂G♀ |  |  |  |  | (H:Lax×Mon)×SGra | 5 | 4 | 0.200 |
|  | H♂H♀ | (H:Bel×Cac)×(H:Bel×Cac) | 18 | 17 | 0.056 | (H:Lou×Cen)×(H:Lou×Cen) | 0 | NA | NA |
|  | H♂H♀ |  |  |  |  | (H:Lou×Cen)×(H:Lou×Lan) | 0 | NA | NA |
|  | H♂H♀ |  |  |  |  | (H:Lou×Lan)×(H:Lou×Cen) | 1 | 0 | **1.000** |
|  | H♂H♀ |  |  |  |  | (H:Lou×Lan)×(H:Lou×Lan) | 4 | 3 | 0.250 |
|  | H♂H♀ |  |  |  |  | (H:Lou×Mon)×(H:Lou×Mon) | 16 | 16 | 0.000 |
|  |  | **Total** | **159** |  |  | **Total** | **205** |  |  |
| **Postmating I – Oviposition** |  |  |  |  |  |  |  |  |  |
| Conspecific crosses | E♂E♀ | Arl×Arl | 19 | 16 | 0.158 | Lou×Lou | 10 | 8 | 0.200 |
|  | E♂E♀ | Bel×Bel | 8 | 8 | 0.000 | Lax×Lax | 28 | 25 | 0.107 |
|  | E♂E♀ | Bel×Swe | 5 | 5 | 0.000 |  |  |  |  |
|  | E♂E♀ | Swe×Swe | 3 | 2 | 0.333 |  |  |  |  |
|  |  | Populations | N | Success | RI | Populations | N | Success | Isolation |
|  | G♂G♀ | Alb×Alb | 14 | 13 | 0.071 | Lan×Lan | 4 | 4 | 0.000 |
|  | G♂G♀ | Cac×Cac | 10 | 10 | 0.000 | Mon×Mon | 12 | 12 | 0.000 |
|  | G♂G♀ | Rio×Rio | 0 | NA | NA |  |  |  |  |
| Heterospecific crosses | E♂G♀ | Arl×Cac | 10 | 9 | 0.100 | Lou×Cen | 3 | 2 | 0.333 |
|  | E♂G♀ | Bel×Cac | 11 | 8 | 0.273 | Lou×Cor | 2 | 2 | 0.000 |
|  | E♂G♀ | Sai×Cac | 3 | 1 | 0.667 | Lou×Lan | 11 | 8 | 0.273 |
|  | E♂G♀ | Swe×Cac | 6 | 1 | 0.833 | Lax×Cac | 53 | 50 | 0.057 |
|  | E♂G♀ |  |  |  |  | Lax×Mon | 13 | 13 | 0.000 |
|  | G♂E♀ | Cac×Bel | 5 | 5 | 0.000 | Cor×Lou | 3 | 3 | 0.000 |
|  | G♂E♀ | Cac×Mar | 1 | 1 | 0.000 | Lan×Lou | 0 | NA | NA |
|  | G♂E♀ |  |  |  |  | Cac×Lax | 0 | NA | NA |
| Postzygotic crosses | E♂H♀ | AEle×(H:Bel×Cac) | 11 | 10 | 0.091 | SEle×(H:Lou×Cen) | 1 | 1 | 0.000 |
|  | E♂H♀ |  |  |  |  | SEle×(H:Lou×Lan) | 12 | 11 | 0.083 |
|  | E♂H♀ |  |  |  |  | SEle×(H:Lax×Cac) | 4 | 4 | 0.000 |
|  | E♂H♀ |  |  |  |  | SEle×(H:Lax×Mon) | 11 | 9 | 0.182 |
|  | G♂H♀ | AGra×(H:Bel×Cac) | 1 | 1 | 0.000 | SGra×(H:Lou×Lan) | 0 | NA | NA |
|  | G♂H♀ |  |  |  |  | SGra×(H:Lax×Mon) | 1 | 1 | 0.000 |
|  | H♂E♀ | (H:Bel×Cac)×AEle | 8 | 5 | 0.375 | (H:LouxCen)×SEle | 0 | NA | NA |
|  | H♂E♀ |  |  |  |  | (H:LouxLan)×SEle | 3 | 3 | 0.000 |
|  | H♂E♀ |  |  |  |  | (H:Lax×Mon)×SEle | 2 | 2 | 0.000 |
|  | H♂G♀ | (H:Bel×Cac)×AGra | 8 | 6 | 0.250 | (H:Lou×Cen)×SGra | 2 | 2 | 0.000 |
|  | H♂G♀ |  |  |  |  | (H:LouxLan)×SGra | 1 | 1 | 0.000 |
|  | H♂G♀ |  |  |  |  | (H:Lax×Mon)×SGra | 4 | 4 | 0.000 |
|  | H♂H♀ | (H:Bel×Cac)×(H:Bel×Cac) | 29 | 24 | 0.172 | (H:Lou×Cen)×(H:Lou×Cen) | 0 | NA | NA |
|  | H♂H♀ |  |  |  |  | (H:Lou×Cen)×(H:Lou×Lan) | 0 | NA | NA |
|  | Cross | Populations | N | Success | RI | Populations | N | Success | Isolation |
|  | H♂H♀ |  |  |  |  | (H:Lou×Lan)×(H:Lou×Cen) | 0 | NA | NA |
|  | H♂H♀ |  |  |  |  | (H:Lou×Lan)×(H:Lou×Lan) | 9 | 9 | 0.000 |
|  | H♂H♀ |  |  |  |  | (H:Lou×Mon)×(H:Lou×Mon) | 20 | 17 | 0.150 |
|  |  | Total | 152 |  |  | Total | 209 |  |  |
| *Postmating II – Fecundity* |  |  |  |  |  |  |  |  |  |
| Conspecific crosses | E♂E♀ | Arl×Arl | 16 | 340.6<sup>†</sup> | 0.000 | Lou×Lou | 8 | 200.6 | 0.290 |
|  | E♂E♀ | Bel×Bel | 8 | 167.3 | 0.408 | Lax×Lax | 25 | 89.5 | 0.683 |
|  | E♂E♀ | Bel×Swe | 5 | 268.1 | 0.051 |  |  |  |  |
|  | E♂E♀ | Swe×Swe | 2 | 274.8 | 0.027 |  |  |  |  |
|  | G♂G♀ | Alb×Alb | 13 | 224.2<sup>†</sup> | 0.206 | Lan×Lan | 4 | 126.3 | 0.553 |
|  | G♂G♀ | Cac×Cac | 10 | 114.6 | 0.594 | Mon×Mon | 12 | 91.9 | 0.675 |
|  | G♂G♀ | Rio×Rio | 0 | NA | NA |  |  |  |  |
| Heterospecific crosses | E♂G♀ | Arl×Cac | 9 | 26.4 | 0.906 | Lou×Cen | 2 | 163.0 | 0.423 |
|  | E♂G♀ | Bel×Cac | 8 | 18.7 | 0.934 | Lou×Cor | 2 | 84.3 | 0.701 |
|  | E♂G♀ | Sai×Cac | 1 | 2.0 | 0.993 | Lou×Lan | 8 | 106.5 | 0.623 |
|  | E♂G♀ | Swe×Cac | 1 | 15.5 | 0.945 | Lax×Cac | 50 | 79.9 | 0.717 |
|  | E♂G♀ |  |  |  |  | Lax×Mon | 13 | 103.6 | 0.633 |
|  | G♂E♀ | Cac×Bel | 5 | 249.8 | 0.115 | Cor×Lou | 3 | 134.0 | 0.526 |
|  | G♂E♀ | Cac×Mar | 1 | 167.0 | 0.409 | Lan×Lou | 0 | NA | NA |
|  | G♂E♀ |  |  |  |  | Cac×Lax | 0 | NA | NA |
| Postzygotic crosses | E♂H♀ | AEle×(H:Bel×Cac) | 10 | 69.9 | 0.752 | SEle×(H:Lou×Cen) | 1 | 162.7 | 0.424 |
|  | E♂H♀ |  |  |  |  | SEle×(H:Lou×Lan) | 11 | 161.9 | 0.427 |
|  | E♂H♀ |  |  |  |  | SEle×(H:Lax×Cac) | 4 | 6.8 | 0.976 |
|  | E♂H♀ |  |  |  |  | SEle×(H:Lax×Mon) | 9 | 124.3 | 0.560 |
|  | G♂H♀ | AGra×(H:Bel×Cac) | 1 | 175.3 | 0.379 | SGra×(H:Lou×Lan) | 0 | NA | NA |
|  | Cross | Populations | N | Success | RI | Populations | N | Success | Isolation |
|  | G♂H♀ |  |  |  |  | SGra×(H:Lax×Mon) | 1 | 38.7 | 0.863 |
|  | H♂E♀ | (H:Bel×Cac)×AEle | 5 | 18.8 | 0.934 | (H:LouxCen)×SEle | 0 | NA | NA |
|  | H♂E♀ |  |  |  |  | (H:LouxLan)×SEle | 3 | 90.9 | 0.678 |
|  | H♂E♀ |  |  |  |  | (H:Lax×Mon)×SEle | 2 | 71.2 | 0.748 |
|  | H♂G♀ | (H:Bel×Cac)×AGra | 6 | 2.7 | 0.990 | (H:Lou×Cen)×SGra | 2 | 191 | 0.325 |
|  | H♂G♀ |  |  |  |  | (H:LouxLan)×SGra | 1 | 24 | 0.916 |
|  | H♂G♀ |  |  |  |  | (H:Lax×Mon)×SGra | 4 | 110 | 0.609 |
|  | H♂H♀ | (H:Bel×Cac)×(H:Bel×Cac) | 24 | 10.9 | 0.961 | (H:Lou×Cen)×(H:Lou×Cen) | 0 | NA | NA |
|  | H♂H♀ |  |  |  |  | (H:Lou×Cen)×(H:Lou×Lan) | 0 | NA | NA |
|  | H♂H♀ |  |  |  |  | (H:Lou×Lan)×(H:Lou×Cen) | 0 | NA | NA |
|  | H♂H♀ |  |  |  |  | (H:Lou×Lan)×(H:Lou×Lan) | 9 | 164 | 0.421 |
|  | H♂H♀ |  |  |  |  | (H:Lou×Mon)×(H:Lou×Mon) | 17 | 69 | 0.755 |
|  |  | **Total** | **125** |  |  | **Total** | **191** |  |  |
| **Postmating III – Fertility** |  |  |  |  |  |  |  |  |  |
| Conspecific crosses | E♂E♀ | Arl×Arl | 16 | 0.729 | 0.271 | Lou×Lou | 8 | 0.828 | 0.172 |
|  | E♂E♀ | Bel×Bel | 8 | 0.594 | 0.406 | Lax×Lax | 25 | 0.737 | 0.263 |
|  | E♂E♀ | Bel×Swe | 5 | 0.744 | 0.256 |  |  |  |  |
|  | E♂E♀ | Swe×Swe | 2 | 0.919 | 0.081 |  |  |  |  |
|  | G♂G♀ | Alb×Alb | 13 | 0.980 | 0.020 | Lan×Lan | 4 | 0.930 | 0.071 |
|  | G♂G♀ | Cac×Cac | 10 | 0.440 | 0.560 | Mon×Mon | 12 | 0.488 | 0.512 |
|  | G♂G♀ | Rio×Rio | 0 | NA | NA |  |  |  |  |
| Heterospecific crosses | E♂G♀ | Arl×Cac | 9 | 0.000 | **1.000** | Lou×Cen | 2 | 0.682 | 0.319 |
|  | E♂G♀ | Bel×Cac | 8 | 0.000 | **1.000** | Lou×Cor | 2 | 0.404 | 0.597 |
|  | E♂G♀ | Sai×Cac | 1 | 0.000 | **1.000** | Lou×Lan | 8 | 0.767 | 0.233 |
|  | E♂G♀ | Swe×Cac | 1 | 0.000 | 1.000 | Lax×Cac | 50 | 0.565 | 0.435 |

| Type of cross | Cross | Allopatry |  |  |  | Sympatry |  |  |  |
| --- | --- | --- | --- | --- | --- | --- | --- | --- | --- |
|  |  | Populations | N | Success | RI | Populations | N | Success | Isolation |
|  | E♂G♀ |  |  |  |  | Lax×Mon | 13 | 0.549 | 0.451 |
|  | G♂E♀ | Cac×Bel | 5 | 0.818 | 0.182 | Cor×Lou | 3 | 0.747 | 0.253 |
|  | G♂E♀ | Cac×Mar | 1 | 0.120 | 0.880 | Lan×Lou | 0 | NA | NA |
|  | G♂E♀ |  |  |  |  | Cac×Lax | 0 | NA | NA |
| Postzygotic crosses | E♂H♀ | AEle×(H:Bel×Cac) | 10 | 0.042 | 0.958 | SEle×(H:Lou×Cen) | 1 | 0.273 | 0.727 |
|  | E♂H♀ |  |  |  |  | SEle×(H:Lou×Lan) | 11 | 0.714 | 0.286 |
|  | E♂H♀ |  |  |  |  | SEle×(H:Lax×Cac) | 4 | 0.733 | 0.267 |
|  | E♂H♀ |  |  |  |  | SEle×(H:Lax×Mon) | 9 | 0.649 | 0.351 |
|  | G♂H♀ | AGra×(H:Bel×Cac) | 1 | 0.869 | 0.131 | SGra×(H:Lou×Lan) | 0 | NA | NA |
|  | G♂H♀ |  |  |  |  | SGra×(H:Lax×Mon) | 1 | 0.138 | 0.862 |
|  | H♂E♀ | (H:Bel×Cac)×AEle | 5 | 0.000 | **1.000** | (H:Lou×Cen)×SEle | 0 | NA | NA |
|  | H♂E♀ |  |  |  |  | (H:Lou×Lan)×SEle | 3 | 0.300 | 0.700 |
|  | H♂E♀ |  |  |  |  | (H:Lax×Mon)×SEle | 2 | 0.492 | 0.508 |
|  | H♂G♀ | (H:Bel×Cac)×AGra | 6 | 0.000 | **1.000** | (H:Lou×Cen)×SGra | 2 | 0.657 | 0.343 |
|  | H♂G♀ |  |  |  |  | (H:Lou×Lan)×SGra | 1 | 0.268 | 0.732 |
|  | H♂G♀ |  |  |  |  | (H:Lax×Mon)×SGra | 4 | 0.072 | 0.928 |
|  | H♂H♀ | (H:Bel×Cac)×(H:Bel×Cac) | 24 | 0.000 | **1.000** | (H:Lou×Cen)×(H:Lou×Cen) | 0 | NA | NA |
|  | H♂H♀ |  |  |  |  | (H:Lou×Cen)×(H:Lou×Lan) | 0 | NA | NA |
|  | H♂H♀ |  |  |  |  | (H:Lou×Lan)×(H:Lou×Cen) | 0 | NA | NA |
|  | H♂H♀ |  |  |  |  | (H:Lou×Lan)×(H:Lou×Lan) | 9 | 0.669 | 0.331 |
|  | H♂H♀ |  |  |  |  | (H:Lou×Mon)×(H:Lou×Mon) | 17 | 0.199 | 0.801 |
|  |  | **Total** | **125** |  |  | **Total** | **191** |  |  |
| †Maximum average fecundity values for conspecific allopatric crosses were used as conspecific correction for the estimation of the fecundity barrier reproductive isolation. |  |  |  |  |  |  |  |  |  |

**Table S3.** GLM model comparisons per prezygotic reproductive barrier in *Ischnura graellsii* and *I. elegans*. Models are sorted by increasing values of the AICc. “+” signs on each parameter shows the inclusion of each parameter in each model. Cross = Types of crosses; Ecology = Sympatry vs Allopatry; Geography = Intrapopulation vs Interpopulation crosses; Cross:Ecology = Interaction between crosses and ecology; df = degrees freedom; logLik = log-likelihood. The model with the lowest scoring AICc per barrier was selected as the best model.

| Model | Intercept | Cross | Ecology | Geography | Cross:Ecology | df | logLik | AICc | delta | weight |
| --- | --- | --- | --- | --- | --- | --- | --- | --- | --- | --- |
| *Premating I – Mechanical barrier (Binomial distribution)* |  |  |  |  |  |  |  |  |  |  |
| 12 | 2.944 | + | + |  | + | 8 | -140.233 | 296.9 | 0 | 0.676 |
| 16 | 17.57 | + | + | + | + | 9 | -139.937 | 298.4 | 1.52 | 0.316 |
| 4 | 2.52 | + | + |  |  | 5 | -148.278 | 306.7 | 9.83 | 0.005 |
| 8 | 16.57 | + | + | + |  | 6 | -147.876 | 308 | 11.1 | 0.003 |
| 2 | 1.723 | + |  |  |  | 4 | -153.317 | 314.8 | 17.85 | 0 |
| 6 | 15.57 | + |  | + |  | 5 | -152.459 | 315.1 | 18.19 | 0 |
| 7 | 1.645 |  | + | + |  | 3 | -174.481 | 355 | 58.13 | 0 |
| 3 | 1.937 |  | + |  |  | 2 | -183.104 | 370.2 | 73.34 | 0 |
| 5 | 0.7438 |  |  | + |  | 2 | -184.627 | 373.3 | 76.38 | 0 |
| 1 | 1.04 |  |  |  |  | 1 | -193.517 | 389 | 92.14 | 0 |
| *Premating II – Mechanical-tactile barrier (Binomial distribution)* |  |  |  |  |  |  |  |  |  |  |
| 3 | 1.781 |  | + |  |  | 2 | -120.511 | 245.1 | 0 | 0.299 |
| 1 | 1.432 |  |  |  |  | 1 | -122.064 | 246.1 | 1.07 | 0.175 |
| 7 | 1.675 |  | + | + |  | 3 | -120.038 | 246.2 | 1.1 | 0.172 |
| 5 | 1.326 |  |  | + |  | 2 | -121.618 | 247.3 | 2.21 | 0.099 |
| 6 | 16.57 | + |  | + |  | 5 | -118.907 | 248.1 | 2.99 | 0.067 |
| 4 | 1.754 | + | + |  |  | 5 | -118.987 | 248.2 | 3.15 | 0.062 |
| 2 | 1.409 | + |  |  |  | 4 | -120.059 | 248.3 | 3.21 | 0.06 |
| 8 | 16.57 | + | + | + |  | 6 | -118.136 | 248.6 | 3.55 | 0.051 |
| 12 | 2.14 | + | + |  | + | 8 | -117.774 | 252.1 | 7.08 | 0.009 |
| 16 | 16.57 | + | + | + | + | 9 | -117.123 | 253 | 7.93 | 0.006 |
| *Postmating I – Oviposition (Binomial distribution)* |  |  |  |  |  |  |  |  |  |  |
| 8 | 17.57 | + | + | + |  | 6 | -76.568 | 165.5 | 0 | 0.32 |
| 4 | 1.483 | + | + |  |  | 5 | -77.661 | 165.6 | 0.08 | 0.308 |
| 12 | 2.048 | + | + |  | + | 8 | -75.024 | 166.7 | 1.18 | 0.177 |
| 16 | 18.57 | + | + | + | + | 9 | -74.366 | 167.5 | 2.03 | 0.116 |
| 2 | 1.962 | + |  |  |  | 4 | -81.312 | 170.8 | 5.29 | 0.023 |
| 6 | 17.57 | + |  | + |  | 5 | -80.629 | 171.5 | 6.01 | 0.016 |
| 7 | 1.257 |  | + | + |  | 3 | -82.708 | 171.5 | 6.01 | 0.016 |
| 3 | 1.597 |  | + |  |  | 2 | -83.932 | 171.9 | 6.41 | 0.013 |
| 1 | 1.996 |  |  |  |  | 1 | -85.701 | 173.4 | 7.91 | 0.006 |
| 5 | 1.792 |  |  | + |  | 2 | -84.992 | 174 | 8.53 | 0.005 |
| *Postmating II – Fecundity (Poisson distribution)* |  |  |  |  |  |  |  |  |  |  |
| 16 | 5.592 | + | + | + | + | 9 | -7780.176 | 15579.3 | 0 | 0.586 |
| 12 | 5.635 | + | + |  | + | 8 | -7781.616 | 15580 | 0.7 | 0.414 |
| 8 | 5.592 | + | + | + |  | 6 | -9300.452 | 18613.3 | 3034.06 | 0 |
| 4 | 5.456 | + | + |  |  | 5 | -9314.166 | 18638.6 | 3059.37 | 0 |
| 6 | 5.592 | + |  | + |  | 5 | -9831.48 | 19673.3 | 4094 | 0 |
| 2 | 5.246 | + |  |  |  | 4 | -9910.267 | 19828.7 | 4249.47 | 0 |
| 7 | 4.912 |  | + | + |  | 3 | -10138.447 | 20283 | 4703.75 | 0 |
| 3 | 5.217 |  | + |  |  | 2 | -10782.699 | 21569.5 | 5990.2 | 0 |
| 5 | 4.553 |  |  | + |  | 2 | -11074.958 | 22154 | 6574.72 | 0 |
| 1 | 4.87 |  |  |  |  | 1 | -12160.64 | 24323.3 | 8744.04 | 0 |
| *Postmating III – Fertility (Binomial distribution)* |  |  |  |  |  |  |  |  |  |  |
| 16 | 1.785 | + | + | + | + | 9 | -10261.8 | 20542.6 | 0 | 1 |
| 12 | 1.422 | + | + |  | + | 8 | -10308.74 | 20634.3 | 91.68 | 0 |
| 8 | 1.785 | + | + | + |  | 6 | -10910.35 | 21833.2 | 1290.57 | 0 |
| 4 | 1.288 | + | + |  |  | 5 | -10997.73 | 22005.8 | 1463.2 | 0 |
| 6 | 1.785 | + |  | + |  | 5 | -11169.1 | 22348.5 | 1805.94 | 0 |
| 2 | 1.466 | + |  |  |  | 4 | -11201.49 | 22411.2 | 1868.6 | 0 |
| 7 | 0.9834 |  | + | + |  | 3 | -11342.68 | 22691.5 | 2148.9 | 0 |
| 5 | 1.092 |  |  | + |  | 2 | -11369.85 | 22743.8 | 2201.17 | 0 |
| 1 | 1.308 |  |  |  |  | 1 | -11594.82 | 23191.7 | 2649.07 | 0 |
| 3 | 1.321 |  | + |  |  | 2 | -11593.89 | 23191.8 | 2649.25 | 0 |

**Table S4.** *Post hoc* GLM modeling for reproductive isolation as a function of types of crosses (RI ∼ Cross) per prezygotic reproductive barrier in *Ischnura graellsii* and *I. elegans*. Mechanical-tactile barrier was excluded as crosses were not a significant parameter in its GLM modeling (Table S3). GLMs were modeled using each cross direction as model intercept to allow pairwise comparisons between types of crosses. S.E. = Standard error; * = Significant p value for differences between a cross and the model intercept (p<0.05/6).

| Barrier | Cross | Estimate | S.E. | z value | p | p<0.05/6 |
| --- | --- | --- | --- | --- | --- | --- |
| Mechanical | **Intercept: elegans♂×graellsii♀** |  |  |  |  |  |
|  | **Intercept** | 1.4224 | 0.1911 | 7.45E+00 | 9.71E-14 | NA |
|  | elegans♂×elegans♀ | 0.3004 | 0.3929 | 0.765 | 0.4446 |  |
|  | graellsii♂×graellsii♀ | 2.1611 | 1.0316 | 2.095 | 0.0362 |  |
|  | graellsii♂×elegans♀ | -2.411 | 0.3497 | -6.90E+00 | 5.37E-12 | * |
|  | **Intercept: graellsii♂×graellsii♀** |  |  |  |  |  |
|  | **Intercept** | 3.584 | 1.014 | 3.535 | 0.000408 | NA |
|  | elegans♂×graellsii♀ | -2.161 | 1.032 | -2.095 | 0.036184 |  |
|  | elegans♂×elegans♀ | -1.861 | 1.07 | -1.738 | 0.082128 |  |
|  | graellsii♂×elegans♀ | -4.572 | 1.055 | -4.33E+00 | 1.47E-05 | * |
|  | **Intercept: elegans♂×elegans♀** |  |  |  |  |  |
|  | **Intercept** | 1.7228 | 0.3433 | 5.02E+00 | 5.22E-07 | NA |
|  | graellsii♂×graellsii♀ | 1.8608 | 1.0703 | 1.738 | 0.0821 |  |
|  | elegans♂×graellsii♀ | -0.3004 | 0.3929 | -0.765 | 0.4446 |  |
|  | graellsii♂×elegans♀ | -2.7114 | 0.4512 | -6.01E+00 | 1.87E-09 | * |
|  | **Intercept: graellsii♂×elegans♀** |  |  |  |  |  |
|  | **Intercept** | -0.9886 | 0.2928 | -3.376 | 0.000736 | NA |
|  | elegans♂×elegans♀ | 2.7114 | 0.4512 | 6.01E+00 | 1.87E-09 | * |
|  | graellsii♂×graellsii♀ | 4.5721 | 1.0552 | 4.33E+00 | 1.47E-05 | * |
|  | elegans♂×graellsii♀ | 2.411 | 0.3497 | 6.90E+00 | 5.37E-12 | * |
| Oviposition | **Intercept: elegans♂×graellsii♀** |  |  |  |  |  |
|  | **Intercept** | 1.6529 | 0.2573 | 6.425 | 1.32E-10 | NA |
|  | elegans♂×elegans♀ | 0.3087 | 0.4392 | 0.703 | 0.4821 |  |
|  | graellsii♂×graellsii♀ | 2.0106 | 1.0449 | 1.924 | 0.0543 |  |
|  | graellsii♂×elegans♀ | 15.9131 | 1318.7268 | 0.012 | 0.9904 |  |
|  | **Intercept: graellsii♂×graellsii♀** |  |  |  |  |  |
|  | **Intercept** | 3.664 | 1.013 | 3.617 | 0.000297 | NA |
|  | elegans♂×graellsii♀ | -2.011 | 1.045 | -1.924 | 0.054327 |  |
|  | elegans♂×elegans♀ | -1.702 | 1.073 | -1.585 | 0.112877 |  |
|  | graellsii♂×elegans♀ | 13.903 | 1318.727 | 0.011 | 0.991589 |  |
|  | **Intercept: elegans♂×elegans♀** |  |  |  |  |  |
|  | **Intercept** | 1.9617 | 0.356 | 5.51 | 3.58E-08 | NA |
|  | graellsii♂×graellsii♀ | 1.7019 | 1.0735 | 1.585 | 0.113 |  |
|  | elegans♂×graellsii♀ | -0.3087 | 0.4392 | -0.703 | 0.482 |  |
|  | graellsii♂×elegans♀ | 15.6044 | 1318.7268 | 0.012 | 0.991 |  |
|  | **Intercept: graellsii♂×elegans♀** |  |  |  |  |  |
|  | **Intercept** | 17.57 | 1318.73 | 0.013 | 0.989 |  |
|  | elegans♂×elegans♀ | -15.6 | 1318.73 | -0.012 | 0.991 |  |
|  | graellsii♂×graellsii♀ | -13.9 | 1318.73 | -0.011 | 0.992 |  |
|  | elegans♂×graellsii♀ | -15.91 | 1318.73 | -0.012 | 0.99 |  |
| **Fecundity** | **Intercept: elegans♂×graellsii♀** |  |  |  |  |  |
|  | **Intercept** | 4.32343 | 0.01187 | 364.1 | <2E-16 | NA |
|  | elegans♂×elegans♀ | 0.92248 | 0.01486 | 62.08 | <2E-16 | * |
|  | graellsii♂×graellsii♀ | 0.65578 | 0.01782 | 36.81 | <2E-16 | * |
|  | graellsii♂×elegans♀ | 0.98539 | 0.02628 | 37.49 | <2E-16 | * |
|  | **Intercept: graellsii♂×graellsii♀** |  |  |  |  |  |
|  | **Intercept** | 4.97921 | 0.01328 | 374.9 | <2E-16 | NA |
|  | elegans♂×graellsii♀ | -0.65578 | 0.01782 | -36.81 | <2E-16 | * |
|  | elegans♂×elegans♀ | 0.2667 | 0.01601 | 16.66 | <2E-16 | * |
|  | graellsii♂×elegans♀ | 0.32961 | 0.02695 | 12.23 | <2E-16 | * |
|  | **Intercept: elegans♂×elegans♀** |  |  |  |  |  |
|  | **Intercept** | 5.245907 | 0.008935 | 587.12 | <2E-16 | NA |
|  | graellsii♂×graellsii♀ | -0.266701 | 0.016007 | -16.661 | <2E-16 | * |
|  | elegans♂×graellsii♀ | -0.922479 | 0.01486 | -62.076 | <2E-16 | * |
|  | graellsii♂×elegans♀ | 0.062911 | 0.025092 | 2.507 | 0.0122 |  |
|  | **Intercept: graellsii♂×elegans♀** |  |  |  |  |  |
|  | **Intercept** | 5.30882 | 0.02345 | 226.42 | <2E-16 | NA |
|  | elegans♂×elegans♀ | -0.06291 | 0.02509 | -2.507 | 0.0122 |  |
|  | graellsii♂×graellsii♀ | -0.32961 | 0.02695 | -12.232 | <2E-16 | * |
|  | elegans♂×graellsii♀ | -0.98539 | 0.02628 | -37.493 | <2E-16 | * |
| **Fertility** | **Intercept: elegans♂×graellsii♀** |  |  |  |  |  |
|  | **Intercept** | 0.98859 | 0.01645 | 60.082 | <2E-16 | NA |
|  | elegans♂×elegans♀ | 0.47708 | 0.02348 | 20.316 | <2E-16 | * |
|  | graellsii♂×graellsii♀ | 0.67899 | 0.02833 | 23.967 | <2E-16 | * |
|  | graellsii♂×elegans♀ | 0.04076 | 0.03572 | 1.141 | 0.254 |  |
|  | **Intercept: graellsii♂×graellsii♀** |  |  |  |  |  |
|  | **Intercept** | 1.66758 | 0.02306 | 72.309 | <2E-16 | NA |
|  | elegans♂×graellsii♀ | -0.67899 | 0.02833 | -23.967 | <2E-16 | * |
|  | elegans♂×elegans♀ | -0.20191 | 0.02851 | -7.083 | 1.41E-12 | * |
|  | graellsii♂×elegans♀ | -0.63823 | 0.03921 | -16.278 | <2E-16 | * |
|  | **Intercept: elegans♂×elegans♀** |  |  |  |  |  |
|  | **Intercept** | 1.46567 | 0.01675 | 87.479 | <2E-16 | NA |
|  | graellsii♂×graellsii♀ | 0.20191 | 0.02851 | 7.083 | 1.41E-12 | * |
|  | elegans♂×graellsii♀ | -0.47708 | 0.02348 | -20.316 | <2E-16 | * |
|  | graellsii♂×elegans♀ | -0.43632 | 0.03586 | -12.167 | <2E-16 | * |
|  | **Intercept: graellsii♂×elegans♀** |  |  |  |  |  |
|  | **Intercept** | 1.02935 | 0.03171 | 32.464 | <2E-16 | NA |
|  | elegans♂×elegans♀ | 0.43632 | 0.03586 | 12.167 | <2E-16 | * |
|  | graellsii♂×graellsii♀ | 0.63823 | 0.03921 | 16.278 | <2E-16 | * |
|  | elegans♂×graellsii♀ | -0.04076 | 0.03572 | -1.141 | 0.254 |  |

**Table S5.**
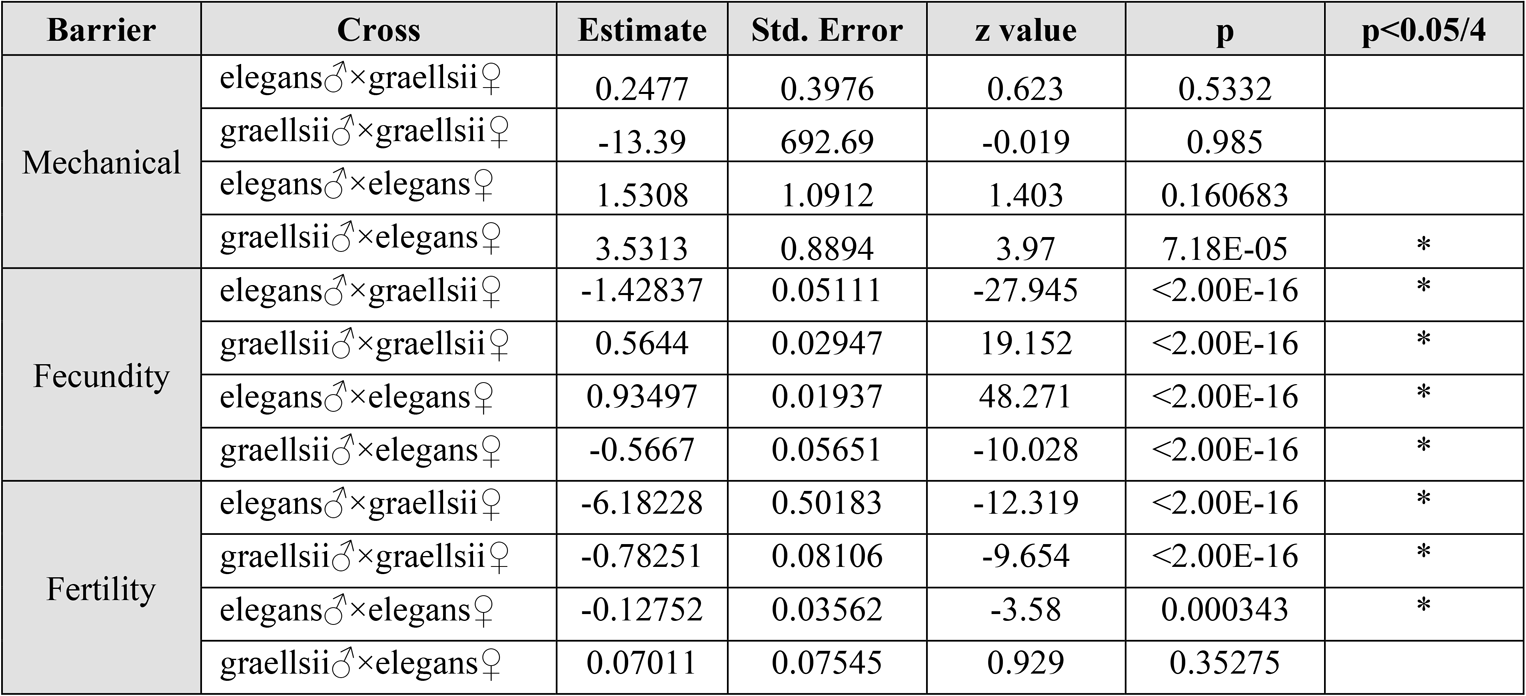
*Post hoc* GLM modeling for reproductive isolation as a function of the interaction ecology and types of crosses (RI ∼ Ecology:Cross) per prezygotic reproductive barrier in *Ischnura graellsii* and *I. elegans*. Mechanical-tactile and oviposition barriers were excluded as ecology and cross interaction were not significant parameters in its GLM modeling (Table S3). Although each cross in each ecology was compared with each other combination, here we report only results for differences between ecologies within each type of cross. * = Significant p value for differences between the allopatric and sympatric ecology (p<0.05/4).

**Table S6.** GLM models comparison per postzygotic reproductive barrier in *Ischnura graellsii* and *I. elegans*. Models are sorted by increasing values of the AICc. “+” signs on each parameter shows the inclusion of each parameter in each model. Cross = Types of crosses; Ecology = Sympatry vs Allopatry; Cross:Ecology = Interaction between crosses and ecology; df = degrees freedom; logLik = log-likelihood. The model with the lowest scoring AICc per barrier was selected as the best model.

| Model | Intercept | Cross | Ecology | Cross:Ecology | df | logLik | AICc | delta | weight |
| --- | --- | --- | --- | --- | --- | --- | --- | --- | --- |
| *Premating I – Mechanical barrier (Binomial distribution)* |  |  |  |  |  |  |  |  |  |
| 8 | 18.57 | + | + | + | 10 | -79.845 | 181.1 | 0 | 0.566 |
| 4 | 3.991 | + | + |  | 6 | -84.545 | 181.6 | 0.53 | 0.434 |
| 3 | 2.262 |  | + |  | 2 | -96.419 | 196.9 | 15.84 | 0 |
| 2 | 2.015 | + |  |  | 5 | -96.243 | 202.9 | 21.78 | 0 |
| 1 | 0.7376 |  |  |  | 1 | -107.015 | 216.1 | 34.98 | 0 |
| *Premating II – Mechanical-tactile barrier (Binomial distribution)* |  |  |  |  |  |  |  |  |  |
| 3 | 3.135 |  | + |  | 2 | -43.943 | 92 | 0 | 0.514 |
| 4 | 4.161 | + | + |  | 6 | -40.18 | 93.1 | 1.14 | 0.29 |
| 8 | 1.792 | + | + | + | 10 | -36.036 | 94.2 | 2.19 | 0.171 |
| 1 | 1.752 |  |  |  | 1 | -48.176 | 98.4 | 6.39 | 0.021 |
| 2 | 2.197 | + |  |  | 5 | -45.63 | 101.8 | 9.82 | 0.004 |
| *Postmating I – Oviposition (Binomial distribution)* |  |  |  |  |  |  |  |  |  |
| 3 | 1.431 |  | + |  | 2 | -48.435 | 101 | 0 | 0.61 |
| 1 | 1.867 |  |  |  | 1 | -49.994 | 102 | 1.05 | 0.36 |
| 4 | 1.605 | + | + |  | 6 | -47.796 | 108.3 | 7.32 | 0.016 |
| 2 | 2.169 | + |  |  | 5 | -49.078 | 108.7 | 7.69 | 0.013 |
| 8 | 2.303 | + | + | + | 10 | -45.653 | 113.2 | 12.23 | 0.001 |
| *Postmating II – Fecundity (Poisson distribution)* |  |  |  |  |  |  |  |  |  |
| 8 | 4.25 | + | + | + | 10 | -3892.949 | 7808 | 0 | 1 |
| 4 | 3.589 | + | + |  | 6 | -4453.062 | 8918.9 | 1110.89 | 0 |
| 3 | 3.3 |  | + |  | 2 | -4656.369 | 9316.8 | 1508.83 | 0 |
| 2 | 4.63 | + |  |  | 5 | -5508.288 | 11027.1 | 3219.11 | 0 |
| 1 | 4.278 |  |  |  | 1 | -5888.144 | 11778.3 | 3970.31 | 0 |
| *Postmating III – Fertility (Binomial distribution)* |  |  |  |  |  |  |  |  |  |
| 8 | -2.372 | + | + | + | 10 | -2140.936 | 4304.7 | 0 | 1 |
| 4 | -2.211 | + | + |  | 6 | -2582.392 | 5177.8 | 873.14 | 0 |
| 3 | -1.501 |  | + |  | 2 | -3113.13 | 6230.4 | 1925.74 | 0 |
| 2 | 0.1178 | + |  |  | 5 | -4321.67 | 8654.1 | 4349.4 | 0 |
| 1 | 0.2091 |  |  |  | 1 | -4392.622 | 8787.3 | 4482.63 | 0 |

**Table S7.** *Post hoc* GLM modeling for reproductive isolation as a function of types of crosses (RI ∼ Cross) per postzygotic reproductive barrier in *Ischnura graellsii* and *I. elegans*. Mechanical-tactile and oviposition barriers were excluded as crosses were not significant parameters in their GLM modeling (Table S6). GLMs were modeled using each cross direction as model intercept to allow pairwise comparisons between types of crosses. S.E. = Standard error; * = Significant p value for differences between a cross and the model intercept (p<0.05/10).

| Barrier | Cross | Estimate | S.E. | z value | p | p<0.05/10 |
| --- | --- | --- | --- | --- | --- | --- |
| Mechanical | **Intercept: hybrid♂×hybrid♀** |  |  |  |  |  |
|  | **Intercept** | 0.4855 | 0.2594 | 1.871 | 0.0613 | NA |
|  | elegans♂×hybrid♀ | 1.5294 | 0.5921 | 2.583 | 0.0098 |  |
|  | graellsii♂×hybrid♀ | -0.7087 | 0.7192 | -0.985 | 0.3245 |  |
|  | hybrid♂×elegans♀ | -0.5427 | 0.4262 | -1.273 | 0.203 |  |
|  | hybrid♂×graellsii♀ | 1.3471 | 0.5978 | 2.254 | 0.0242 |  |
|  | **Intercept: elegans♂×hybrid♀** |  |  |  |  |  |
|  | **Intercept** | 2.0149 | 0.5323 | 3.785 | 0.000153 | NA |
|  | hybrid♂×hybrid♀ | -1.5294 | 0.5921 | -2.583 | 0.0098 |  |
|  | graellsii♂×hybrid♀ | -2.238 | 0.8563 | -2.613 | 0.008962 |  |
|  | hybrid♂×elegans♀ | -2.0721 | 0.6306 | -3.286 | 0.001017 | * |
|  | hybrid♂×graellsii♀ | -0.1823 | 0.7572 | -0.241 | 0.809719 |  |
|  | **Intercept: hybrid♂×graellsii♀** |  |  |  |  |  |
|  | **Intercept** | 1.8326 | 0.5385 | 3.403 | 0.000666 | NA |
|  | elegans♂×hybrid♀ | 0.1823 | 0.7572 | 0.241 | 0.809719 |  |
|  | hybrid♂×hybrid♀ | -1.3471 | 0.5978 | -2.254 | 0.024223 |  |
|  | graellsii♂×hybrid♀ | -2.0557 | 0.8602 | -2.39 | 0.016861 |  |
|  | hybrid♂×elegans♀ | -1.8897 | 0.6359 | -2.972 | 0.002961 | * |
|  | **Intercept: hybrid♂×elegans♀** |  |  |  |  |  |
|  | **Intercept** | -0.05716 | 0.3382 | -0.169 | 0.86579 | NA |
|  | hybrid♂×graellsii♀ | 1.88974 | 0.63591 | 2.972 | 0.00296 | * |
|  | elegans♂×hybrid♀ | 2.07206 | 0.63064 | 3.286 | 0.00102 | * |
|  | hybrid♂×hybrid♀ | 0.54267 | 0.42625 | 1.273 | 0.20297 |  |
|  | graellsii♂×hybrid♀ | -0.16599 | 0.75125 | -0.221 | 0.82514 |  |
|  | **Intercept: graellsii♂×hybrid♀** |  |  |  |  |  |
|  | **Intercept** | -0.2231 | 0.6708 | -0.333 | 0.7394 | NA |
|  | hybrid♂×elegans♀ | 0.166 | 0.7513 | 0.221 | 0.82514 |  |
|  | hybrid♂×graellsii♀ | 2.0557 | 0.8602 | 2.39 | 0.01686 |  |
|  | elegans♂×hybrid♀ | 2.238 | 0.8563 | 2.613 | 0.00896 |  |
|  | hybrid♂×hybrid♀ | 0.7087 | 0.7192 | 0.985 | 0.32449 |  |
| Fecundity | **Intercept: elegans♂×hybrid♀** |  |  |  |  |  |
|  | **Intercept** | 4.63026 | 0.01623 | 285.203 | <2E-16 | NA |
|  | graellsii♂×hybrid♀ | 0.04257 | 0.07026 | 0.606 | 0.545 |  |
|  | hybrid♂×hybrid♀ | -0.62293 | 0.02463 | -25.292 | <2E-16 | * |
|  | hybrid♂×elegans♀ | -0.70236 | 0.04724 | -14.866 | <2E-16 | * |
|  | hybrid♂×graellsii♀ | -0.43363 | 0.0377 | -11.503 | <2E-16 | * |
|  | **Intercept: hybrid♂×hybrid♀** |  |  |  |  |  |
|  | **Intercept** | 4.00733 | 0.01852 | 216.359 | <2E-16 | NA |
|  | elegans♂×hybrid♀ | 0.62293 | 0.02463 | 25.292 | <2E-16 | * |
|  | graellsii♂×hybrid♀ | 0.6655 | 0.07082 | 9.397 | <2E-16 | * |
|  | hybrid♂×elegans♀ | -0.07944 | 0.04808 | -1.652 | 0.0985 | . |
|  | hybrid♂×graellsii♀ | 0.18929 | 0.03874 | 4.887 | 1.03E-06 | * |
|  | **Intercept: hybrid♂×graellsii♀** |  |  |  |  |  |
|  | **Intercept** | 4.19662 | 0.03402 | 123.355 | <2E-16 | NA |
|  | hybrid♂×hybrid♀ | -0.18929 | 0.03874 | -4.887 | 1.03E-06 | * |
|  | elegans♂×hybrid♀ | 0.43363 | 0.0377 | 11.503 | <2E-16 | * |
|  | graellsii♂×hybrid♀ | 0.47621 | 0.07636 | 6.237 | 4.47E-10 | * |
|  | hybrid♂×elegans♀ | -0.26873 | 0.05591 | -4.806 | 1.54E-06 | * |
|  | **Intercept: hybrid♂×elegans♀** |  |  |  |  |  |
|  | **Intercept** | 3.9279 | 0.04437 | 88.53 | <2E-16 | NA |
|  | hybrid♂×graellsii♀ | 0.26873 | 0.05591 | 4.806 | 1.54E-06 | * |
|  | hybrid♂×hybrid♀ | 0.07944 | 0.04808 | 1.652 | 0.0985 |  |
|  | elegans♂×hybrid♀ | 0.70236 | 0.04724 | 14.866 | <2E-16 | * |
|  | graellsii♂×hybrid♀ | 0.74493 | 0.08149 | 9.141 | <2E-16 | * |
|  | **Intercept: graellsii♂×hybrid♀** |  |  |  |  |  |
|  | **Intercept** | 4.67283 | 0.06836 | 68.358 | <2E-16 | NA |
|  | hybrid♂×elegans♀ | -0.74493 | 0.08149 | -9.141 | <2E-16 | * |
|  | hybrid♂×graellsii♀ | -0.47621 | 0.07636 | -6.237 | 4.47E-10 | * |
|  | hybrid♂×hybrid♀ | -0.6655 | 0.07082 | -9.397 | <2E-16 | * |
|  | elegans♂×hybrid♀ | -0.04257 | 0.07026 | -0.606 | 0.545 |  |
| Fertility | **Intercept: hybrid♂×hybrid♀** |  |  |  |  |  |
|  | **Intercept** | 0.16714 | 0.0267 | 6.259 | 3.87E-10 | NA |
|  | elegans♂×hybrid♀ | -0.04939 | 0.03491 | -1.415 | 0.157 |  |
|  | graellsii♂×hybrid♀ | 0.86205 | 0.09351 | 9.219 | <2E-16 | * |
|  | hybrid♂×elegans♀ | 0.32044 | 0.05865 | 5.464 | 4.66E-08 | * |
|  | hybrid♂×graellsii♀ | 0.08488 | 0.05952 | 1.426 | 0.154 |  |
|  | **Intercept: elegans♂×hybrid♀** |  |  |  |  |  |
|  | **Intercept** | 0.11775 | 0.02248 | 5.238 | 1.63E-07 | NA |
|  | hybrid♂×hybrid♀ | 0.04939 | 0.03491 | 1.415 | 0.1571 |  |
|  | graellsii♂×hybrid♀ | 0.91144 | 0.09239 | 9.865 | <2E-16 | * |
|  | hybrid♂×elegans♀ | 0.36983 | 0.05685 | 6.505 | 7.75E-11 | * |
|  | hybrid♂×graellsii♀ | 0.13427 | 0.05775 | 2.325 | 0.0201 |  |
|  | **Intercept: hybrid♂×elegans♀** |  |  |  |  |  |
|  | **Intercept** | 0.48758 | 0.05222 | 9.338 | <2E-16 | NA |
|  | elegans♂×hybrid♀ | -0.36983 | 0.05685 | -6.505 | 7.75E-11 | * |
|  | hybrid♂×hybrid♀ | -0.32044 | 0.05865 | -5.464 | 4.66E-08 | * |
|  | graellsii♂×hybrid♀ | 0.54161 | 0.10372 | 5.222 | 1.77E-07 | * |
|  | hybrid♂×graellsii♀ | -0.23556 | 0.07454 | -3.16 | 0.00158 | * |
|  | **Intercept: graellsii♂×hybrid♀** |  |  |  |  |  |
|  | **Intercept** | 1.0292 | 0.08962 | 11.484 | <2E-16 | NA |
|  | hybrid♂×elegans♀ | -0.54161 | 0.10372 | -5.222 | 1.77E-07 | * |
|  | elegans♂×hybrid♀ | -0.91144 | 0.09239 | -9.865 | <2E-16 | * |
|  | hybrid♂×hybrid♀ | -0.86205 | 0.09351 | -9.219 | <2E-16 | * |
|  | hybrid♂×graellsii♀ | -0.77717 | 0.10422 | -7.457 | 8.84E-14 | * |
|  | **Intercept: hybrid♂×graellsii♀** |  |  |  |  |  |
|  | **Intercept** | 0.25202 | 0.0532 | 4.737 | 2.16E-06 | NA |
|  | graellsii♂×hybrid♀ | 0.77717 | 0.10422 | 7.457 | 8.84E-14 | * |
|  | hybrid♂×elegans♀ | 0.23556 | 0.07454 | 3.16 | 0.00158 | * |
|  | elegans♂×hybrid♀ | -0.13427 | 0.05775 | -2.325 | 0.02008 |  |
|  | hybrid♂×hybrid♀ | -0.08488 | 0.05952 | -1.426 | 0.15388 |  |

**Table S8.** *Post hoc* GLM modeling for reproductive isolation as a function of the interaction ecology and types of crosses (RI ∼ Ecology:Cross) per postzygotic reproductive barrier in *Ischnura graellsii* and *I. elegans*. Mechanical-tactile and oviposition barriers were excluded as ecology and cross interaction were not significant parameters in its GLM modeling (Table S6). Although each cross in each ecology was compared with each other combination, here we report only results for differences between ecologies within each type of cross. * = Significant p value for differences between the allopatric and sympatric ecology (p<0.05/5).

| Barrier | Cross | Estimate | Std. Error | z value | p | p<0.05/5 |
| --- | --- | --- | --- | --- | --- | --- |
| **Mechanical** | hybrid♂×hybrid♀ | 18.6996 | 1537.4007 | 0.012 | 0.9903 |  |
|  | elegans♂×hybrid♀ | 16.81687 | 2465.32572 | 0.007 | 0.994557 |  |
|  | hybrid♂×graellsii♀ | -0.08004 | 1.07715 | -0.074 | 0.94076 |  |
|  | hybrid♂×elegans♀ | 1.7272 | 0.7976 | 2.166 | 0.030341 |  |
|  | graellsii♂×hybrid♀ | 19.4824 | 4612.2021 | 0.004 | 0.99663 |  |
| **Fecundity** | elegans♂×hybrid♀ | -0.49114 | 0.04183 | -11.741 | <2e-16 | * |
|  | hybrid♂×hybrid♀ | -2.13006 | 0.06487 | -32.835 | <2e-16 | * |
|  | hybrid♂×graellsii♀ | -2.40932 | 0.0708 | -34.032 | <2e-16 | * |
|  | hybrid♂×elegans♀ | -1.49568 | 0.11473 | -13.037 | <2e-16 | * |
|  | graellsii♂×hybrid♀ | 1.5012 | 0.1771 | 8.478 | <2e-16 | * |
| **Fertility** | hybrid♂×hybrid♀ | -5.67083 | 0.57957 | -9.785 | <2e-16 | * |
|  | elegans♂×hybrid♀ | -3.16696 | 0.0831 | -38.109 | <2e-16 | * |
|  | hybrid♂×elegans♀ | -5.62044 | 0.5841 | -9.622 | <2e-16 | * |
|  | graellsii♂×hybrid♀ | 3.7232 | 0.2986 | 12.467 | <2e-16 | * |
|  | hybrid♂×graellsii♀ | -4.2015 | 1.01183 | -4.152 | 3.29E-05 | * |

**Table S9.** GLM models comparison per prezygotic reproductive barrier comparing the two reciprocal heterospecific crosses (crosses between *Ischnura elegans* males and *I. graellsii* females vs crosses between *I. graellsii* males and *I. elegans* females). Models are sorted per reproductive barrier by increasing values of the AICc. “+” sign on the cross parameter show the inclusion of heterospecific crosses as a parameter explaining RI. df = degrees freedom; logLik = log-likelihood. If the model including the cross parameter had the lowest AICc value, we concluded significant prezygotic asymmetries were present on that barrier.

| Barrier | Model | Intercept | Cross | df | logLik | AICc | delta | weight |
| --- | --- | --- | --- | --- | --- | --- | --- | --- |
| **All allopatric data** |  |  |  |  |  |  |  |  |
| **Mechanical** | 1 | 1.58 |  | 1 | -37.478 | 77 | 0 | 0.741 |
|  | 2 | 1.576 | + | 2 | -37.477 | 79.1 | 2.1 | 0.259 |
| **Mechanical-Tactile** | 1 | 1.442 |  | 1 | -33.179 | 68.4 | 0 | 0.669 |
|  | 2 | 1.344 | + | 2 | -32.82 | 69.8 | 1.41 | 0.331 |
| **Oviposition** | 2 | 0.5465 | + | 2 | -19.715 | 43.8 | 0 | 0.789 |
|  | 1 | 0.821 |  | 1 | -22.158 | 46.4 | 2.64 | 0.211 |
| **Fecundity** | 2 | 3.062 | + | 2 | -462.131 | 928.8 | 0 | 1 |
|  | 1 | 4.289 |  | 1 | -1629.01 | 3260.2 | 2331.39 | 0 |
| **Fertility** | 2 | -5.061 | + | 2 | -875.601 | 1755.7 | 0 | 1 |
|  | 1 | 0.5421 |  | 1 | -1575.386 | 3152.9 | 1397.2 | 0 |
| **All sympatric data** |  |  |  |  |  |  |  |  |
| **Mechanical** | 2 | 1.328 | + | 2 | -71.849 | 147.8 | 0 | 1 |
|  | 1 | 0.3455 |  | 1 | -103.124 | 208.3 | 60.5 | 0 |
| **Mechanical-Tactile** | 1 | 1.175 |  | 1 | -48.627 | 99.3 | 0 | 0.709 |
|  | 2 | 1.214 | + | 2 | -48.47 | 101.1 | 1.78 | 0.291 |
| **Oviposition** | 1 | 2.411 |  | 1 | -24.181 | 50.4 | 0 | 0.687 |
|  | 2 | 2.372 | + | 2 | -23.918 | 52 | 1.57 | 0.313 |
| **Fecundity** | 2 | 4.49 | + | 2 | -3243.226 | 6490.6 | 0 | 1 |
|  | 1 | 4.509 |  | 1 | -3271.21 | 6544.5 | 53.86 | 0 |
| **Fertility** | 1 | 1.119 |  | 1 | -2830.357 | 5662.8 | 0 | 0.711 |
|  | 2 | 1.121 | + | 2 | -2830.204 | 5664.6 | 1.8 | 0.289 |
| **Allopatry: Cachadas×Belgium vs Belgium×Cachadas** |  |  |  |  |  |  |  |  |
| **Mechanical** | 1 | 1.459 |  | 1 | -25.668 | 53.4 | 0 | 0.746 |
|  | 2 | 1.447 | + | 2 | -25.666 | 55.6 | 2.16 | 0.254 |
| **Mechanical-Tactile** | 1 | 1.194 |  | 1 | -23.321 | 48.7 | 0 | 0.639 |
|  | 2 | 1.022 | + | 2 | -22.789 | 49.9 | 1.14 | 0.361 |
| **Oviposition** | 1 | 1.466 |  | 1 | -7.721 | 17.7 | 0 | 0.511 |
|  | 2 | 0.9808 | + | 2 | -6.445 | 17.8 | 0.09 | 0.489 |
| **Fecundity** | 2 | 2.931 | + | 2 | -82.398 | 170 | 0 | 1 |
|  | 1 | 4.679 |  | 1 | -872.914 | 1748.2 | 1578.2 | 0 |
| **Fertility** | 2 | -5.056 | + | 2 | -374.343 | 753.9 | 0 | 1 |
|  | 1 | 1.122 |  | 1 | -851.08 | 1704.5 | 950.64 | 0 |
| **Sympatry: Corrubedo×Louro vs Louro×Corrubedo** |  |  |  |  |  |  |  |  |
| **Mechanical** | 1 | 1.386 |  | 1 | -2.502 | 8.3 | 0 | 0.956 |
|  | 2 | 18.57 | + | 2 | -2.249 | 14.5 | 6.16 | 0.044 |
| **Mechanical-Tactile** | 1 | 23.57 |  | 1 | 0 | 4 | 0 | 0.998 |
|  | 2 | 23.57 | + | 2 | 0 | 16 | 12 | 0.002 |
| **Oviposition** | 1 | 24.57 |  | 1 | 0 | 3.3 | 0 | 0.966 |
|  | 2 | 24.57 | + | 2 | 0 | 10 | 6.67 | 0.034 |
| **Fecundity** | 2 | 4.431 | + | 2 | -124.964 | 259.9 | 0 | 1 |
|  | 1 | 4.736 |  | 1 | -138.639 | 280.6 | 20.68 | 0 |
| **Fertility** | 1 | 1.168 |  | 1 | -47.679 | 98.7 | 0 | 0.615 |
|  | 2 | 1.389 | + | 2 | -44.814 | 99.6 | 0.94 | 0.385 |
| **Sympatry: Lanzada×Louro vs Louro×Lanzada** |  |  |  |  |  |  |  |  |
| **Mechanical** | 2 | 19.57 | + | 2 | -4.157 | 12.7 | 0 | 1 |
|  | 1 | -1.056 |  | 1 | -17.702 | 37.5 | 24.8 | 0 |
| **Mechanical-Tactile** | 1 | 0.5108 |  | 1 | -5.293 | 13.3 | 0 | 0.682 |
|  | 2 | 0.9163 | + | 2 | -4.188 | 14.8 | 1.52 | 0.318 |
| **Sympatry: Cachadas×Laxe vs Laxe×Cachadas** |  |  |  |  |  |  |  |  |
| **Mechanical** | 2 | 1.347 | + | 2 | -38.465 | 81.1 | 0 | 1 |
|  | 1 | 0.55 |  | 1 | -53.85 | 109.8 | 28.67 | 0 |
| **Mechanical-Tactile** | 1 | 1.316 |  | 1 | -26.831 | 55.7 | 0 | 0.659 |
|  | 2 | 1.386 | + | 2 | -26.406 | 57.1 | 1.31 | 0.341 |

## Text S1. Supplementary Methods: Estimation of the absolute strength of the reproductive barriers between *Ischnura graellsii* and *I. elegans*

Mechanical and mechanical-tactile barriers measure the incompatibility between the males’ caudal appendages and the females’ prothorax, the failure in the stimulation by the male to the female in the tandem position, and the incompatibility between the males’ and females’ genital structures.

We estimated the first premating barrier (Premating I – mechanical barrier) as:

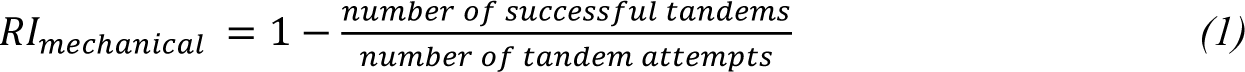

in which we defined a tandem attempt when a male flew towards a female and curled his abdomen to try to grab her with his caudal appendages. If a male tried several times to grab a specific female (either on the same day or on multiple experimental days), we only counted this interaction as a single tandem attempt. By doing this, the sample size shows the number of male-female pairs in which at least one tandem was attempted. If, in at least one of these tandem attempts, the male correctly grabbed the female and the couple remained together in tandem position (Fig. 2A), a successful tandem was recorded; i.e., multiple tandems made by the same male-female pair were recorded as a single successful event.

We estimated the second premating barrier (premating II – mechanical-tactile barrier) as:

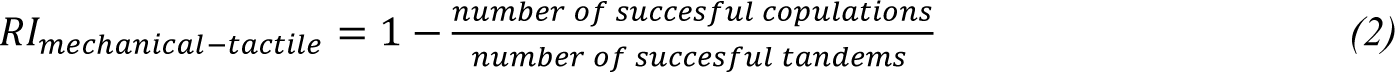

in which a successful copulation was recorded if a female in the tandem position bent her abdomen and placed it in contact with male genitalia (Fig. 2B). Male-female pairs that formed the mating position, reverted to a tandem (or free themselves completely) and then formed a second mating position were considered as a single successful copulation event; i.e., the number of successful copulations shows the number of male-female pairs that achieved at least one successful mating position. Pairs that achieved this position were carefully observed and isolated on individual jars. To avoid additional copulations of females, we considered as “mated” each female that achieved this position without regarding the length pairs remained in copula or the number of copulas they formed.

The first two postmating barriers measure: 1) postmating I – oviposition, how the heterospecific ejaculate fails to stimulate female oviposition (number of females that laid eggs; Fig. 2C); and 2) postmating II – fecundity, how the heterospecific ejaculate reduces the frequency of oviposition [number of laid eggs by female (Coyne and Orr 2004); Fig. 2D]. The third postmating barrier (postmating III – fertility) measures several processes: poor transfer or storage sperm, unviability of gametes in the foreign reproductive tract, poor movement or cross-attraction, or failure of fertilization when gametes contact each other [(Coyne and Orr 2004); Fig. 2D]. We measured postmating barriers II and III using the first three clutches.

We estimated the first postmating barrier (postmating I – oviposition) as:

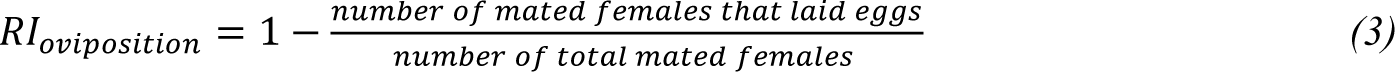

For the second postmating barrier (postmating II – fecundity), first for each mated female we measured the mean number of eggs they laid per clutch in the first three clutches. We excluded females that did not lay eggs (oviposition barrier) and, if females survived less than the first three oviposition days, we averaged the number of eggs they laid on the days they lived (i.e. in one or two clutches). We refer to this value as the eggs per clutch index. Then, we averaged this number for all females of the same type of cross per population (i.e. the population column in Table S2), and used a mathematical correction to estimate a RI strength value in a range from 0 to 1, as we had with the other barriers:

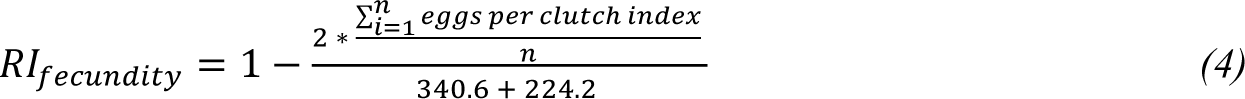

in which n refers to the number of laying females for each type of cross per cross of populations. The 340.6 and 224.4 values on the denominator of equation 4 refers to the maximum average eggs per clutch index seen in allopatric conspecific crosses. While the former refers to the average fecundity of *I. elegans* allopatric crosses in Arles, the latter refers to the average fecundity of *I. graellsii* allopatric crosses in Alba (Table S2). By using the same conspecific values in all fecundity RI estimations, our results reflected only the changes in heterospecific eggs per clutch indices. When the average eggs per clutch index of a population cross was higher than the average of the conspecific corrections, and thus a negative value of RI was estimated, we rounded up the RI value to zero.

Finally, we estimated the third postmating barrier (postmating III – fertility) as:

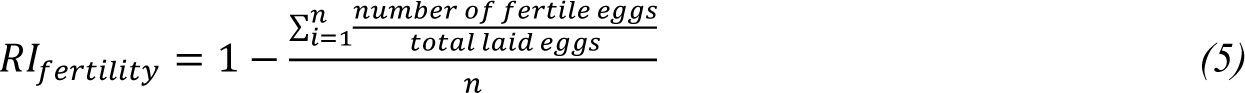

in which we identified fertile eggs as those having evidence of hatching or of a developing embryo, and n refers to the number of laying females per type of cross per population.

## Text S2. Supplementary Methods: Testing reinforcement predictions

### Strengthening of prezygotic barriers

Since Dobzhansky’s earliest work in reinforcement (Dobzhansky 1937, 1940; Dobzhansky and Koller 1938), the classical test of reinforcement is done by contrasting the strength of prezygotic isolation in sympatry *versus* in allopatry (Coyne and Orr 1989, 1997, 2004). We expected stronger total prezygotic isolation in sympatry than in allopatry in *Ischnura graellsii* and *I. elegans,* and stronger absolute isolation in sympatry than in allopatry in the reproductive barrier under reinforcement.

### Rarer female effect

Since usually females pay higher fitness costs of hybridization than males (Coyne and Orr 2004), and females of the rarer species have a higher chance of being involved in an heterospecific mating than females of the more common species, reinforcement is expected to strengthen prezygotic isolation faster in the cross direction involving females of the rarer species (Yukilevich 2012). Since *I. elegans* is the invader species in Spain, this species is less frequent in the sympatry zone than *I. graellsii* (Sánchez-Guillén et al. 2011). Thus, for this prediction we expected stronger prezygotic isolation in crosses between *I. graellsii* males and *I. elegans* females than the reciprocal cross in sympatry but not in allopatry. Additionally, since in a local-scale species frequencies vary between localities (Sánchez-Guillén et al. 2023), we expected stronger prezygotic isolation in sympatric crosses between Corrubedo (*I. graellsii* males) and Louro (*I. elegans* females), and Lanzada (*I. graellsii* males) and Louro (*I. elegans* females) than in crosses between Cachadas (*I. graellsii* males) and Laxe (*I. elegans* females). The reason for this is that while historically Louro has been an *I. graellsii*-dominant locality, Laxe has been an *I. elegans-*dominant locality (Table S1).

### Concordant prezygotic and postzygotic isolation asymmetries

Unidirectionally inherited Bateson-Dobzhansky-Müller (BDM) incompatibilities associated with sex or cytoplasmic chromosomes cause postzygotic isolation to be asymmetric between reciprocal crosses (Turelli and Moyle 2007). Since under reinforcement, hybridization costs (postzygotic barriers) and prezygotic isolation are expected to be positively correlated (Ortiz-Barrientos et al. 2009), concordant prezygotic and postzygotic isolation asymmetries between reciprocal crosses are expected in sympatry but not in allopatry (Yukilevich 2012). For this prediction we expected that cross directions with stronger postzygotic isolation (highest hybridization costs) have also stronger total prezygotic isolation in sympatry but not in allopatry.

### Greater premating asymmetries and weaker postzygotic isolation in sympatry than in allopatry

Species pairs with asymmetric postzygotic isolation in sympatry are expected to have higher premating asymmetries in sympatry than in allopatry under reinforcement (Turelli et al. 2014). Additionally, since gene flow operates only in sympatry, crosses from sympatry should also have weaker postzygotic isolation owing to the suppression of BDM incompatibilities (Turelli et al. 2014). For this prediction we expected statistically significant differences in premating isolation (mechanical or mechanical-tactile barriers) between reciprocal heterospecific crosses in sympatry but not in allopatry, and weaker postzygotic isolation in sympatry than in allopatry. To detect significant asymmetries in prezygotic isolation we compared the null model GLM to one that included RI as a function of the two heterospecific crosses per prezygotic barrier. If the model including the crosses variable scored a lower AICc value, then we considered this statistical support for difference between heterospecific crosses. These tests were made: i) pooling all sympatric and allopatric data, and ii) in specific population crosses in which we measured the two reciprocal directions.

